# Uncovering hypergraphs of cell-cell interaction from single cell RNA-sequencing data

**DOI:** 10.1101/566182

**Authors:** Koki Tsuyuzaki, Manabu Ishii, Itoshi Nikaido

## Abstract

Complex biological systems can be described as a multitude of cell-cell interactions (CCIs). Recent single-cell RNA-sequencing technologies have enabled the detection of CCIs and related ligand-receptor (L-R) gene expression simultaneously. However, previous data analysis methods have focused on only one-to-one CCIs between two cell types. To also detect many-to-many CCIs, we propose `scTensor`, a novel method for extracting representative triadic relationships (hypergraphs), which include (i) ligand-expression, (ii) receptor-expression, and (iii) L-R pairs. When applied to simulated and empirical datasets, `scTensor` was able to detect some hypergraphs including paracrine/autocrine CCI patterns, which cannot be detected by previous methods.

## Background

Complex biological systems such as tissue homeostasis [1, 2], neurotransmission [3, 4], immune response [5], ontogenesis [6], and stem cells niche [7, 8] are composed by cell-cell interaction (CCI). Many molecular biology studies have been decomposed the system into constituent parts (e.g., genes, proteins, and metabolites) to clarify their functions. Nevertheless, more sophisticated methodologies are still required, because CCI is the difference between the whole system and sum of their parts.

Previous studies have investigated CCIs using technologies such as fluorescence microscopy [9–13], microdevise-based methods such as microwells, micropatterns, single-cell traps, droplet microfluidics, and micropillars [14–22], and transcriptome-based method [23–53]. In particular, the recent development of single-cell RNA-sequencing (scRNA-seq) technologies has enabled the detection of exhaustive cell-type-level CCIs based on ligand and receptor (L-R) coexpression.

To assess the coexpression of known L-R genes, circle plots [24, 33, 36, 44, 45, 50], bigraph/Sankey diagrams [29, 30, 34, 49, 52, 53], network diagrams [26–28, 31, 37, 40, 41, 44, 46, 54] and heatmaps [30, 32, 34, 39, 46, 47, 49] are often drawn. Some studies have also introduced more systematical approaches for quantifying the degree of CCIs based on the L-R coexpression, such as the number of coexpressed L-R pairs [23, 24, 26, 27], Spearman correlation coefficients between L-R expression profiles [26, 31], original interaction scores between L-R coexpression [39], or hypothetical test based on random cell-type label permutation [29, 32, 37, 55].

All the approaches described above implicitly suppose that CCIs are the one-to-one relationships between two cell types and that the corresponding L-R coexpression is observed as the cell-type-specific manner. In real empirical data, however, the situation can be more complex; CCIs often exhibit many-to-many relationships involving many cell types, and an particular L-R pair can also function across multiple cell-type pairs. Therefore, in this work, we propose `scTensor`, which is a novel method based on a tensor decomposition algorithm. Our method regards CCIs as hypergraphs and extracts some representative triadic relationship from the data tensor, which includes (i) ligand-expressing cell types, (ii) receptor-expressing cell types, and (iii) L-R pairs.

### CCI as hypergraph (*CaH*) and CCI-tensor

The simplest CCI representation is perhaps a directed graph, where each node represents a cell type and each edge represents the coexpression of all L-R pairs (Figure 1a, left). The direction of the edge is set as ligand *→* receptor. Such a data structure can also be described as an asymmetric adjacency matrix, in which each row and column represents a ligand and receptor, respectively. If some combinations of cell types are regarded as interacting, corresponding elements of the matrix are filled with 1 and otherwise 0. If the degree of CCI is not a binary relationship, weighted graphs and corresponding weighted adjacent matrices may also be used. The previous analytical methods are categorized within this approach [23, 24, 26–34, 36, 37, 39–41, 44–47, 49, 50, 52, 53, 55].

**Figure 1.**
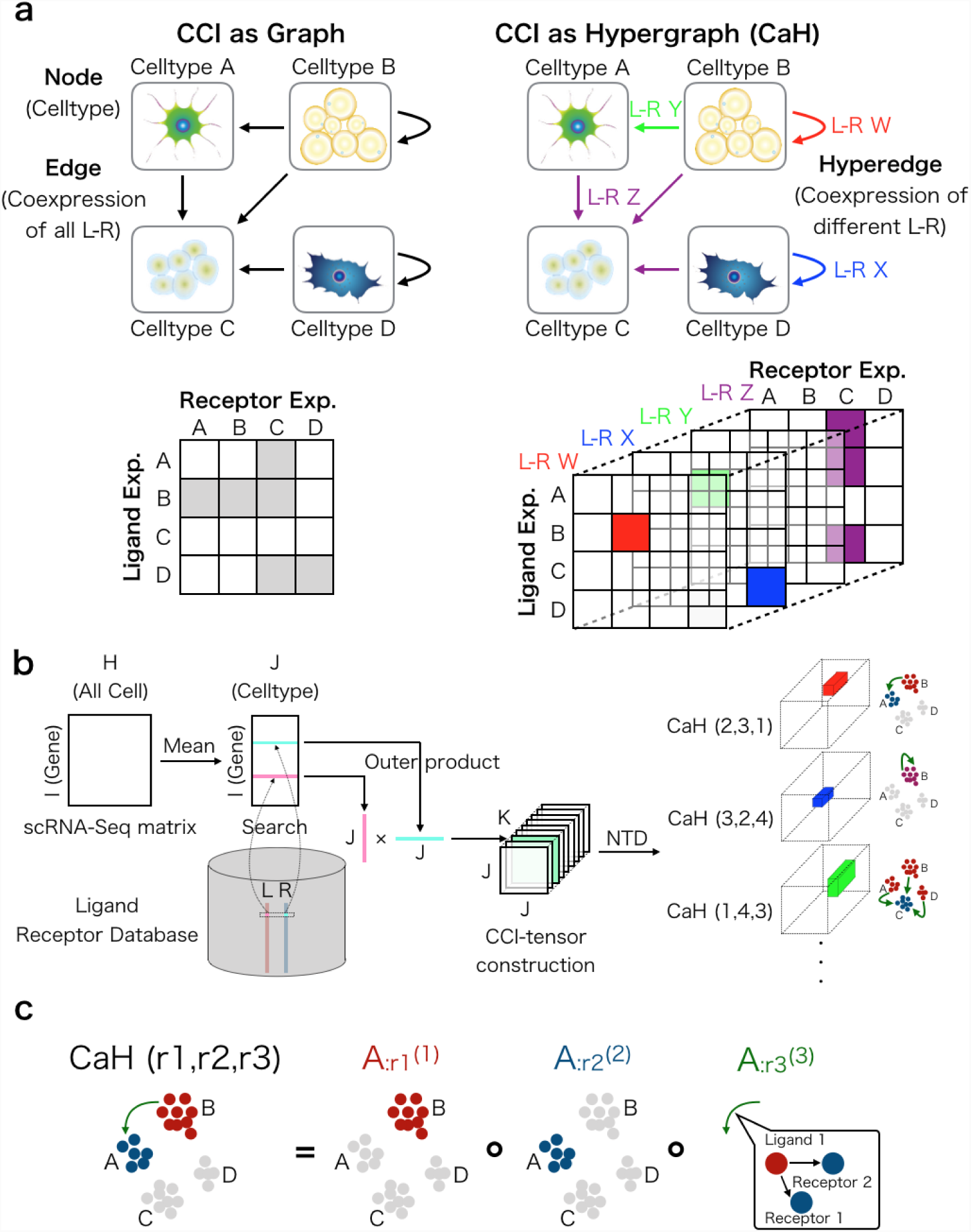
CCI as a hypergraph. (a) Previous studies have regarded CCIs as graphs, and the corresponding data structure is expressed as an adjacency matrix (left). In this work, CCIs are regarded as hypergraphs, and the corresponding data structure is a tensor. (b) The CCI-tensor is generated by users’ scRNA-Seq matrices, cell-type labels, and L-R databases. NTD is used to extract *CaH*s from the CCI-tensor. (c) Each *CaH*(r1,r2,r3) is equal to the outer product of three vectors. 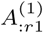 represents the ligand expression pattern, 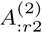 represents the receptor expression pattern, and 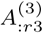 represents the related L-R pairs pattern.

**Figure 2.**
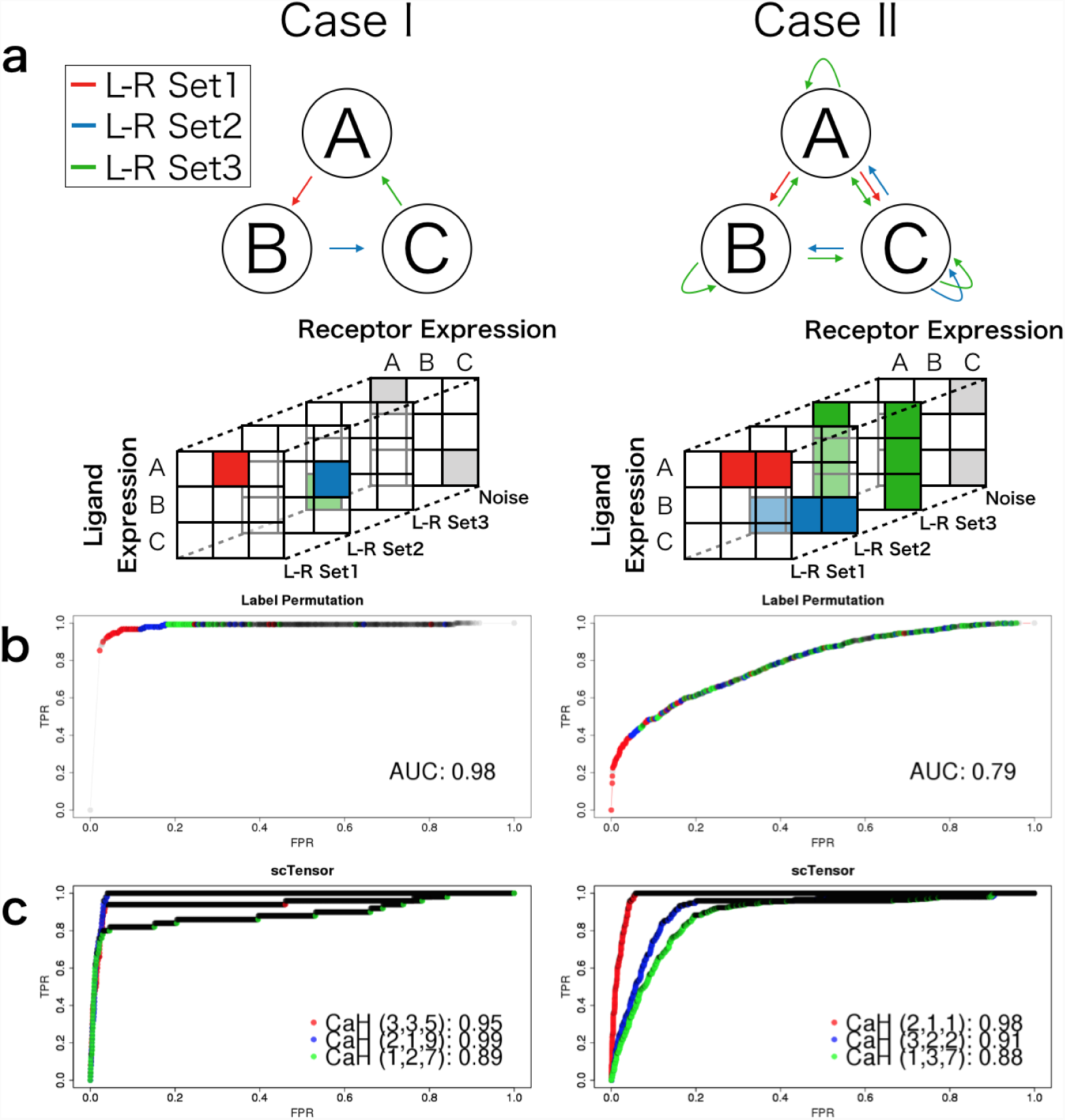
Comparison between the label permutation method and scTensor using simulated datasets. (a) Two datasets simulating two cases were constructed. In the first case, all CCIs consisted of one-to-one relationships between two cell types. In the second case, CCIs consisted of many-to-many relationships involving many cell types. (b) The ROC curve for the label permutation method. (c) The ROC curves of the *CaH*s detected by `scTensor`.

In contrast, this work describes CCIs as directed hypergraphs (CCI as hypergraph; *CaH*), where each node is a cell type but the edges are distinguished from each other by the different related L-R pair sets (Figure 1a, right). Such a context-aware edge is called a hyperedge and is described as multiple different adjacency matrices and the set of matrices is called a higher-order matrix or tensor. In contrast with the simple adjacency matrix, tensor contains considerable high-resolution information owing to its higher-order.

### Prediction of many-to-many CCIs using tensor decomposition

Tensor data are constructed through the following steps (Figure 1b). Here, a scRNA-seq matrix and the cellular label specifying cell types are supposed to be provided by users. Firstly, Freeman-Tukey transformation [56] 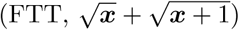, which is a variance-stabilizing transformation, is performed to the data matrix. Next, the matrix is converted to a cell-type-level average matrix according to the label. Combined with a L-R database, two corresponding row-vectors of an L-R pair are extracted from the matrix. We originally developed the databases for 12 organisms (for more details, see Additional Files 1, 2, Table 2 and 3). The outer product (direct product) of the two vectors is then calculated, and a matrix is generated. The matrix can be considered as the similarity matrix of all possible cell-type combinations using the L-R pair. Finally, for each L-R pair, the matrix is calculated, and a tensor is generated as the combined matrices. In this work, this is called the CCI-tensor.

**Table 1.** Empirical datasets

| Name | Organisms | ID | # Gene | # Cell | # Cell type | Unit |
| --- | --- | --- | --- | --- | --- | --- |
| Germline.Female [25] | *Homo sapiens* | GSE86146 | 2717 | 992 | 8 | TPM |
| Germline.Male [25] | *Homo sapiens* | GSE86146 | 2790 | 852 | 7 | TPM |
| Melanoma [59] | *Homo sapiens* | GSE72056 | 6603 | 2250 | 6 | TPM |
| NonMyocyte [27] | *Mus musculus* | E-MTAB-6173 | 2587 | 10519 | 12 | UMI count |
| Habenular.Larva [60] | *Danio rerio* | GSE109158 | 15206 | 4233 | 16 | UMI count |

**Table 2.** Summary of `LRBase.XXX.eg.db` for 12 organisms (1/2)

| XXX | Organisms | SWISSPROT<br>(Secreted / Membrane) | TrEMBL<br>(Secreted / Membrane) | STRING<br>(PPI) |
| --- | --- | --- | --- | --- |
| Hsa | *Homo sapiens* | 1592 / 2269 | 176 / 334 | 18838 |
| Mmu | *Mus musculus* | 1309 / 1806 | 325 / 1555 | 19715 |
| Ath | *Arabidopsis thaliana* | 1260 / 1001 | 244 / 80 | 24174 |
| Rno | *Rattus norvegicus* | 643 / 983 | 232 / 1229 | 19963 |
| Bta | *Bos taurus* | 517 / 448 | 192 / 390 | 18349 |
| Cel | *Caenorhabditis elegans* | 198 / 247 | 28 / 60 | 13545 |
| Dme | *Drosophila melanogaster* | 249 / 333 | 89 / 148 | 11903 |
| Dre | *Danio rerio* | 119 / 169 | 318 / 376 | 21746 |
| Gga | *Gallus gallus* | 175 / 173 | 185 / 154 | 13084 |
| Pab | *Pongo abelii* | 80 / 134 | 212 / 211 | 16691 |
| Xtr | *Xenopus Silurana tropicalis* | 57 / 83 | 141 / 114 | 15338 |
| Ssc | *Sus scrofa* | 223 / 153 | 202 / 445 | 18683 |

**Table 3.** Summary of `LRBase.XXX.eg.db` for 12 organisms (2/2)

| XXX | # L-R Pairs (SWISSPROT × STRING) | # L-R Pairs (TrEMBL × STRING) |
| --- | --- | --- |
| Hsa | 21882 | 472 |
| Mmu | 16386 | 476 |
| Ath | 8697 | 94 |
| Rno | 5270 | 65 |
| Bta | 2220 | 237 |
| Cel | 106 | 1 |
| Dme | 384 | 9 |
| Dre | 99 | 432 |
| Gga | 140 | 105 |
| Pab | 34 | 184 |
| Xtr | 19 | 107 |
| Ssc | 277 | 130 |

After the construction of a CCI-tensor, we perform non-negative Tucker decomposition (NTD [57, 58]), which is known as a tensor decomposition algorithm. We originally implemented this algorithm and confirmed its convergence within a realistic number of iterations (for more details, see Additional File 3). NTD assumes that the CCI-tensor can be approximated by the summation of some representative *CaH* s. NTD has the three rank parameters (*R*1, *R*2, *and R*3) and a *CaH* is calculated as the outer product of the column vectors of three factor matrices ***A***^(1)^ ∈ ℝ^*J×R*1^, ***A***^(2)^ ∈ ℝ^*J×R*2^, and ***A***^(3)^ ∈ ℝ^*K×R*3^ calculated by NTD (Figure 1c). Each *CaH* - strength is calculated by the core tensor 𝒢 (*r*1, *r*2, *r*3) ∈ ℝ^*R*1*×R*2*×R*3^ of NTD. In this work, each *CaH* is termed 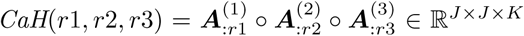, where r1, r2, and r3 are the indexes of the columns of three factor matrices. All *CaH* s are ordered by the size of elements of the core tensor, and the patterns explaining the top 40 % of cumulative core tensor value are reserved as representative *CaH* s. For more details on *CaH*, see the Materials and Methods. The *CaH* s are extracted in a data-driven way without the assumption of one-to-one CCIs. Therefore, it can also detect many-to-many CCIs according to the data complexity.

## Results and discussion

### Evaluation of multiple CCI prediction

#### Accuracy of the detection of CCIs and the related L-R pairs

Here, we demonstrate the efficacy of `scTensor` by using the two simulation datasets (Figure2a). Three different cell types are indicated as ”A”, ”B”, and ”C”. In the case I dataset, all CCIs represent the one-to-one relationships between two cell types. CCIs corresponding to A *→* B, B *→* C, and C *→* A are colored by red, blue, and green, respectively. In contrast, the CCIs in the case II dataset represent many-to-many relationships involving many cell types such as A *→* B/C (red), C *→* A/B/C (blue), and A/B/C *→* B/C (green). To evaluate whether such ground truth combination of cell types and their related L-R pairs are enriched by `scTensor`, receiver operating characteristic (ROC) curves and their corresponding area under the curve (AUC) values were calculated (Figure2b). In the analysis of each L-R set, only the *CaH* s with the maximum AUC value for each ground truth L-R set were regarded as the corresponding *CaH* being accurately detected by `scTensor` (Figure2c, for more details on the simulation datasets, see the Materials and Methods).

Again, note that the *CaH* s detected by `scTensor` are not just CCIs, but sets of CCIs and their related L-R pairs. To the best of our knowledge, the label permutation method implemented in CellPhoneDB [37] is the only previous method that can detect CCIs and their related L-R pairs simultaneously. To demonstrate the efficacy of `scTensor` in terms of the detection of many-to-many CCIs, we also originally implemented this method and compared it with `scTensor` (for more details on the algorithm, see the Materials and Methods). Note that each combination of a CCI and its related L-R pair is separately extracted by NTD of `scTensor`, but under the label permutation method, the combinations are not separated and are just sorted in ascending order of their *P*-values. Combinations with low *P*-values indicate significant triadic relationships.

In the case I dataset, ground truth L-R sets are highly enriched according to the measures of both methods, and the AUC values show that there is no difference in their performance (Figure2b and c, left). On the other hand, in the case II dataset, the label permutation method cannot correctly detect blue or green L-R sets, and the AUC value becomes lower (Figure2b, right). In the `scTensor` analysis, red, blue, and green L-R sets are separately extracted as three *CaH* s (Figure2c, right), and the AUC values are still high. This is because, the label permutation is implicitly hypothesized the CCI as a one-to-one relationship. Therefore, in the case II dataset, many-to-many CCIs such as the CCIs corresponding to green L-R sets, are hard to detect by the method. This is because for each L-R pair, mean values for any combination of cell types are basically high in such situations, and a *P*-value corresponding to a one-to-one CCI tends to be large (i.e., not significant); accordingly, the observed L-R coexpression and the null distribution calculated are hard to distinguish. In the analysis of real datasets presented later, however, the L-R gene expression pairs are not always the cell-type specific, and it is more natural that the CCI corresponding to the L-R has a many-to-many relationship. This simulation shows that `scTensor` is a more general method for detecting CCIs and their related L-R pairs at once, irrespective of whether a particular CCI is one-to-one or many-to-many.

#### Biological interpretation of real datasets

To demonstrate the efficacy of `scTensor` in the analysis of empirical datasets, we applied `scTensor` to four real scRNA-seq datasets (Table 1). First, we used the scRNA-seq data derived from fetal germ cells (FGCs (♀)) and their gonadal niche cells (Soma) from female human embryos (Germline Female [25]). As a known *CaH* pattern, CCIs of Soma with FGCs (♀) involving the BMP signaling pathway are reported, and `scTensor` accurately detects the CCIs and CCI-related L-R pairs as *CaH* (4,2,5) (Figure 3). Moreover, `scTensor` was able to extract some putative *CaH* s such as *CaH* (3,4,14) and *CaH* (1,4,22). *CaH* (3,4,14) is the autocrine-type CCI within FGCs (♀) and L-R pairs such as Wnt (WNT5A/WNT6) and some growth factor genes (NGF/IGF/FGFR/VEGF), suggesting that this *CaH* is related to the proliferation and differentiation of Soma. *CaH* (1,4,22) is the CCI corresponding to FGCs (♀) and Soma, and most of the receptors are related to G protein-coupled receptor (GPCR). The conjugated ligands are some neuropeptides, and this suggests that the peptides are related to the activation of GPCR in FGCs (♀) by some mechanism.

**Figure 3.**
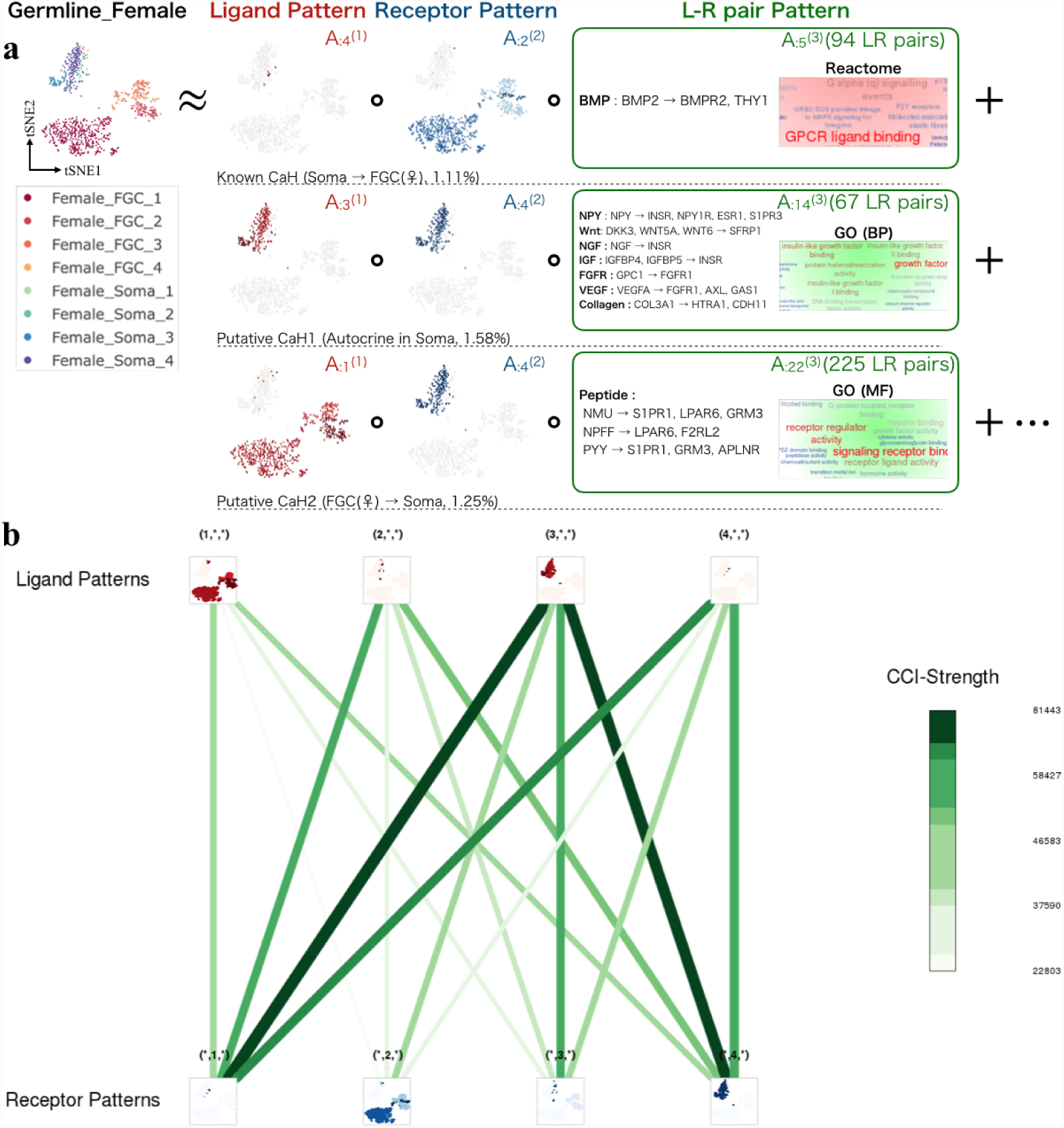
Results of scTensor for an empirical dataset (Germline Female) (a) Detected *CaH*s. (b) Overview of the L-R patterns.

We also used the scRNA-seq data derived from FGCs (♂) and their gonadal niche cells from male human embryos (Germline Male [25]). In contrast to the Germline Female data, the original study reported that FGCs (♂) interact with Soma through AMH-BMPR interactions, and `scTensor` also detected the triadic relationship as *CaH* (1,1,10) (Figure 4). The conjugate receptor AMHR2 is also detected in this *CaH*. In this dataset, like in Germline Female, in this data, autocrinetype CCIs such as *CaH* (3,3,9) and CCIs corresponding to FGCs (♂) and Soma (*CaH* (1,3,16)) were detected.

**Figure 4.**
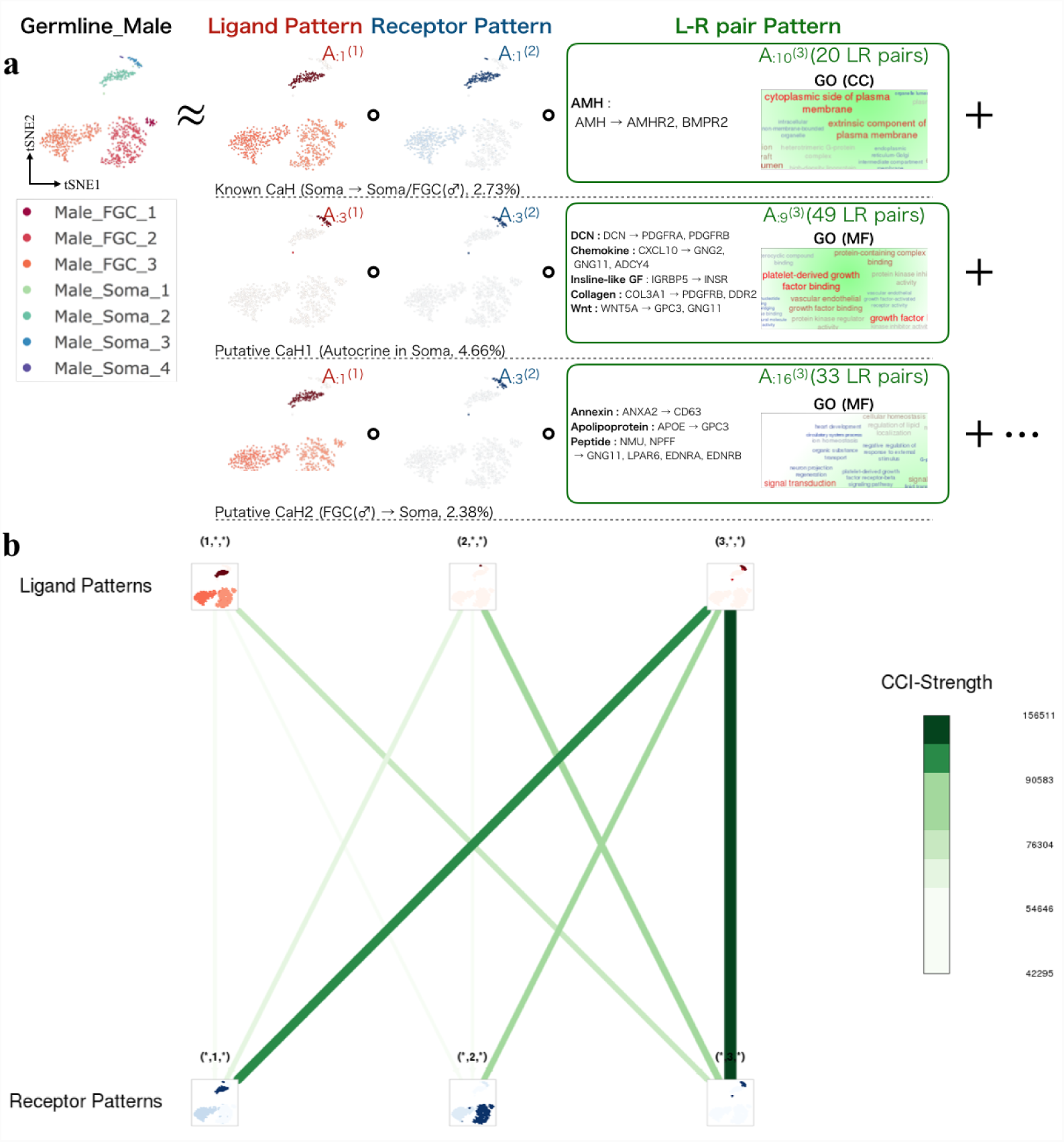
Results of scTensor for an empirical dataset (Germline Male) (a) Detected *CaH*s. (b) Overview of the L-R patterns.

Next, we used the scRNA-seq data derived from immune cells isolated from the metastatic melanoma patients (Melanoma [59]). The meta-analysis of scRNA-seq and TCGA datasets in the original published study claimed that T cell abundance and the expression of complement system-related genes in cancer-associated fibroblasts (CAFs) are correlated, and CCIs between T cells and CAFs were therefore inferred. `scTensor` detected these CCIs as *CaH* (1,1,3)/*CaH* (1,1,9) and complement system-related genes such as C3, CXCL12, CFB, and C4A were also detected (Figure 5). `scTensor` was also able to capture a well-known CCI between T cells and B cells as *CaH* (2,2,6), although this was not a focus of the original study; this CCI is the antigen-presenting from B cells to T cells by class II major histocompatibility complex (MHC) through the coexpression of CD8 (T cell receptor) and human leukocyte antigen (HLA) genes. `scTensor` also detected *CaH* (4,1,13) and *CaH* (1,4,8), which are the CCIs of macrophages with T cells and an autocrine-type CCI of known chemokine ligands with their receptors.

**Figure 5.**
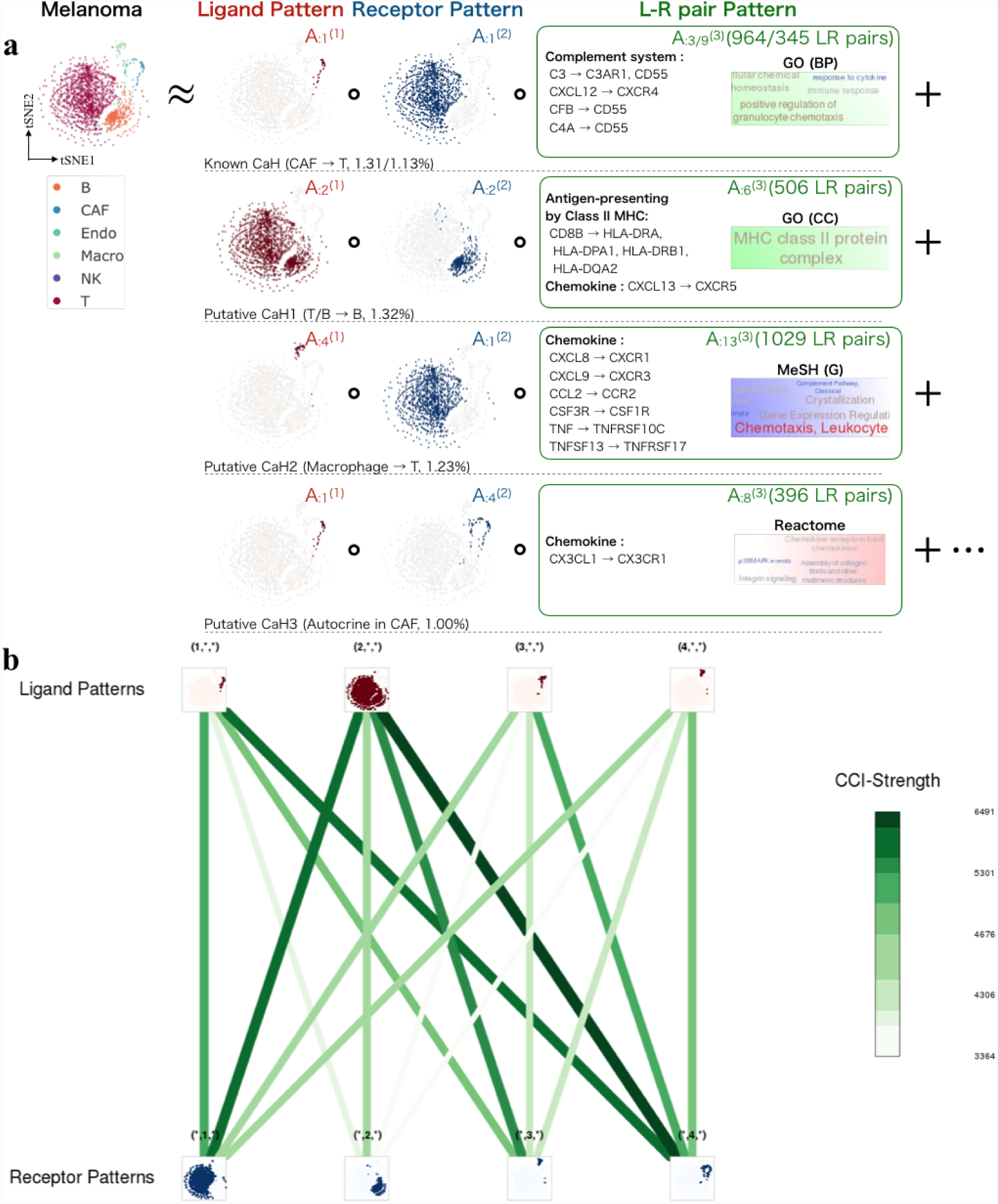
Results of scTensor for an empirical dataset (Melanoma) (a) Detected *CaH*s. (b) Overview of the L-R patterns.

Finally, we used data derived from non-myocyte cells isolated from mouse heart (NonMyocyte [27]). The original study focused on the CCIs of pericytes/fibroblasts with macrophages through the L-R pairs Il34/Csf1 and Csf1r, and the corresponding CCIs were detected as *CaH* (2,2,19) by `scTensor` (Figure 6). `scTensor` also detected *CaH* (1,2,17) and *CaH* (3,2,21), which are the autocrine-type CCI among macrophages and the CCI of NK cells with macrophages by known chemokine ligands and their receptors.

**Figure 6.**
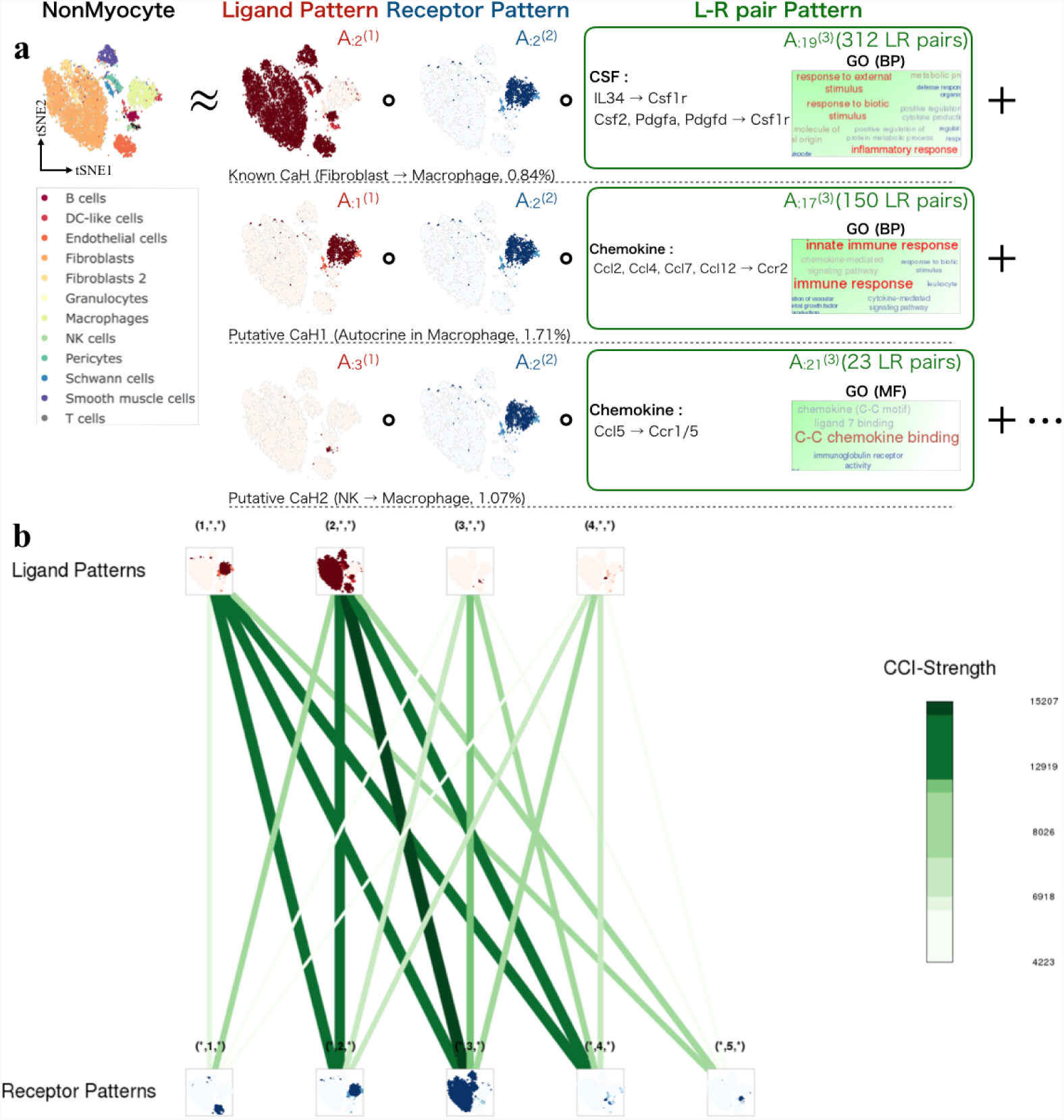
Results of scTensor for an empirical dataset (NonMyocyte) (a) Detected *CaH*s. (b) Overview of the L-R patterns.

### Application to minor organism

To demonstrate the applicability of using `scTensor` in the species that is not mouse or human, we used a scRNA-seq data derived from zebrafish habenular neurons (Habenular Larva [60]). Although the original study did not focus on the CCIs among the neuronal cell types, `scTensor` detected some triadic relationships as *CaH* (3,3,4) (La Hb01/03/07 with La Hb02/08), *CaH* (2,1,3) (La Hb09 with La Hb02/08), and *CaH* (1,3,1) (La Hb02/04-06/08/11-15/Olf with La Hb02/08) (Figure 7). The spatial distribution of the cell types measured by RNA-fluorescence *in situ* hybridization (FISH) shows that the cell-type pairs detected as *CaH* (3,3,4) and *CaH* (2,1,3) are dorsally located and proximal to each other in the habenula. However, *CaH* (1,3,1) was a more global interaction related to the entire habenula regions. Although the spatial distribution of rare cell types La Hb03/05/14 could not be determined by RNA-FISH in the original study, `scTensor` was able to assign the conjugated cell types of La Hb03 as La Hb02/08 in the dorsal region. This result suggests that `scTensor` may also be useful in spatial transcriptomics [61, 62].

**Figure 7.**
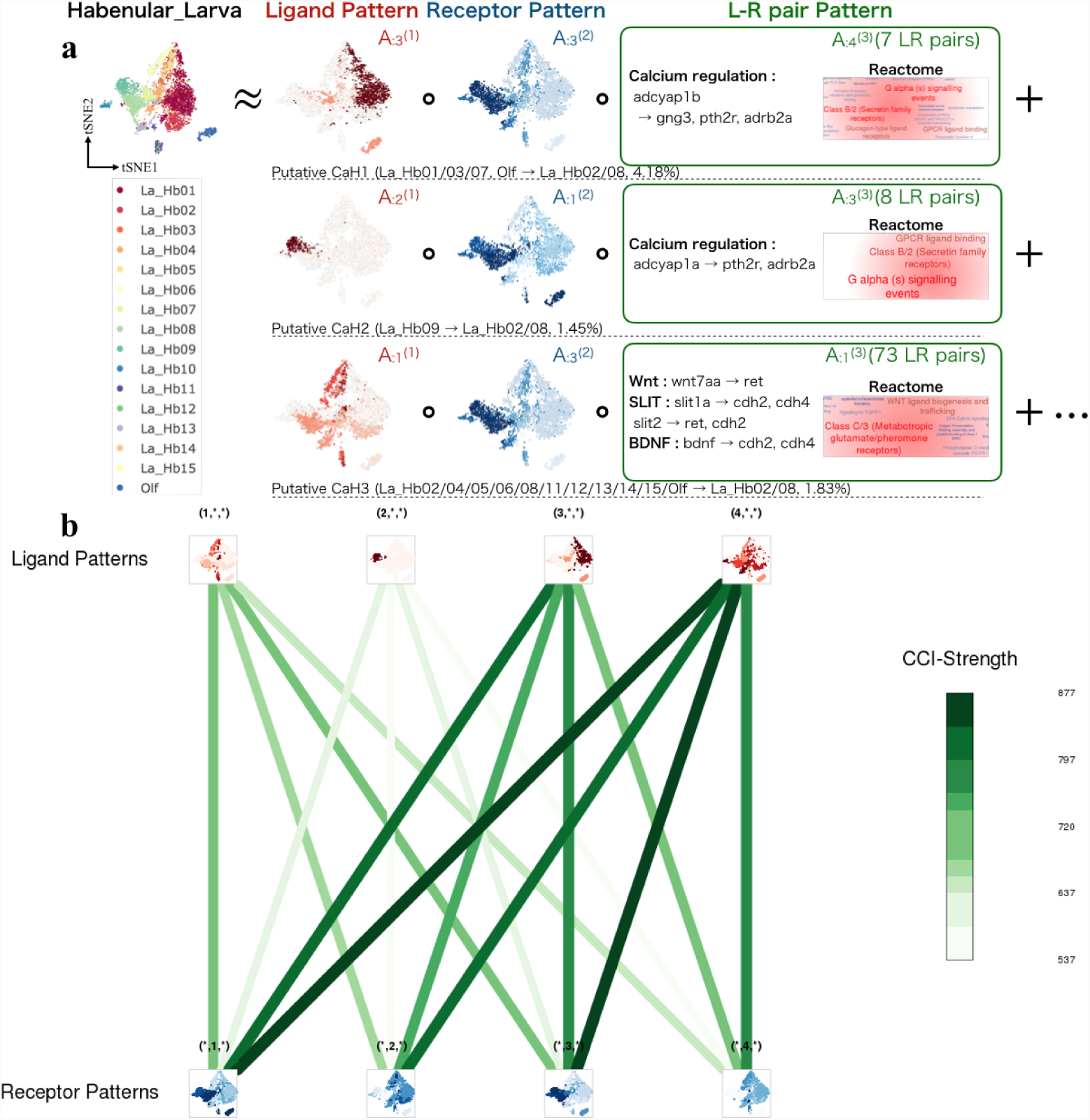
Results scTensor for an empirical dataset (Habenular Larva) (a) Detected *CaH*s. (b) Overview of the L-R patterns.

### scTensor and L-R database implementations as R/Bioconductor packages

All the algorithms and L-R lists are available as R/Bioconductor packages and a web application described below.

### nnTensor and scTensor packages

NTD is implemented as the function of `nnTensor` R/CRAN package and internally imported in `scTensor`. `scTensor` constructs the CCI-tensor, decomposes the tensor by NTD, and generates an HTML report. `scTensor` is assumed to be used with `LRBase.XXX.eg.db`, which are the L-R databases for multiple organisms. To enhance the biological interpretation of *CaH* s, a wide variety of gene information is assigned to the L-R lists through the other R/Bioconductor packages (Figure 8). For example, gene annotation is assigned by `biomaRt` [63] (Gene Name, Description, Gene Ontology (GO), STRING, and UniProtKB), `reactome.db` [64] (Reactome) and `MeSH.XXX.eg.db` [65] (Medical Subject Headings; MeSH), while the enrichment analysis (also known as over-representative analysis; ORA) is performed by `GOstats` [66] (GO-ORA), `meshr` [65] (MeSH-ORA), `ReactomePA` [67] (Reactome-ORA), and `DOSE` [68] (Disease Ontology; DO, Network of Cancer Gene; NCG, and DisGeNET-ORA). To validate that the detected gene expression of L-R gene pair is also consistently detected in the other data with tissue- or cell-type-level transcriptome data, the hyperlinks to RefEx [69], Expression Atlas [70], Single-Cell Expression Atlas [71], scRNASeqDB [72], PanglaoDB [73] are embedded in the HTML report, facilitating comparisions of the L-R expression with the data from large-scale genomics projects such as GTEx [74], FANTOM5 [75], NIH Epigenomics Roadmap [76], ENCODE [77], and Human Protein Atlas [78]. Additionally, in consideration of users who might want to experimentally investigate detected CCIs, we embedded the hyperlinks to Connectivity Map (CMap [79]), which provides the relationships between perturbation by the addition of particular chemical compounds/genetic reagents and succeeding gene expression change.

**Figure 8.**
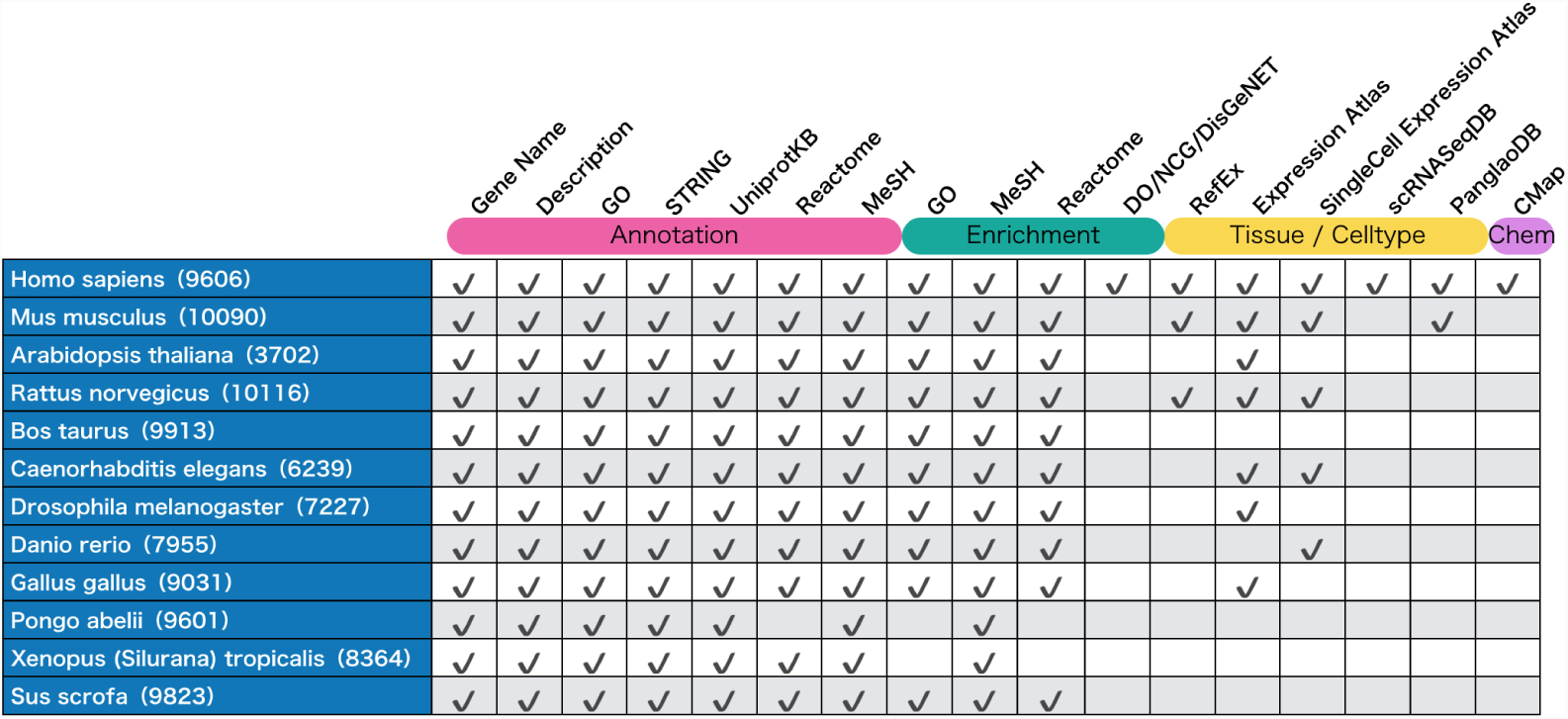
Correspondence table containing the available L-R gene information for each organism. For each detected *CaH*, gene annotation, enrichment analysis, tissue- or cell-type-specific gene expression, and chemical-gene expression relationships are assigned.

### LRBase.XXX.eg.db-type packages

For data sustainability and the extension to the wide range of organisms, in this work, we originally constructed L-R databases as R/Bioconductor packages named `LRBase.XXX.eg.db` (where XXX represents the abbreviation for an organism, such as “Hsa” for Homo sapiens). `LRBase.XXX.eg.db` currently provides the L-R databases for 12 organisms (Table 2 and 3). The data process pipeline is almost the same as that of the FANTOM5 project for constructing the putative L-R lists. Precise differences between `LRBase.XXX.eg.db` and FANTOM5 are summarized in Table S4 in Additional File 1.

### LRBaseDbi package

All the `LRBase.XXX.eg.db` packages are generated by `LRBaseDbi`, which is the another R/Bioconductor package. `LRBaseDbi` generates the `LRBase.XXX.eg.db` packages from the CSV files, in which NCBI Gene IDs are saved as two columns describing the L-R relationship (we call this function as “meta”-packaging).

In addition to the `LRBase.XXX.eg.db` packages we summarized, the users may want to specify the user’s original L-R list. For example, there are some other L-R databases such as IUPHAR [80], DLRP [81], FANTOM5 [75], CellPhoneDB [28], and Cell-Cell Interaction Database summarized by Gary Bader et. al. (http://baderlab.org/CellCellInteractions) (Additional File 1). Besides, if the users want to apply the `scTensor` to minor species, the L-R list may be constructed by the orthologous relationship with major species (e.g., human) [27]. The corresponding `LRBase.XXX.eg.db` is easily generated from the original L-R list by `LRBaseDbi` and can be used with `scTensor`.

### CellCelldb

All analytical results `scTensor` are outputted as HTML reports. Hence, combined with a cloud web service such as Amazon Simple Storage Service (Amazon S3), reports can be used as simple web applications, enabling the user to share their results with collaborators or to develop an exhaustive CCI database. We have already performed `scTensor` analyses using a wide-variety of scRNA-seq datasets, including the five empirical datasets examined in this study (https://q-brain2.riken.jp/CellCelldb/).

## Conclusions

In this work, CCIs were regarded as *CaH* s, which are hypergraphs that represent triadic relationships, and a novel algorithm `scTensor` for detecting such *CaH* was developed. In evaluations with empirical datasets from previous CCI studies, previously reported CCIs were also detected by `scTensor`. Moreover, some CCIs were detected only by `scTensor`, suggesting that the previous studies may have over-looked some CCIs. To extend the use of `scTensor` to a wide range of organisms, we also developed multiple L-R databases as `LRBase.XXX.eg.db`-type packages. When combined with `LRBase.XXX.eg.db`, `scTensor` can currently be applied to 12 organisms.

There are still some plans for improving both `LRBase.XXX.eg.db` and `scTensor` to build on the advantages of this current framework. For example, the range of corresponding organisms and the L-R lists can be extended with the spread of genome-wide researches. Additionally, the algorithm can be improved, for example, by utilizing acceleration techniques such as randomized algorithm/sketching methods [82], incremental algorithm/stochastic optimization [83, 84], or distributed computing with MapReduce/Hadoop on large-scale memory machines [85] for NTD, which is now available. Tensors are a very flexible way to represent heterogeneous biological data [86], and easily integrate the side information about genes or cell types with semi-supervised manner. Such information will improve the accuracy and extend the scope of the data.

We aim to tackle such problems and develop the framework further through updates of the R/Bioconductor packages. In the package registration process for R/Bioconductor, package source code is peer-reviewed via the Bioconductor single package builder system and assigned to a curator (https://github.com/ Bioconductor/packagebuilder), and even after the package is accepted, the daily package builder tests the source code every day (https://www.bioconductor.org/checkResults/. Furthermore, biannual updates of Bioconductor require that the internal data of all data packages are updated (https://www.bioconductor.org/ developers/release-schedule/). Such strict check systems for source code and internal data improve the sustainability and usability of packages. Our team has maintained over one hundred R/Bioconductor packages since 2015 [65], and we are still organizing a system for the maintenance of the combined `LRBase.XXX.eg.db` and `scTensor` framework.

## Materials and methods

### Construction of L-R list

#### Public databases

To compare our database with other databases (Additional File 1 and 2), the data from FANTOM5 (http://fantom.gsc.riken.jp/5/suppl/Ramilowski$_$et$_$al$_$2015/data/PairsLigRec.txt), DLRP (http://dip.doe-mbi.ucla.edu/dip/dlrp/dlrp.txt), IUPHAR (http://www.guidetopharmacology.org/DATA/interactions.csv), and HPRD (http://ftp://ftp.ebi.ac.uk/pub/databases/genenames/new/tsv/locus$_$groups/protein-coding$_$gene.txt) were downloaded. The subcellular localization data from SWISSPROT and TrEMBL were downloaded from UniProtKB (https://www.uniprot.org/downloads). Protein-protein interaction (PPI) data of STRING (v-10.5) were downloaded from https://stringdb-static.org/cgi/download.pl. To unify the gene identifier as NCBI Gene ID, we retrieved the corresponding table from `Biomart` (Ensembl release 92). All the data were downloaded by RESTful access using the wget command and query.xml http://www.biomart.org/martservice.html.

#### Simulation datasets

The simulated single-cell gene expression data were sampled from the negative binomial distribution *NB* (*FC*_*gc*_*m*_*g*_, *ϕ*_*g*_), where *FC*_*gc*_ is the fold-change (FC) for gene *g* and cell type *c*, and *m*_*g*_ and *ϕ*_*g*_ are the average gene expression and the dispersion parameter of gene *g*, respectively. For the setting of differentially expressed genes (DEGs) and non-DEGs, *FC*_*gc*_ values were calculated based on the non-linear relationship of FC and the gene expression level log10 *FC*_*gc*_ = *a* exp(*-b* log10 (*m*_*g*_ + 1)) as follows:

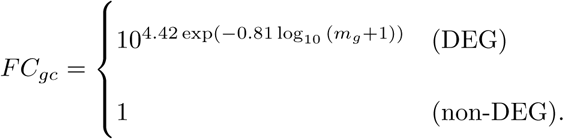

The *m*_*g*_ and gene-wise variance *v*_*g*_ were calculated from the scRNA-seq dataset of human embryonic stem cells (hESCs) measured by Quartz-Seq [87], and the gene-wise dispersion parameter *ϕ*_*g*_ was estimated as 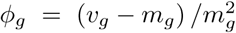. The NB distribution reduces to Poisson when *ϕ*_*g*_ = 0. To simulate the “dropout” phenomena of scRNA-seq experiments, we introduced the dropout probability 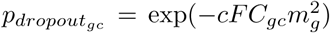, which is used in `ZIFA` [88] (default: *c*=1), and the expression values were randomly converted to 0 based on this dropout probability.

For the setting of the case I datasets, 150 *×* 150 *×* 500 CCI-tensor was constructed. For each cell type, 50 cells were established, and in total, three cell types were set. For L-R set 1 (red), 50 L-R pairs were established, and the cell-type-wise ligand and receptor patterns were (1,0,0) and (0,1,0), respectively (0, non-DEG; 1, DEG). For L-R set 2 (blue), 50 L-R pairs were established, and the cell-type-wise ligand and receptor patterns were (0,1,0) and (0,0,1), respectively. For L-R set 3 (green), 50 L-R pairs were established, and the cell-type-wise ligand and receptor patterns were (0,0,1) and (1,0,0), respectively. The other 350 L-R pairs were sampled randomly as non-DEGs.

For the setting of the case II datasets, 150 *×* 150 *×* 500 CCI-tensor were constructed. For each cell type, 50 cells were established, and in total, three cell types were established. For L-R set 1 (red), 50 L-R pairs were established, and the cell-type-wise ligand and receptor patterns were (1,0,0) and (0,1,1), respectively. For L-R set 2 (blue), 50 L-R pairs were established, and the cell-type-wise ligand and receptor patterns were (0,0,1) and (1,1,1), respectively. For L-R set 3 (green), 50 L-R pairs were established, and the cell-type-wise ligand and receptor patterns were (1,1,1) and (1,0,1), respectively. The other 350 L-R pairs were sampled randomly as non-DEGs.

#### Real datasets

The gene expression matrix and cellular labels for Germline Female and Germline Male scRNA-seq data were retrieved from the GEO database (GSE86146), and only highly variable genes (HVGs: http://pklab.med.harvard.edu/scw2014/subpoptutorial.html) with low *P*-values (≤ 1E-7) were extracted. The gene expression matrix and cellular label of Melanoma scRNA-seq data were retrieved from the GEO database (GSE72056), and only HVGs with low *P*-values (≤ 1E-10) were extracted. The gene symbols were converted to NCBI GeneIDs using the R/Bioconductor *Homo.sapiens* package. The gene expression matrix and cellular labels for NonMyocyte scRNA-seq data were retrieved from the ArrayExpress database (E-MTAB-6173), and only HVGs with low *P*-values (≤ 1E-10) were extracted. The gene symbols are converted to NCBI GeneIDs using the R/Bioconductor *Mus.musculus* package. For each dataset, t-Distributed Stochastic Neighbor Embedding (t-SNE) with 40 perplexity was performed using the Rtsne R package.

### scTensor algorithm details

#### Construction of CCI-tensor

Here we assume that a data matrix ***Y*** ∈ ℝ^*I×H*^ is the gene expression matrix of scRNA-seq, where *I* is the number of genes and *H* is the number of cells. Next, the matrix ***Y*** is converted to cell-type mean matrix ***X*** ∈ ℝ^*I×J*^, where *J* is the number of mean vectors for each cell type. The cell-type label is supposed to be specified by user’s prior analysis such as clustering or confirmation of marker gene expression.

The relationship between the ***X*** and ***Y*** is described as below:

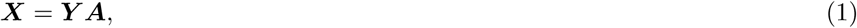

where the matrix ***A*** ∈ ℝ^*H×J*^ converts cellular-level matrix ***Y*** to cell-type-level matrix *X* and each element of ***A*** is

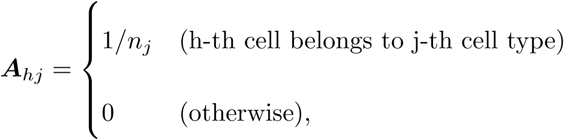

where *n*_*j*_ is the number of cells belonging to *j*’s cell type.

Next, NCBI gene IDs of each L-R pair stored in `LRBase.XXX.eg.db` are searched in the row names of matrix *X* and if both IDs are found, corresponding *J*-length row-vectors of the ligand and receptor genes (***x***_*L*_ and ***x***_*R*_) are extracted.

Finally, a *J × J* matrix is calculated as the outer product of ***x***_*L*_ and ***x***_*R*_ and incrementally stored as a sub-tensor (frontal slice) of the CCI-tensor *χ* ∈ ℝ^*J×J×K*^ as below:

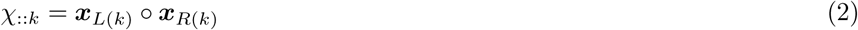

where *K* is the number of L-R pairs found in the row names of matrix ***X***.

#### CANDECOMP/PARAFAC and Tucker decomposition

Here, we suppose that the CCI-tensor has some representative triadic relationship. To extract the triadic relationships from a CCI-tensor, here we consider performing some tensor decomposition algorithms. There are two typical decomposition methods; CANDECOMP/PARAFAC (CP) and Tucker decomposition [57, 58].

In CP decomposition, CCI-tensor *χ* is decomposed as follows:

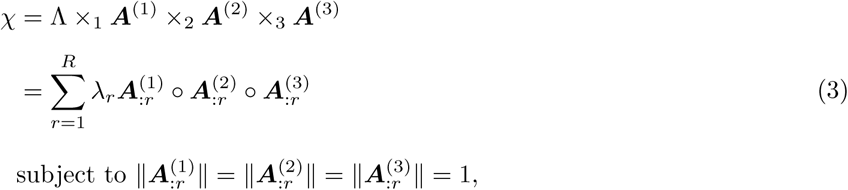

where *×*_*n*_ is mode-*n* product, *R* is the rank of *χ*, and Λ is diagonal cubical tensor, in which the element *λ*_*r*_ on the superdiagonal can be non-zero. ***A***^(1)^ ∈ ℝ^*J×R*^, ***A***^(2)^ ∈ ℝ^*J×R*^, and ***A***^(3)^ ∈ ℝ^*K×R*^ are factor matrices. ***a***_*r*_ *○* ***b***_*r*_ *○* ***c***_*r*_ is rank-1 tensor, and the scalar *λ*_*r*_ is the size of rank-1 tensor. The rank-1 tensor indicates the triadic relationship described above, and CP model suppose that CCI-tensor is approximated by the summation of *R* rank-1 tensor. There are some algorithms for optimizing CP decomposition problem such as alternative least squares (ALS) or power method [58]. Despite its wide use, CP decomposition has a drawback when using the problem in this work; the number of columns of three factor matrices must be a common number *R*, and the correspondence of 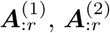, and 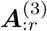 in each *r* is one-to-one. This constraint is sometimes too strict and unnatural for biological applications. For example, in the CCI-tensor case, the number of ligand expression, receptor expression, and L-R-pair patterns are commonly *R*, and all of them must correspond to each other in each *r*. Thus, if an L-R pair assigned in a CCI, this L-R pair cannot be part of other CCIs.

To deal with this problem, the application of Tucker decomposition can be considered next. In Tucker decomposition, a CCI-tensor is decomposed as follows:

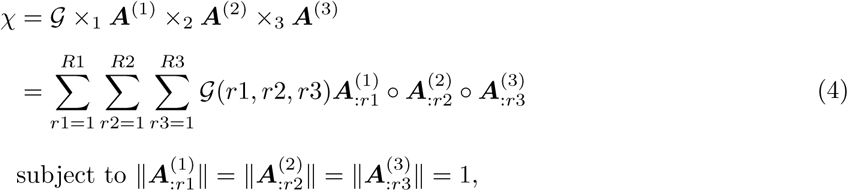

where *R*1, *R*2, and *R*3 are the rank of mode-1,2, and 3, respectively. Unlike the CP model, the constraint conditions of the Tucker model are relaxed, that is, three factor matrices ***A***^(1)^ ∈ ℝ^*J×R*1^, ***A***^(2)^ ∈ ℝ^*J×R*2^, and ***A***^(3)^ ∈ ℝ^*K×R*3^ can differ in their numbers of columns and any combination of 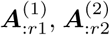, and 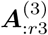 can be considered. This is because 𝒢 ∈ ℝ^*R*1*×R*2*×R*3^ is a dense core tensor and any element, including non-diagonal elements, can have a non-zero value. There are some algorithms for optimizing Tucker decomposition, such as higher order singular value decomposition (HOSVD) or higher orthogonal iteration of tensors (HOOI) [58].

#### Non-negative Tucker decomposition

Despite its effectiveness, Tucker decomposition cannot be directly applied to the extraction of *CaH* s. This is because the factor matrices of the Tucker model can have negative value elements, and these make interpretation difficult. For example, if 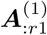 contains very large positive elements and very small negative elements, we cannot determine which cell type is highly related to a ligand expression pattern and which cell type is not.

For the above reason, here we utilize NTD. Unlike Tucker decomposition based on singular value decomposition (SVD), NTD is based on non-negative matrix factorization (NMF), which is another matrix decomposition method. NMF is formalized as follows:

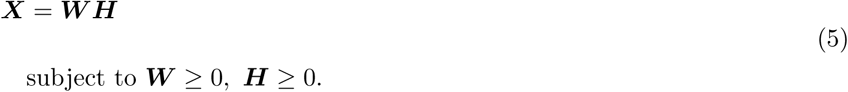

The typical algorithm for optimizing the NMF problem is multiplicative updating (MU) [58]. Two widely used forms are considered. The first form is minimization problem of Euclidean distance (*min* ‖ ***X*** *–* ***WH*** ‖ _*Euclid*_), where ***H*** and ***W*** are iteratively updated by considering Gaussian noise:

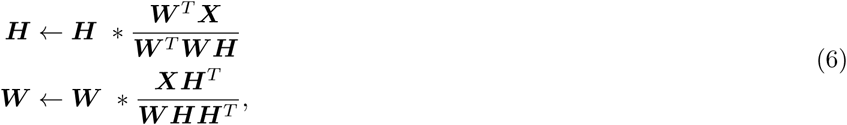

where *∗* is the element-wise (Hadamard) product. The second form is a minimization problem of Kullback-Leibler (KL) divergence (*min* ‖***X*** *-* ***WH***‖_*KL*_), where ***H*** and ***W*** are iteratively updated by considering Poisson noise:

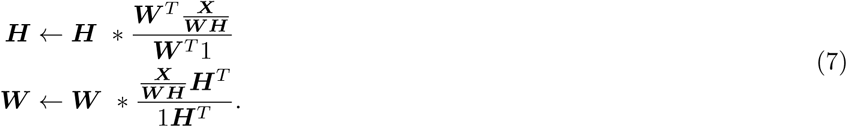

These update rules are derived from the element-wise gradient descent with the spatial form of the learning rate [58]. Starting with a random non-negative initial value, update of ***W*** and ***H*** are updated iteratively until convergence. In this work, MU with the KL-form, which shows a stable convergence with simulation data, is used for following initialization step of NTD (for more details, see Additional File 3).

To extend the KL form of MU to NTD, we consider iterative updating ***A***^(*n*)^𝒢_*n*_***A***^(*-n*)*T*^, which is the matricized expression of Tucker decomposition. Here ***A***^(*-n*)^ represents the factor matrices without ***A***^(*n*)^. For example, if *n*=1, this part becomes ***A***^(2)*T*^ ***A***^(3)*T*^. By considering a part of ***A***^(*n*)^ 𝒢_*n*_***A***^(*-n*)*T*^ as a variable and fixing other parts as constants, the KL form of MU can be performed to the matricized tensor, such as ***A***^(1)^ *→* ***A***^(2)^ *→* ***A***^(3)^ *→* 𝒢 *→ · · ·*. Each updating rule for ***A***^(*n*)^ is as follows:

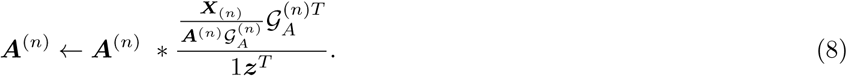

Additionally, the updating rule for core tensor 𝒢 is:

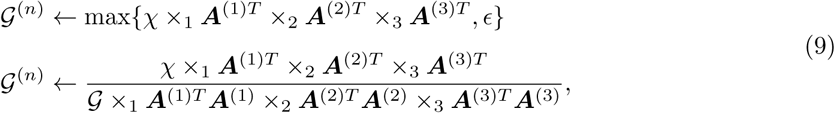

where *ϵ* is a small value included to avoid generating negative values. In the `nnTensor`, 1e-10 is used.

#### Extraction of CCIs as hypergraphs

To extract the representative *CaH* s, `scTensor` estimates the NTD ranks by SVD performed for each matricized CCI-tensor (*X*^(*n*)^, n=1,2, and 3). The eigenvalues and eigenvectors that explain the top 80% to 90% of variance are selected. With the estimated ranks of 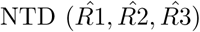, NTD is performed, and only the triads (r1,r2,r3) with large core tensor values are selected as representative *CaH* s. In its default mode, `scTensor` selects the *CaH* s that explain the top 40% of cumulative core tensor values. For each *CaH*, corresponding column vectors of factor matrices were selected as 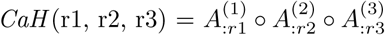.

To enhance the interpretation, each column vector is binarized in advance by two-class hierarchical clustering using Ward’s minimum variance method, and only large values are converted to 1, with other values becoming 0.

CCI-strength (cf. Figure 3, 4, 5, 6, 7) is calculated as the summation of mode-3 of reconstructed tensor from all *CaH* s. With the selected indexes in each mode 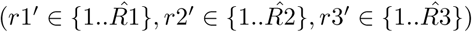, CCI-strength is defined as follows:

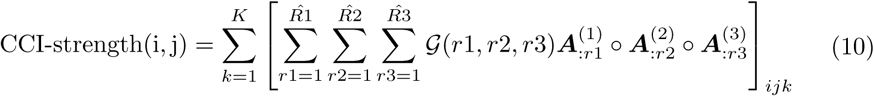

#### Label permutation method

In this method, the cluster labels of all cells are randomly permuted 1000 times, and the average ligand expression level of a cluster and the average receptor expression level of a cluster are calculated [37]. For each L-R pair, the mean values of the averaged L-R expression level are calculated in all possible combinations of the cell types. This process generates 1,000 of synthetic L-R coexpression matrices and these are used to generate the null distribution, that is, in a combination of cell types, the proportion of the means which are “as or more extreme” than the observed mean is the calculated as *P*-value.

## Availability and requirements

- `scTensor`: https://bioconductor.org/packages/devel/bioc/html/scTensor.html
- `nnTensor`: https://cran.r-project.org/web/packages/nnTensor/index.html
- `LRBase.Hsa.eg.db:` https://bioconductor.org/packages/release/bioc/html/LRBase.Hsa.eg.db.html
- `LRBase.Mmu.eg.db:` https://bioconductor.org/packages/release/bioc/html/LRBase.Mmu.eg.db.html
- `LRBase.Ath.eg.db:` https://bioconductor.org/packages/release/bioc/html/LRBase.Ath.eg.db.html
- `LRBase.Rno.eg.db:` https://bioconductor.org/packages/release/bioc/html/LRBase.Rno.eg.db.html
- `LRBase.Bta.eg.db:` https://bioconductor.org/packages/release/bioc/html/LRBase.Bta.eg.db.html
- `LRBase.Cel.eg.db:` https://bioconductor.org/packages/release/bioc/html/LRBase.Cel.eg.db.html
- `LRBase.Dme.eg.db:` https://bioconductor.org/packages/release/bioc/html/LRBase.Dme.eg.db.html
- `LRBase.Dre.eg.db:` https://bioconductor.org/packages/release/bioc/html/LRBase.Dre.eg.db.html
- `LRBase.Gga.eg.db:` https://bioconductor.org/packages/release/bioc/html/LRBase.Gga.eg.db.html
- `LRBase.Pab.eg.db:` https://bioconductor.org/packages/release/bioc/html/LRBase.Pab.eg.db.html
- `LRBase.Xtr.eg.db:` https://bioconductor.org/packages/release/bioc/html/LRBase.Xtr.eg.db.html
- `LRBase.Ssc.eg.db:` https://bioconductor.org/packages/release/bioc/html/LRBase.Ssc.eg.db.html
- `LRBaseDbi`: https://bioconductor.org/packages/release/bioc/html/LRBaseDbi.html
- Operating system: Linux, Mac OS X, Windows
- Programming language: R (v-3.5.0 or higher)
- License: Artistic-2.0
- Any restrictions to use by non-academics: For non-profit use only

## Abbreviations

CCI: cell-cell interaction; scRNA-seq: single-cell RNA sequencing; L-R: ligand and receptor; *CaH*: CCI as hypergraph; FTT: Freeman-Tukey transformation; NTD: non-negative Tucker decomposition; ROC: receiver operating characteristic; AUC: area under the curve; FGCs: fetal germ cells; Soma: gonadal niche cells; GPCR: G protein-coupled receptor; CAFs: cancer-associated fibroblasts; MHC: major histocompatibility complex; HLA: human leukocyte antigen; RNA-FISH: RNA-fluorescence *in situ* hybridization; PM: plasma membrane; PPI: protein-protein interaction; GO: Gene Ontology; MeSH: medical subject headings; ORA: over-representative analysis; DO: Disease Ontology; NCG: network of cancer gene; CMap: Connectivity Map; Amazon S3: Amazon simple storage service; FC: fold-change; DEGs: differentially expressed genes; human embryonic stem cells: hESCs; HVGs: highly variable genes; t-SNE: t-distributed stochastic neighbor embedding; CP: CANDECOMP/PARAFAC; ALS: alternative least squares; HOSVD: higher order singular value decomposition; HOOI: higher orthogonal iteration of tensors; NMF: non-negative matrix factorization; MU: multiplicative updating; KL: Kullback-Leibler

## Competing interests

The authors declare that they have no competing interests.

## Funding

This work was supported by MEXT KAKENHI Grant Number 16K16152 and by JST CREST grant number JPMJCR16G3, Japan.

## Authors’ contributions

KT and IN designed the study. KT designed the algorithm and benchmark, retrieved and preprocessed the test data to evaluate the proposed method, implemented the source code, and performed all analyses. MI implemented the pipeline for bi-annual automatic update of the R/Bioconductor packages. All authors have read and approved the manuscript.

## Acknowledgements

Some cell images used in Figure 1 were previously presented by ©2016 DBCLS TogoTV. We thank Dr. Yoshihiro Taguchi for discussions about the algorithms. We thank Mr. Akihiro Matsushima for their assistance with the IT infrastructure for the data analysis. We are also grateful to all members of the Laboratory for Bioinformatics Research, RIKEN Center for Biosystems Dynamics Research for their helpful advice.

## Additional Files

Additional file 1 — Development of L-R databases for multiple organisms (PDF 2.2 MB)

Additional file 2 — Distributions of and correlations among 8 STRING-scores (ZIP 18.5 MB)

Additional file 3 — Convergence of NTD with toy model and empirical data (PDF 9.5 MB)

