## Additional File 1 for "Uncovering hypergraphs of cell-cell interaction from single cell RNA-sequencing data"

### Supplementary material

#### Content

*SUPPLEMENTARY ANALYSES* ........ *2*

Construction of LRBase.XXX.eg.db-type packages ........ 2

### SUPPLEMENTARY ANALYSES

#### Construction of LRBase.XXX.eg.db-type packages

Many CCI studies based on ligand-receptor (L-R) gene coexpression utilize data from the FANTOM5 project [23], which was constructed using the following pipeline.

First, the project retrieved candidate L-R pairs according to two methods.

- ***Subcellular Localization***: The terms "Secreted" and "Plasma Membrane" were extracted from the UniProtKB [64] and HPRD [65] databases and were regarded as candidate ligand and receptor genes, respectively. Computational prediction by LocTree3 [66] and PolyPhobius [67] was also performed.
- ***Physical Binding of Proteins***: Experimentally validated protein-protein interaction (PPI) information was extracted from HPRD and STRING [68] databases. From the STRING database, only PPI information with a 700 confidence score for "physical-binding interactions" and "experimental interactions" was used.

Next, the project extracted common gene pairs satisfying the two criteria described above and also merged existing L-R databases, including DLRP [62], IUPHAR [61], and HPMR [63].

Finally, related citations supporting the interaction were found, and interactions with and without PubMed ID were categorized as "known" and "putative" interactions, respectively.

Though the FANTOM5 data are useful, they still have the following problems.

- ***Sustainability***: Although the L-R list from FANTOM5 is open and anyone can access the data, it has not been maintained since July 2015 (http://fantom.gsc.riken.jp/5/suppl/Ramilowski_et_al_2015/data/). Based on our investigations, some source databases and ontologies used in the analytical pipeline from the original paper are obsolete or no longer accessible.
- ***Scope of Species***: FANTOM5 constructed only a human L-R list, but scRNA-Seq studies are also conducted in other species as well. Although Skelly et al. originally constructed the L-R list for mouse from FANTOM5 data by constructing orthologous mouse and human gene sets, it is still unclear that such an approach can be applied to other organisms. Homology may have been lost over the course of evolution, as there is a substantial genetic distance from human.

For the above reasons, we constructed L-R databases. Our workflow for detecting putative interactions was similar to that of FANTOM5. We first retrieved the annotation for subcellular localization from SWISSPROT (Figure S1). In addition to the manual curation of SWISSPROT, we also added predicted subcellular localization results. Although the FANTOM5 team constructed prediction tools themselves, we used TrEMBL annotation, which is the result of an automatic annotation pipeline using prediction tools such as TMHMM, SignalP, Phobius, and Coils (https://www.uniprot.org/help/sam).

Next, we retrieved PPI annotation from the STRING database. Eight evidence types were assigned to the annotation (Table S1); each evidence type has a confidence score ranging from 0 to 1000 (cf. http://version10.string-db.org/help/getting_started/, https://string-db.org/help//faq/#how-to-extract-high-confidence-07-interactions-from-information-on-combined-score-in-proteinlinkstxtgz). The values of this score are categorized as follows:

- Low confidence: 0–150
- Medium confidence: 400–700
- High confidence: 700–900
- Highest confidence: 900–1000

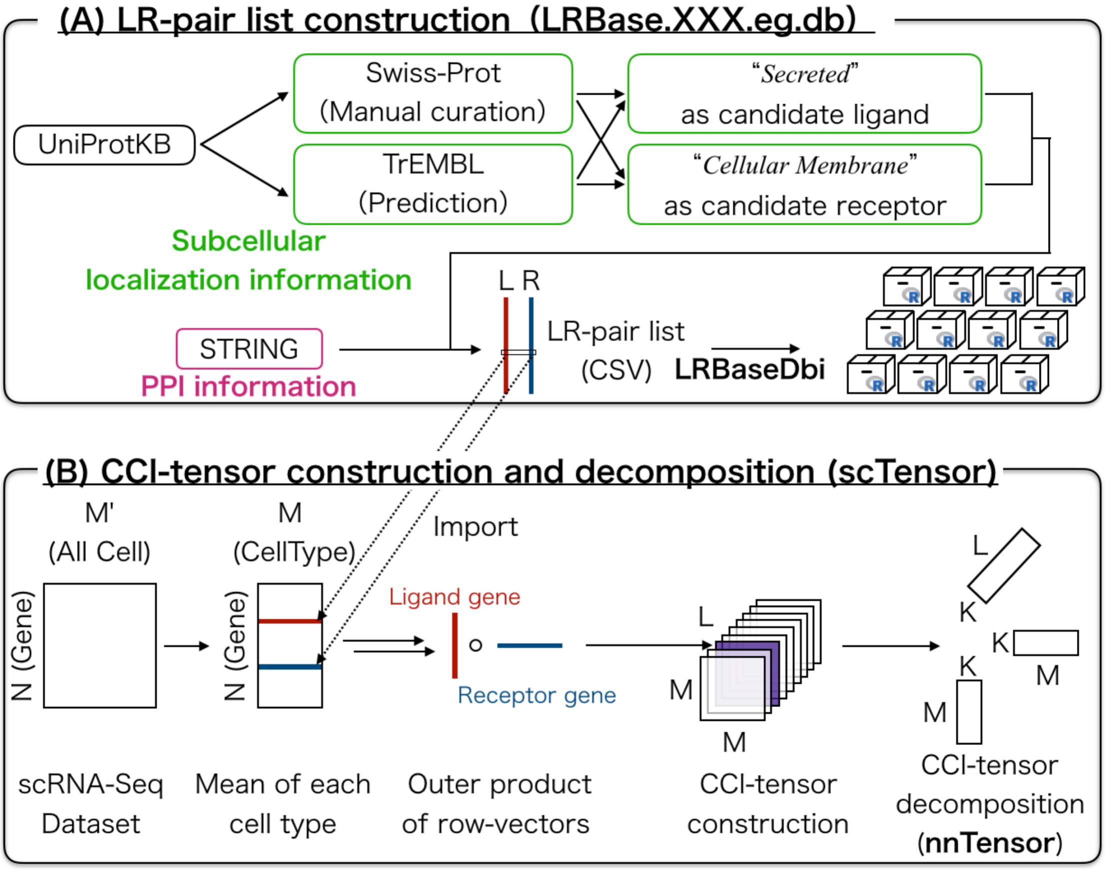


**Supplementary Figure 1 | Workflow for constructing LRBase.XXX.eg.db databases and relationships with other related packages** (A) LRBase.XXX.eg.db databases are constructed based on UniProtKB and STRING databases (B) scTensor internally imports LRBase.XXX.eg.db databases

**Supplementary Table 1 | Evidence Type from the STRING Database**

Eight evidence codes are defined in STRING.

| Association Evidence Type | Abstract |
|---|---|
| 1. *neighborhood* | Similar genomic context across different species implying similar function of the proteins. |
| 2. *fusion* | Gene fusions occurred in some genome implying functional relationship. |
| 3. *cooccurrence* | Phylogenetic co-occurrence across different species. |
| 4. *coexpression* | Co-expression detected from gene expression profile (e.g. microarray and RNA-Seq). |
| 5. *experimental* | Biological experiments in the lab such as yeast two-hybrid (Y2H). |
| 6. *database* | Imported from other pathway database (e.g. MINT, HPRD, BIND, ...and so on) asserted by a human expert curator. |
| 7. *textmining* | A large amount of scientific literatures (e.g. SGD, OMIM, FlyBase, PubMed) are parsed and associated by text-mining techniques. |
| 8. *combined* | Combined score of the scores of 1. to 7. |

Although two evidence codes "physical-binding interactions" and "experimental interactions" are used for FANTOM5, these evidence codes are no longer used in the current STRING database (perhaps having been merged as *experimental*). Here, we introduced a slightly moderate approach; we extracted the PPI list with over 400 *combined* confidence scores for usage across a wide range of organisms. Indeed, the combined score is highly correlated with *coexpression*, *experimental*, *database*, and *textmining* in many organisms (Additional File 2), and the same results are also expected with FANTOM5. Under such a condition, a relatively large number of elements were extracted from the PPI list of 12 organisms. Combined with the subcellular localization data from SWISSPROT and TrEMBL described above, the numbers of L-R pairs obtained for each organism are summarized in Tables S2 and S3.

Finally, the L-R databases for the 12 organisms were constructed as multiple R packages (Tables 2 and 3 in the main manuscript). These packages were generated from CSV files by the LRBaseDbi package. LRBaseDbi's roles are package generation and class definition of the accessor functions in LRBase.XXX.eg.db (Figure S1). The created LRBase.XXX.eg.db databases are specified in the scTensor package, which performs non-negative Tucker decomposition of nnTensor and extracts representative triadic relationship from CCI-tensors (Figure S1).

**Supplementary Table 2 | Total number of L-R pairs at each threshold of STRING combined scores (SWISSPROT)**

Manual annotation of subcellular localization in SWISSPROT was used.

| Organism | ≥150 | **≥400** | ≥700 |
|---|---|---|---|
| *Homo sapiens* | 87642 | **21882** | 14074 |
| *Mus musculus* | 79847 | **16386** | 9553 |
| *Arabidopsis thaliana* | 35392 | **8697** | 426 |
| *Rattus norvegicus* | 21491 | **5270** | 2796 |
| *Bos taurus* | 8518 | **2220** | 1117 |
| *Caenorhabditis elegans* | 555 | **106** | 22 |
| *Drosophila melanogaster* | 2803 | **384** | 108 |
| *Danio rerio* | 509 | **99** | 29 |
| *Gallus gallus* | 815 | **140** | 32 |
| *Pongo abelii* | 318 | **34** | 2 |
| *Xenopus (Silurana) tropicalis* | 114 | **19** | 8 |
| *Sus scrofa* | 1230 | **277** | 126 |

**Supplementary Table 2 | Total number of L-R pairs at each threshold of combined STRING scores (TrEMBL)**

The predicted annotation of subcellular localization from TrEMBL was used.

| Organism | ≥150 | **≥400** | ≥700 |
|---|---|---|---|
| *Homo sapiens* | 1388 | **472** | 344 |
| *Mus musculus* | 1792 | **476** | 306 |
| *Arabidopsis thaliana* | 291 | **94** | 5 |
| *Rattus norvegicus* | 376 | **65** | 31 |
| *Bos taurus* | 634 | **237** | 178 |
| *Caenorhabditis elegans* | 8 | **1** | 0 |
| *Drosophila melanogaster* | 85 | **9** | 3 |
| *Danio rerio* | 1371 | **432** | 345 |
| *Gallus gallus* | 292 | **105** | 85 |
| *Pongo abelii* | 1079 | **184** | 16 |
| *Xenopus (Silurana) tropicalis* | 323 | **107** | 84 |
| *Sus scrofa* | 413 | **130** | 77 |

The coverage of the L-R databases, including the STRING-SWISSPROT/TrEMBL portions of LRBase.Hsa.eg.db (Hsa: *Homo sapiens*), FANTOM5, IUPHAR, and DLRP are summarized in Figure S2. Although FANTOM5 integrated IUPHAR and DLRP, the set difference is large, and only 32% of L-R pairs from FANTOM5 are covered by IUPHAR and DLRP (Figure S2, SWISSPROT × STRING, ≥400). This might be because FANTOM5 also merged HPMR and HPRD, which are no longer accessible databases. Using LRBase.Hsa.eg.db, about 56% of L-R pairs from FANTOM5 are covered (Figure S2, SWISSPROT × STRING; ≥400).

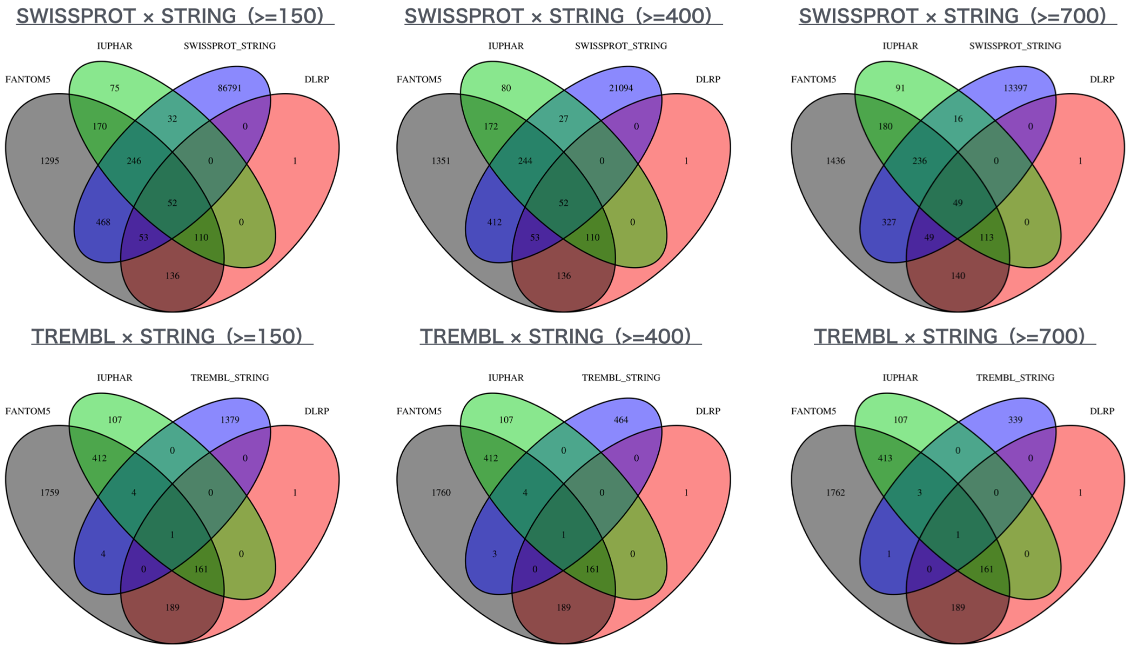


**Supplementary Figure 2 | Coverage of STRING× SWISSPROT/TrEMBL data with different combined STRING scores**

The L-R pairs from FANTOM5, IUPHAR, and DLRP are also compared.

Contrary to the fact that FANTOM5 data are not maintained and HPMR and HPRD are inaccessible, LRBase.XXX.eg.db is sustainable in the context of the changing knowledge regarding subcellular localization and PPIs. Compared with the 2014 data, when the FANTOM5 paper was published, available TrEMBL sequence entries and STRING PPIs have grown by $(1.07 \times 10^8)/(5.16 \times 10^7) \approx 2.07$ fold and $(1.38 \times 10^9)/(3.32 \times 10^8) \approx 4.15$ fold, respectively (Figure S3). Other differences between our workflow and that of FANTOM5 are summarized in Table S4.

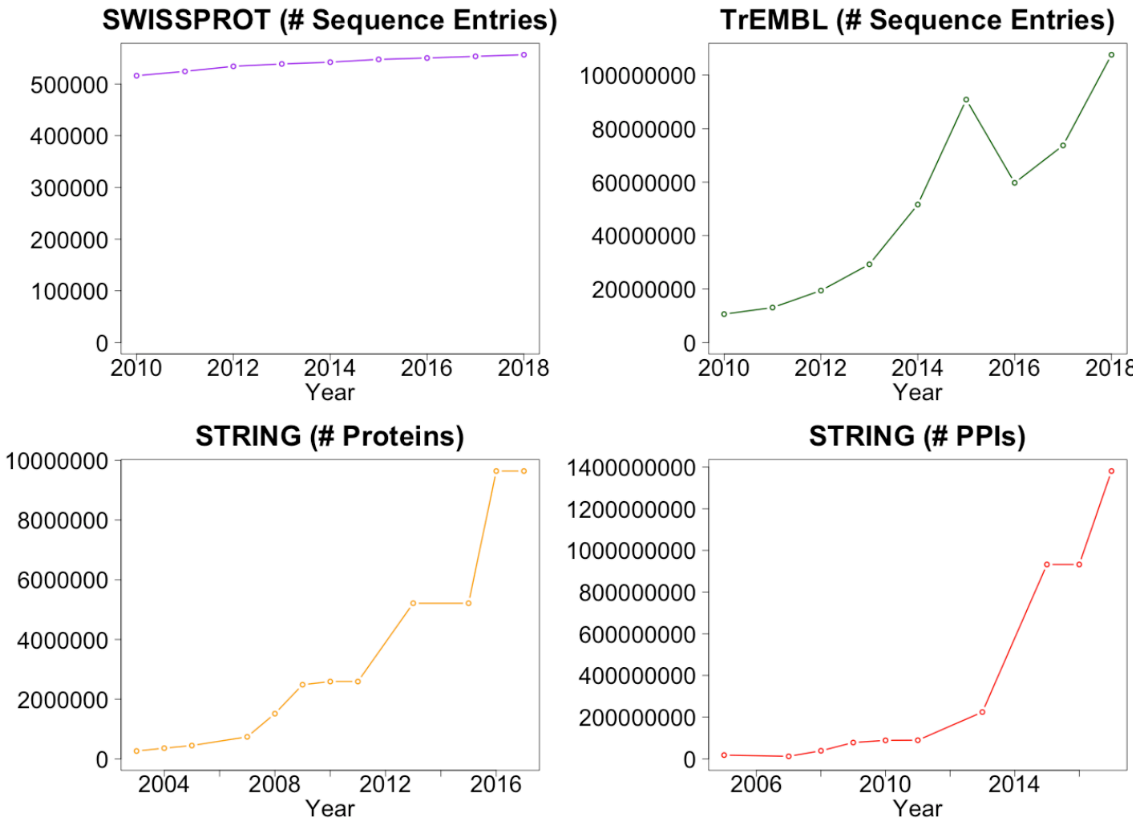


**Supplementary Figure 3 | Changes in SWISSPROT, TrEMBL, and STRING entries over time**

The number of sequence entries for SWISSPROT (purple) / TrEMBL (green) and the number of proteins (orange) and PPIs (red) in STRING are summarized for each year.

**Supplementary Table 4 | Differences in data preprocessing workflow between FANTOM5 and LRBase.XXX.eg.db**

| | FANTOM5 | LRBase.XXX.eg.db | Reason |
|---|---|---|---|
| Difference of Ontology | PM: Plasma Membrane | Cellular Membrane | It is discontinued in UniProtKB |
| Inaccessible database | HPMR and HPRD | No use | The corresponding information cannot be found |
| Human specific database | HPRD, DLRP, IUPHAR, and HPMR | No use | Such a database cannot be applied to other species |
| Identifier of Gene | Gene Name (HGNC) | NCBI Gene ID (Entrez Gene ID) | For uniqueness and consistency of analytical pipelines combined with other Bioconductor packages |
| Prediction of subcellular localization | PolyPhobius and LocTree3 | TMHMM, SignalP, Phobius, and Coils (TrEMBL) | The corresponding information is still predicted by TrEMBL |
| The threshold of PPI pairs of STRING | Over 700 confidence score at "physical-binding interactions" and "experimental interactions" | Over 400 confidence score at "combined" | For usage with a wide range of organisms |
| Organisms | Only Human | 12 organisms | For usage with a wide range of organisms |
| Repository | FANTOM5 web site | Bioconductor | For sustainable maintenance |

On the other hand, we also found that some well-known L-R pairs, such as Delta-Notch, are not included by the approach using STRING×

SWISSPROT/TrEMBL alone. Therefore, only in the case of LRBase.Hsa.eg.db, we also integrated L-R pairs from IUPHAR and DLRP. We compared the coverage of the L-R list from the final version of LRBase.Hsa.eg.db described above with that of FANTOM5 (Figure S4). We also compared LRBase.Hsa.eg.db with recent human L-R databases such as CellPhoneDB (https://www.cellphonedb.org) created by the Teichmann Lab of the Wellcome Sanger Institute and Cell-Cell Interaction Database created by the Bader Laboratory at the University of Toronto (BaderLab, http://baderlab.org/CellCellInteractions).

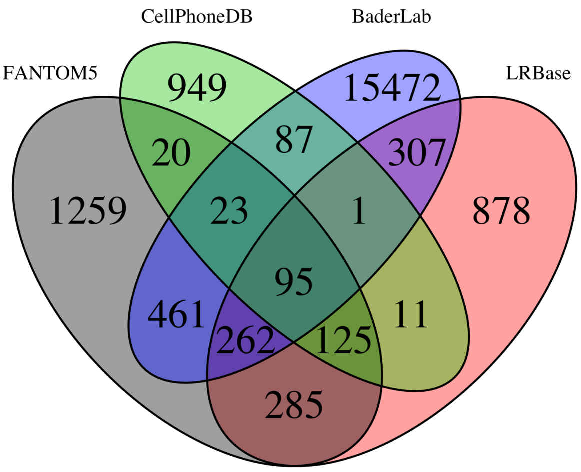


**Supplementary Figure 4 | Coverage of the STRING× SWISSPROT/TrEMBL merged with IUPHAR and DLRP**

The L-R pairs from FANTOM5, CellPhoneDB, and BaderLab are also compared.

The Venn diagram shows that the coverage of these databases is small. This may be caused by the individual policies of each database, such as the using PPI databases and the L-R pairing methods. For example, CellPhoneDB also introduced the FANTOM5-like approach with its use of PPI×Secreted/Membrane annotation, but the database also uses protein complex annotation from PDB and literature mining, which does not have a one-to-one relationship. The Bader Lab uses some PPI databases, but the pairing is not restricted to L-R relationships; rather, it is extended to all the combinations of ligand, receptor, and extracellular matrix elements as annotated by Gene Ontology. Under the condition that there are few correct CCIs and related L-R pairs, it is difficult to determine which approach is best; the approach containing many genes may cause false-positive pairs. We intend to perform benchmarking with exhaustive CCI datasets in the future.
