## Supplementary figures and images for "Uncovering hypergraphs of cell-cell interaction from single cell RNA-sequencing data"

### SWISSPROT_STRING_Ath_PAIRPLOT.png

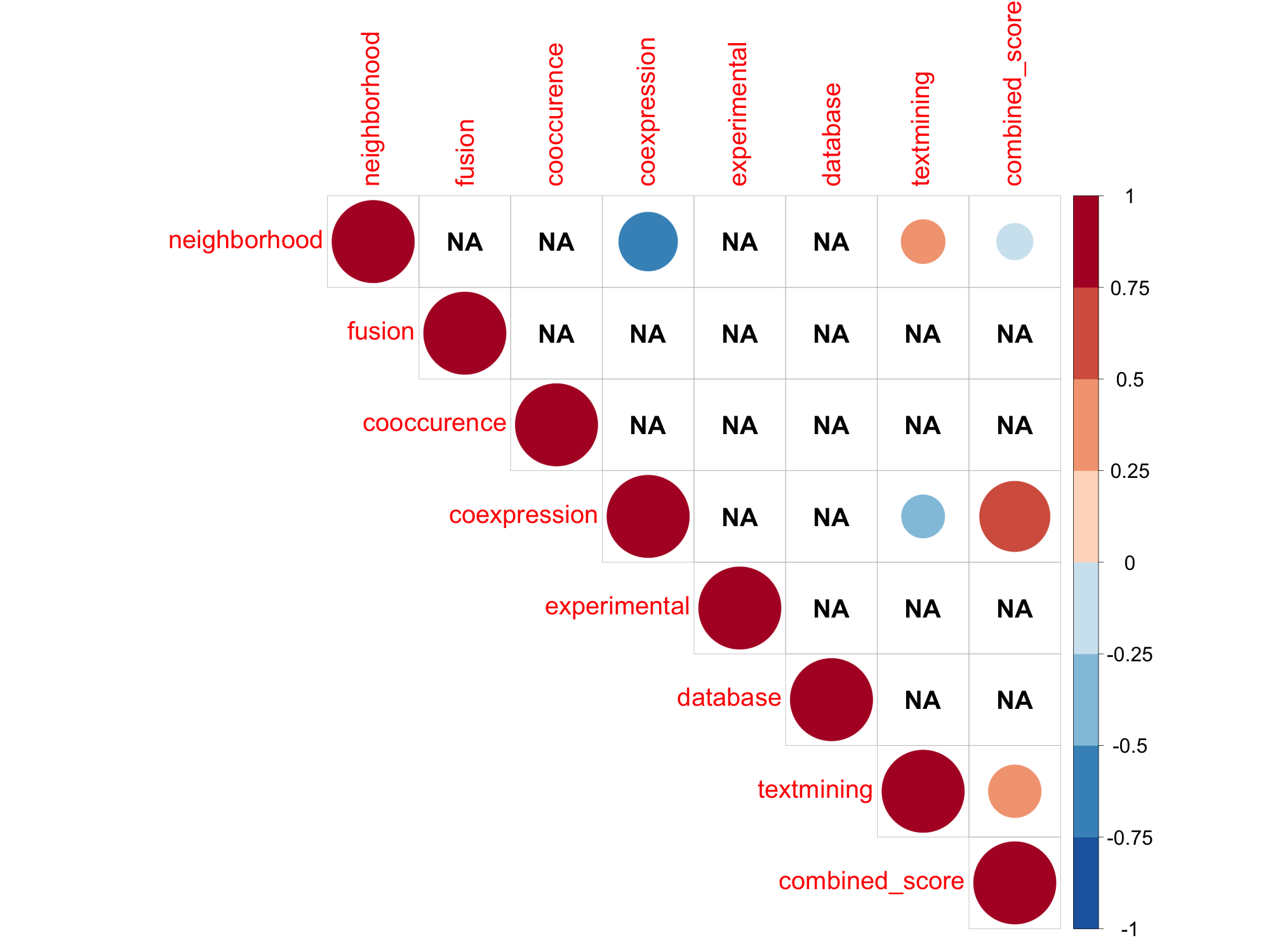

### SWISSPROT_STRING_Ath_TABLEPLOT.png

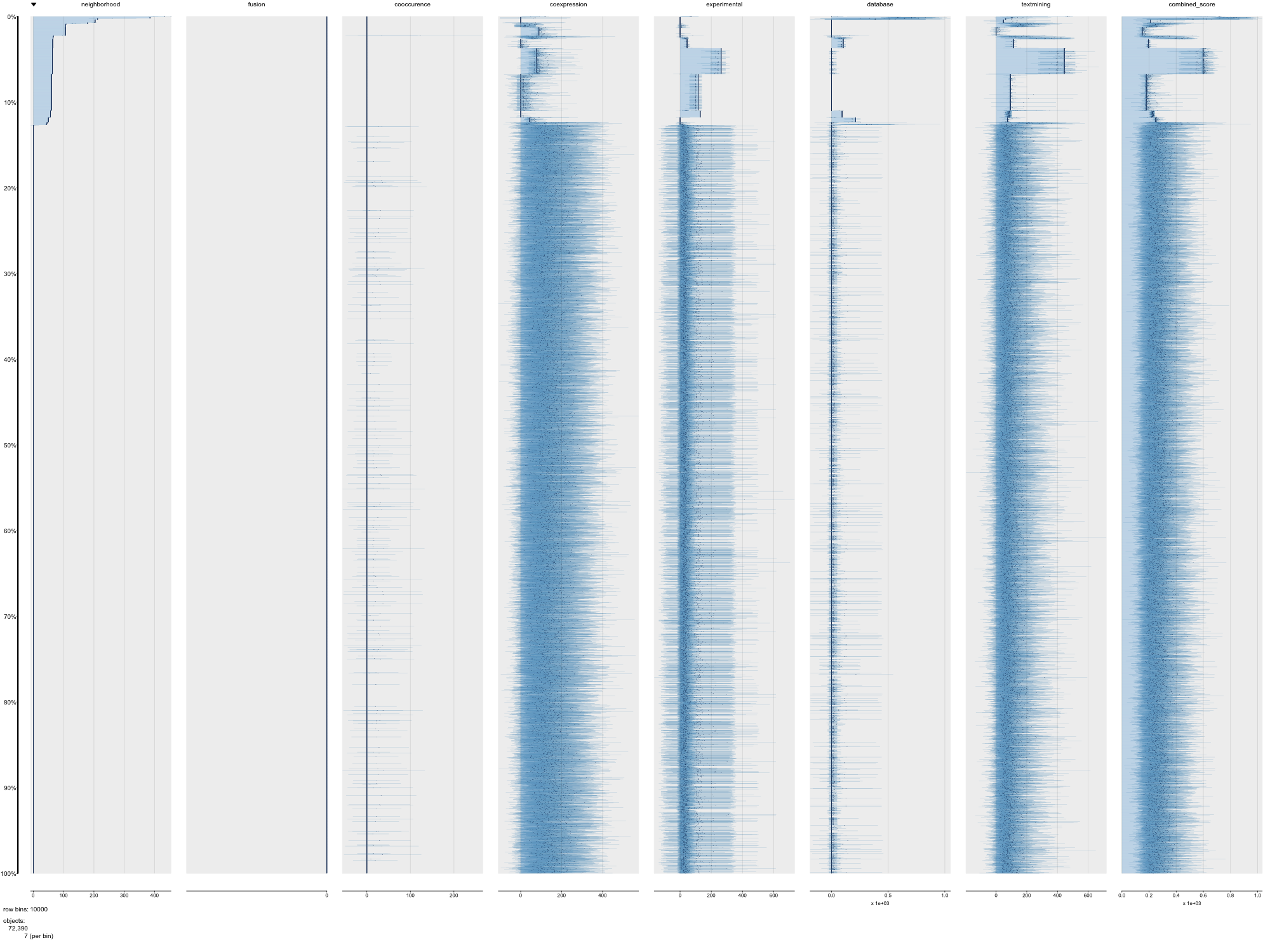

### SWISSPROT_STRING_Bta_PAIRPLOT.png

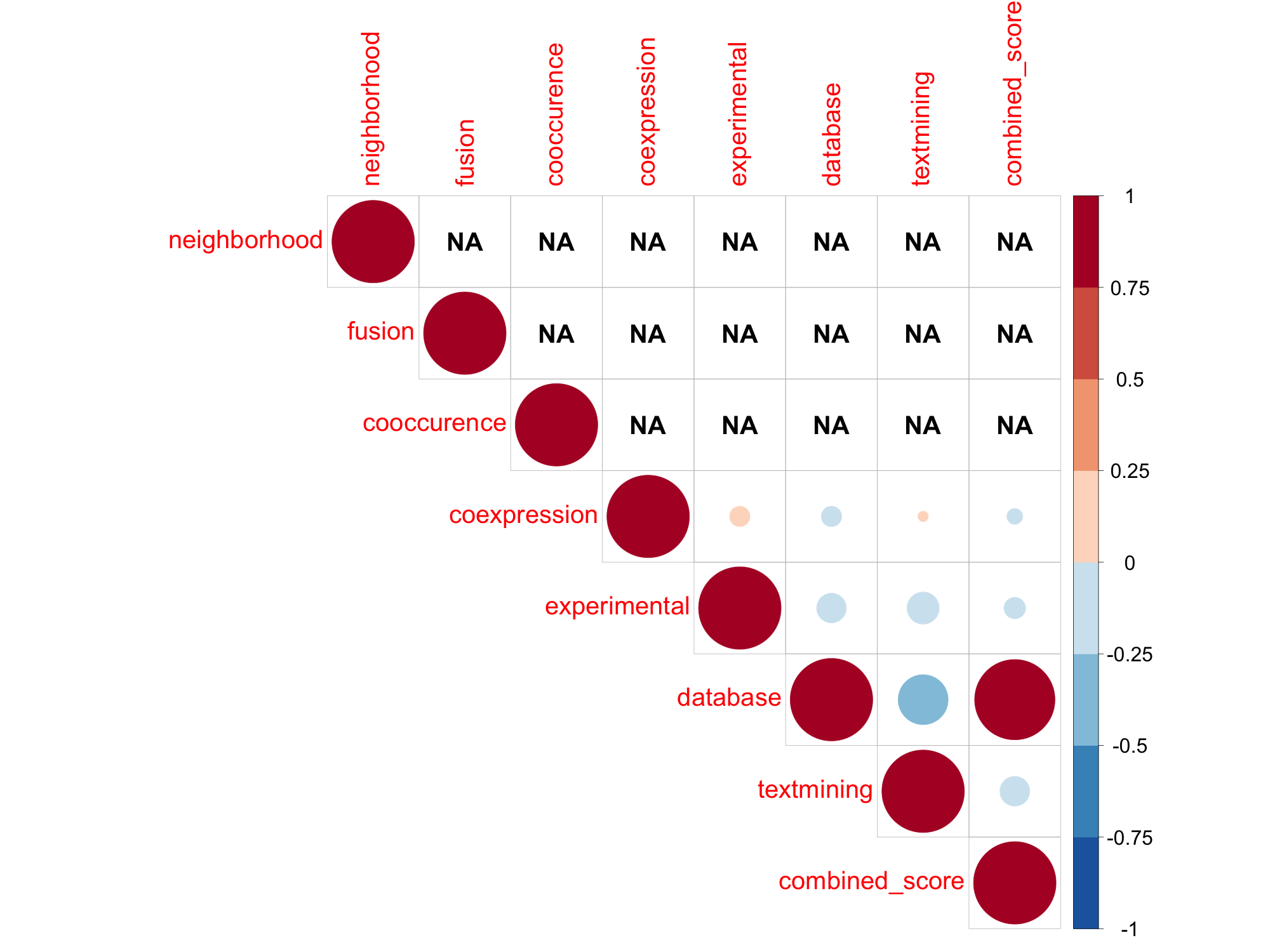

### SWISSPROT_STRING_Bta_TABLEPLOT.png

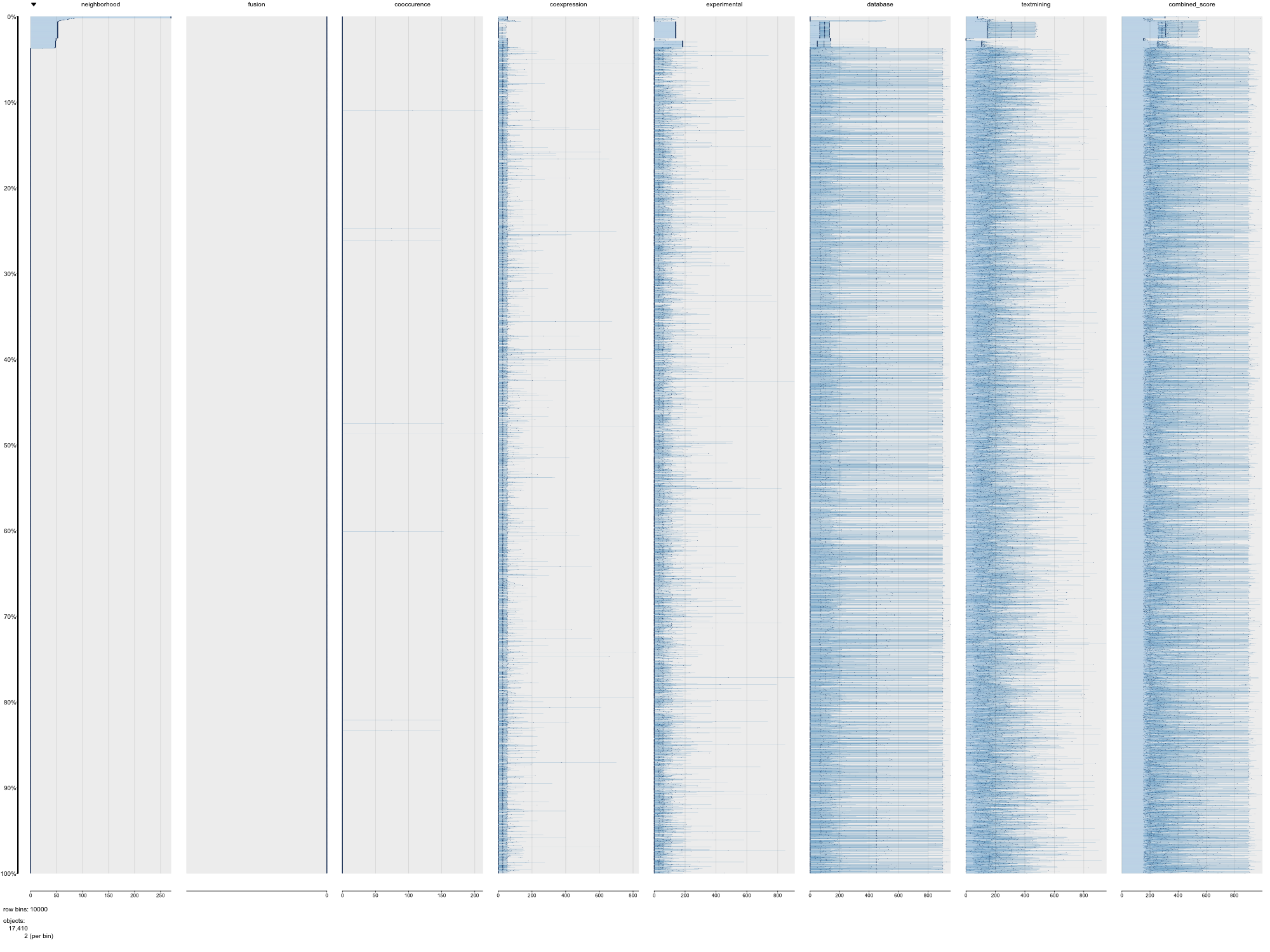

### SWISSPROT_STRING_Cel_PAIRPLOT.png

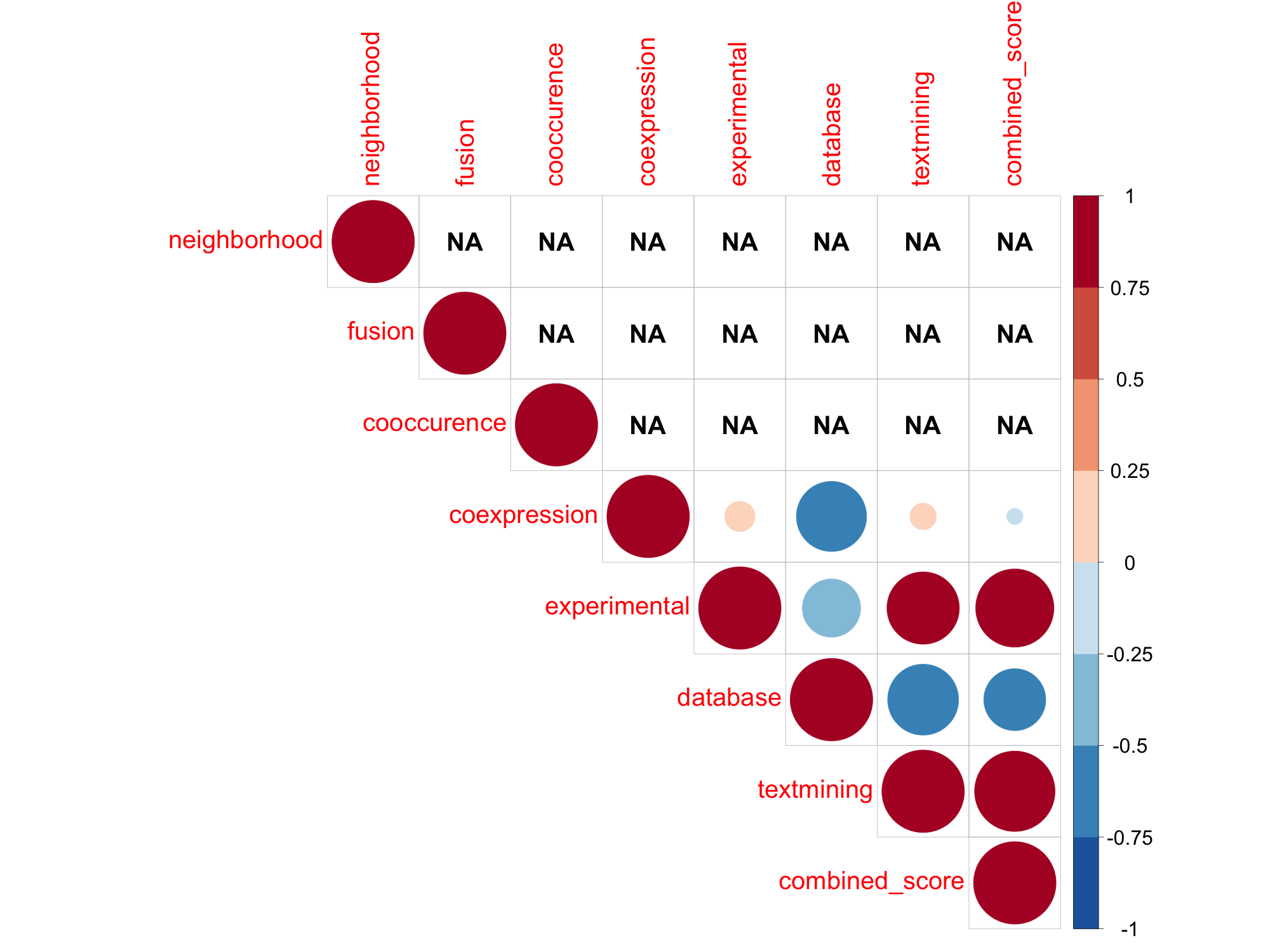

### SWISSPROT_STRING_Cel_TABLEPLOT.png

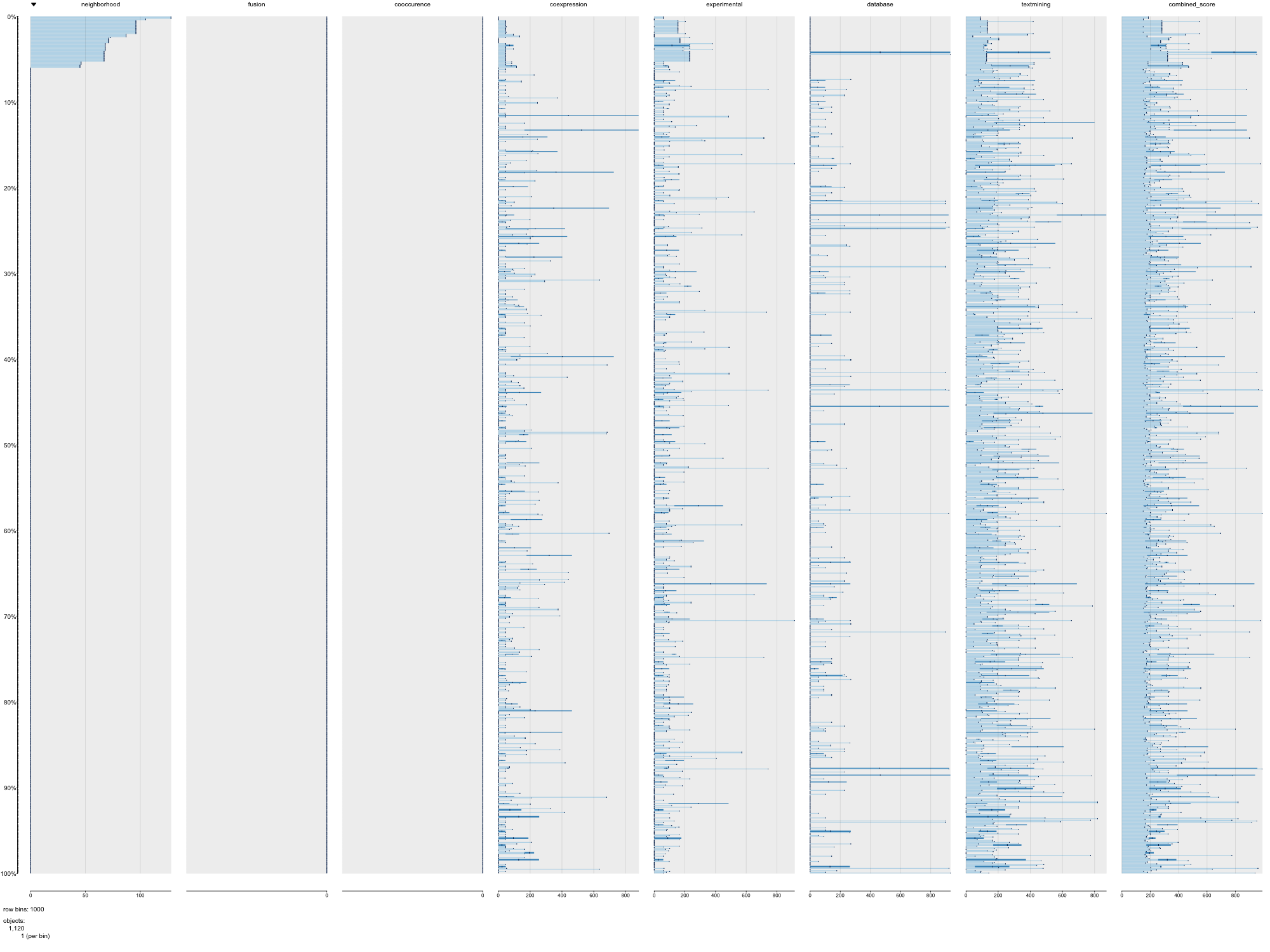

### SWISSPROT_STRING_Dme_PAIRPLOT.png

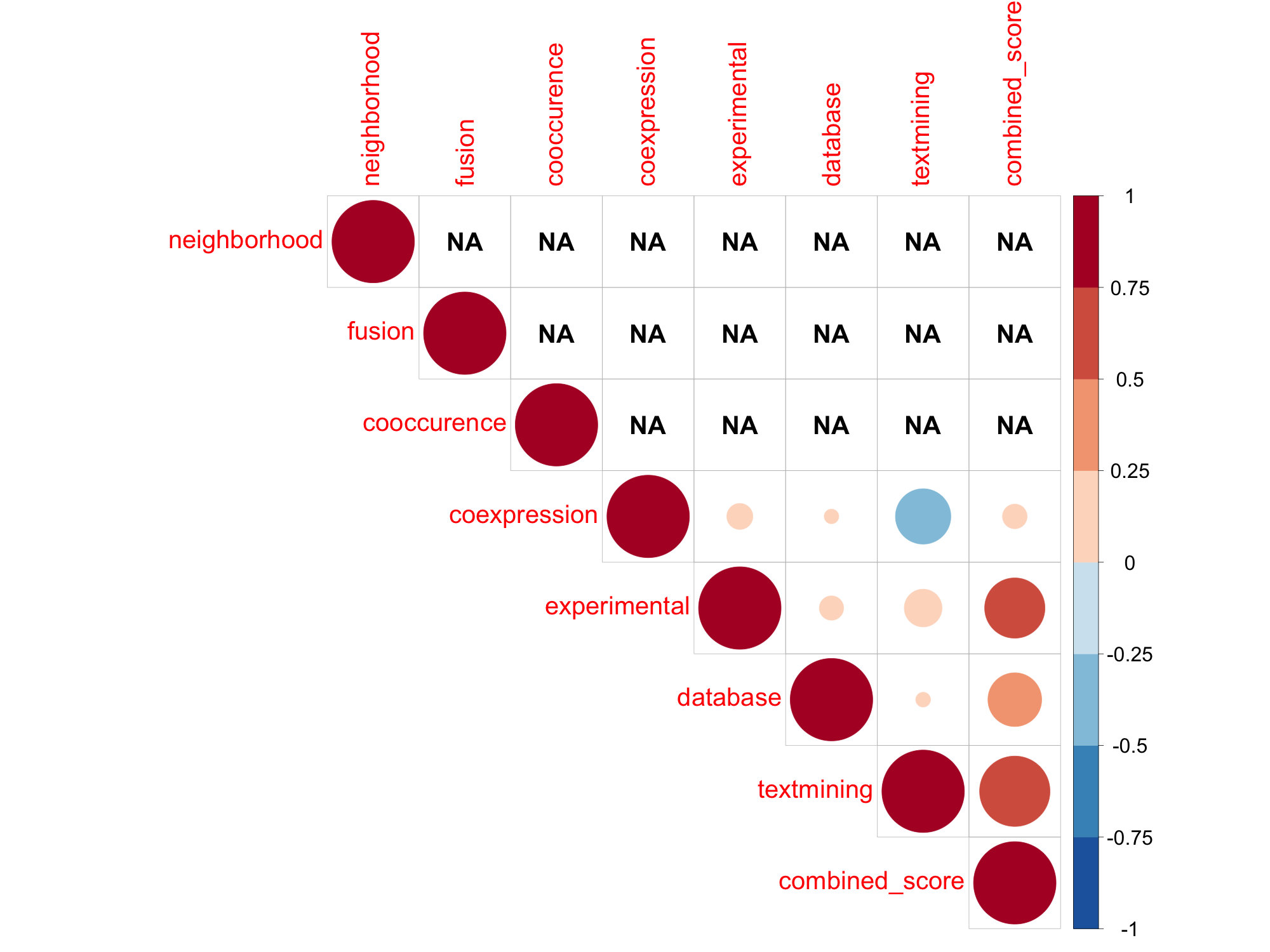

### SWISSPROT_STRING_Dme_TABLEPLOT.png

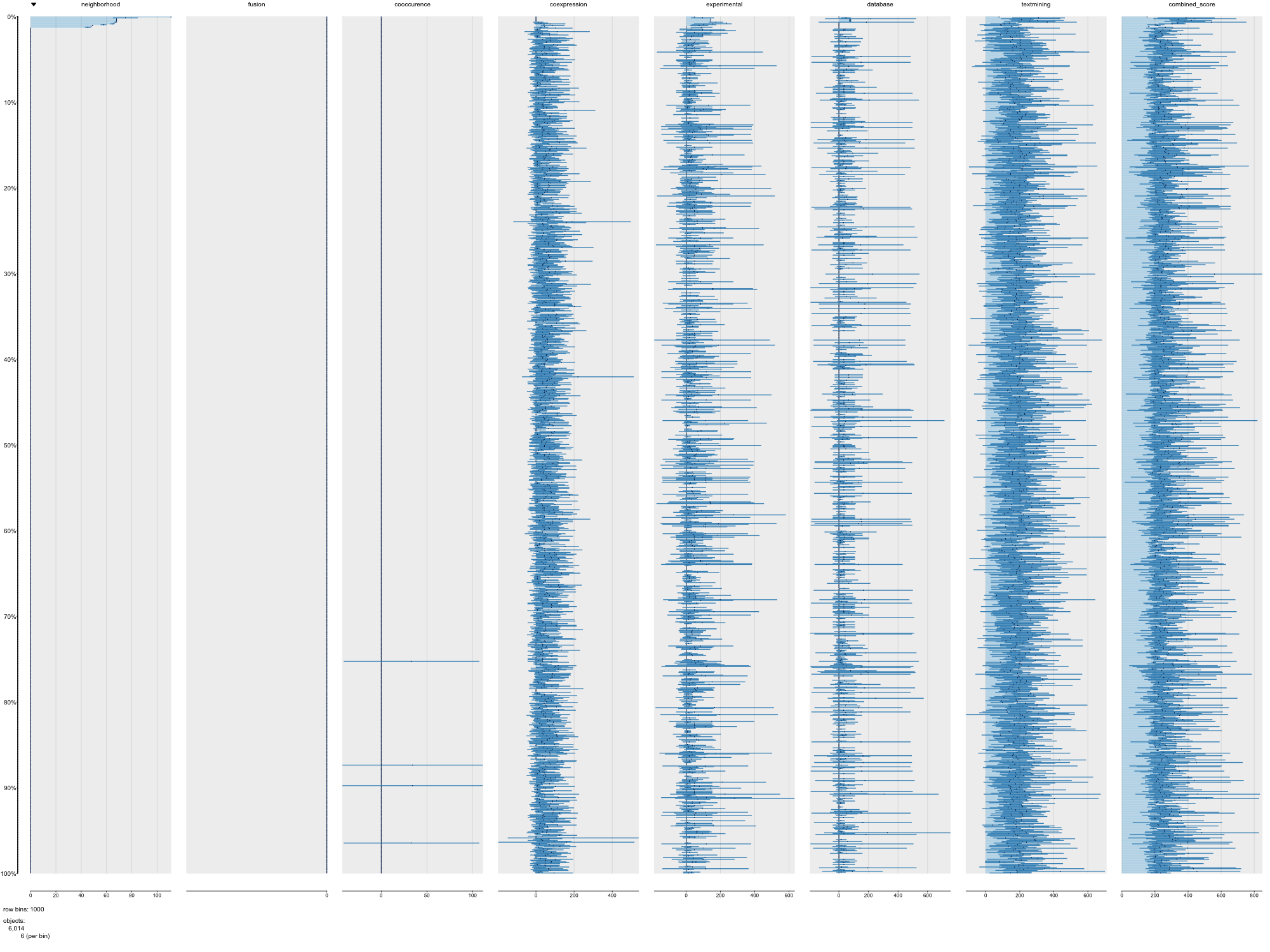

### SWISSPROT_STRING_Dre_PAIRPLOT.png

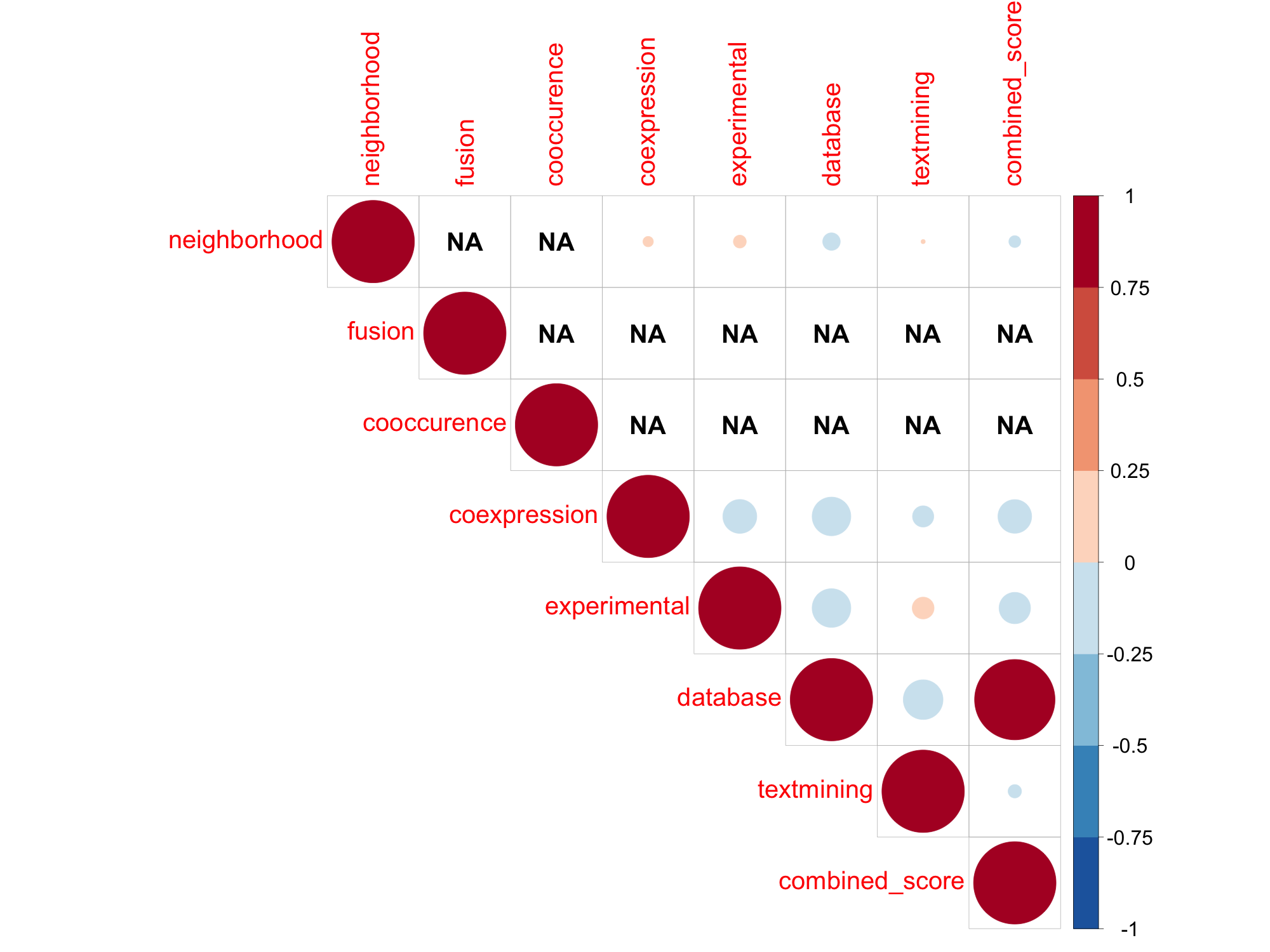

### SWISSPROT_STRING_Dre_TABLEPLOT.png

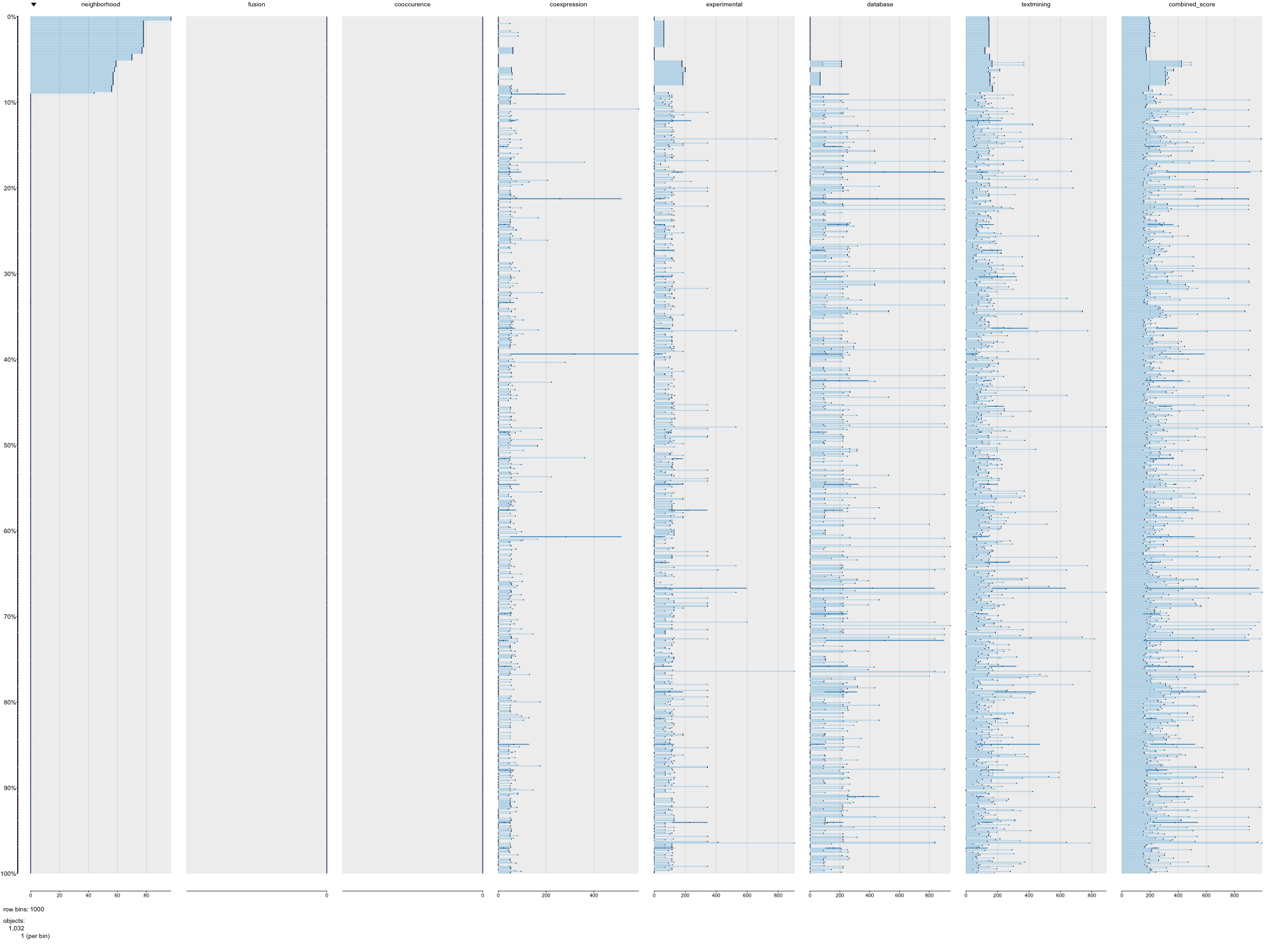

### SWISSPROT_STRING_Gga_PAIRPLOT.png

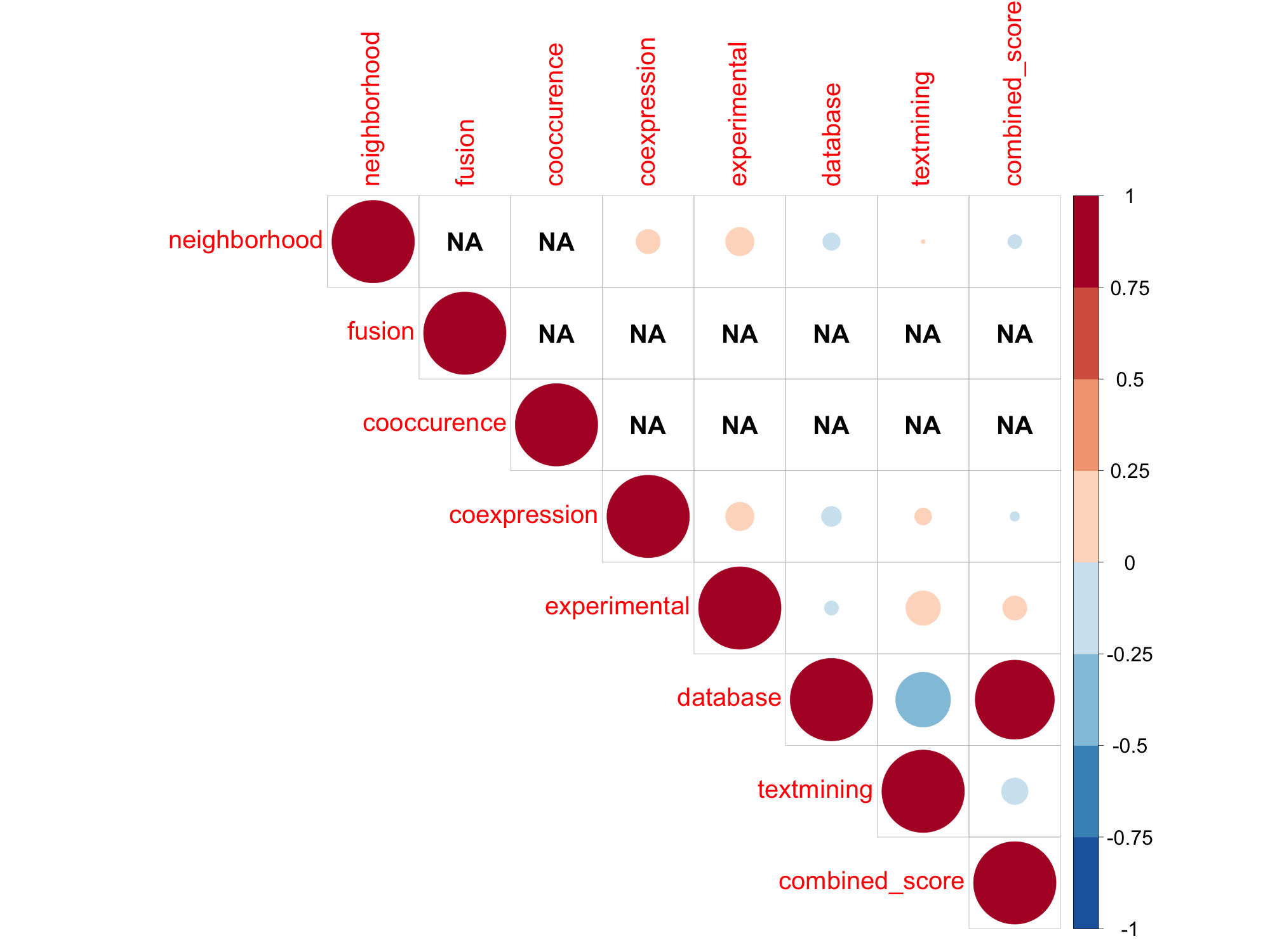

### SWISSPROT_STRING_Gga_TABLEPLOT.png

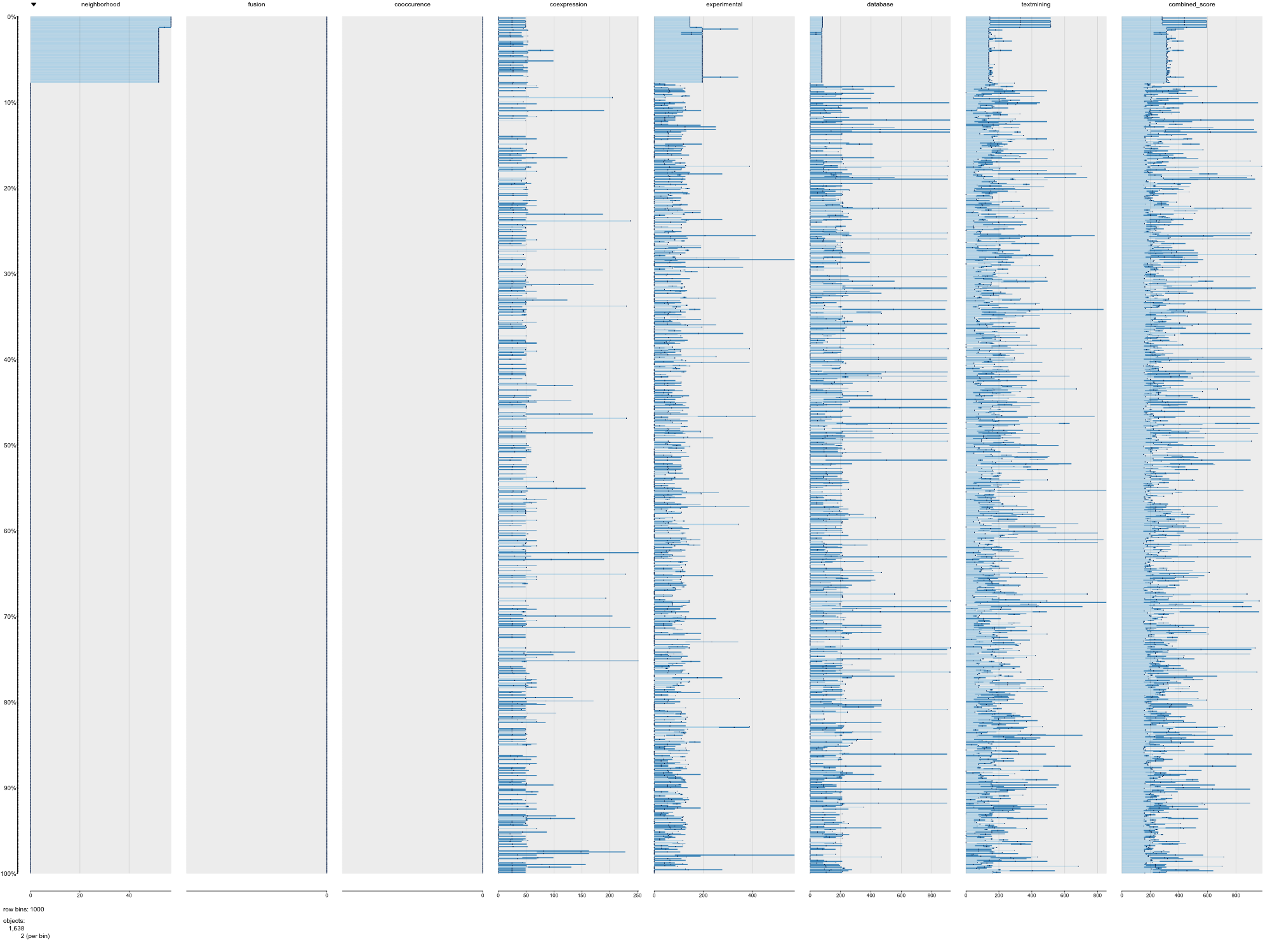

### SWISSPROT_STRING_Hsa_PAIRPLOT.png

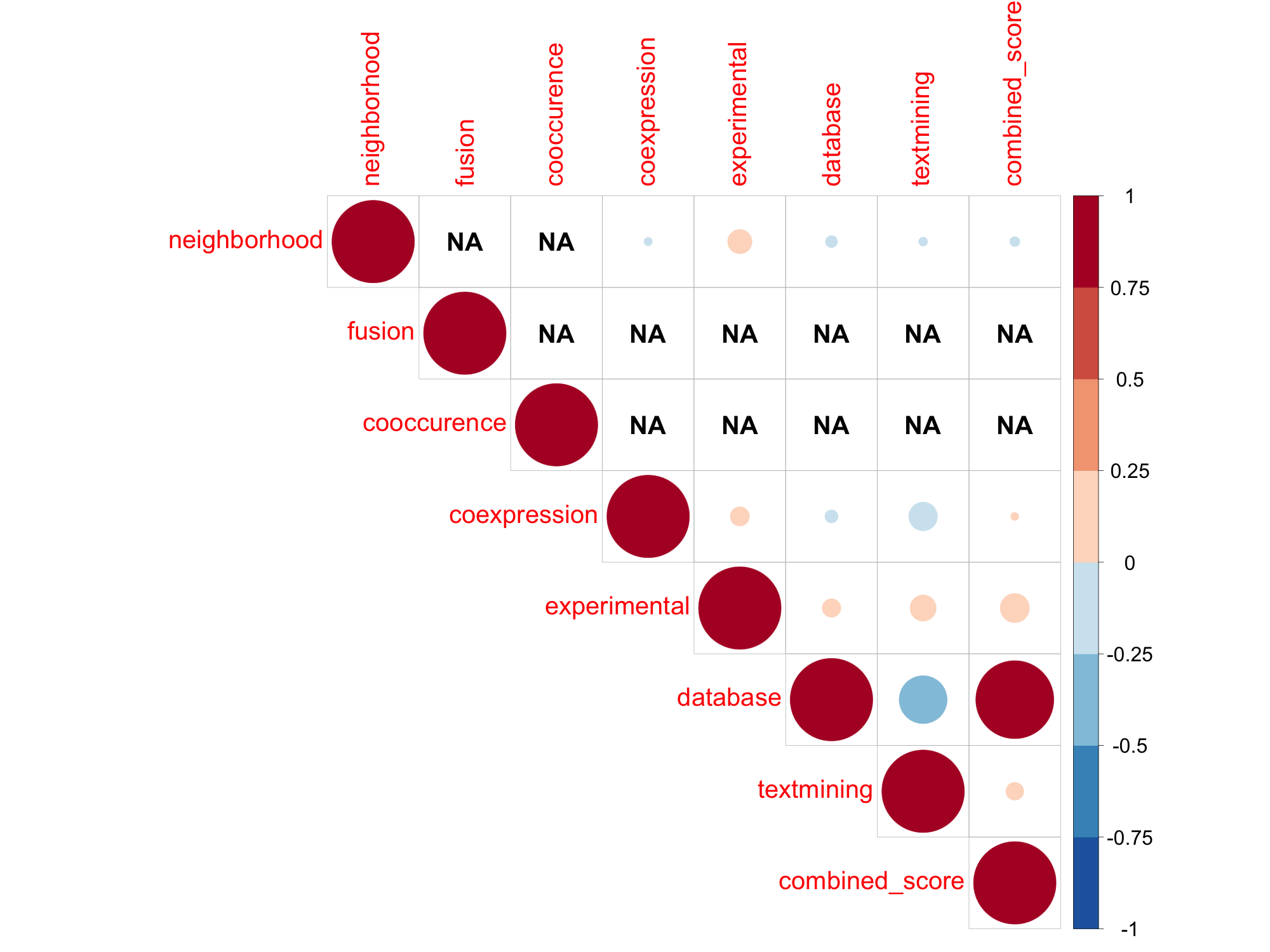

### SWISSPROT_STRING_Hsa_TABLEPLOT.png

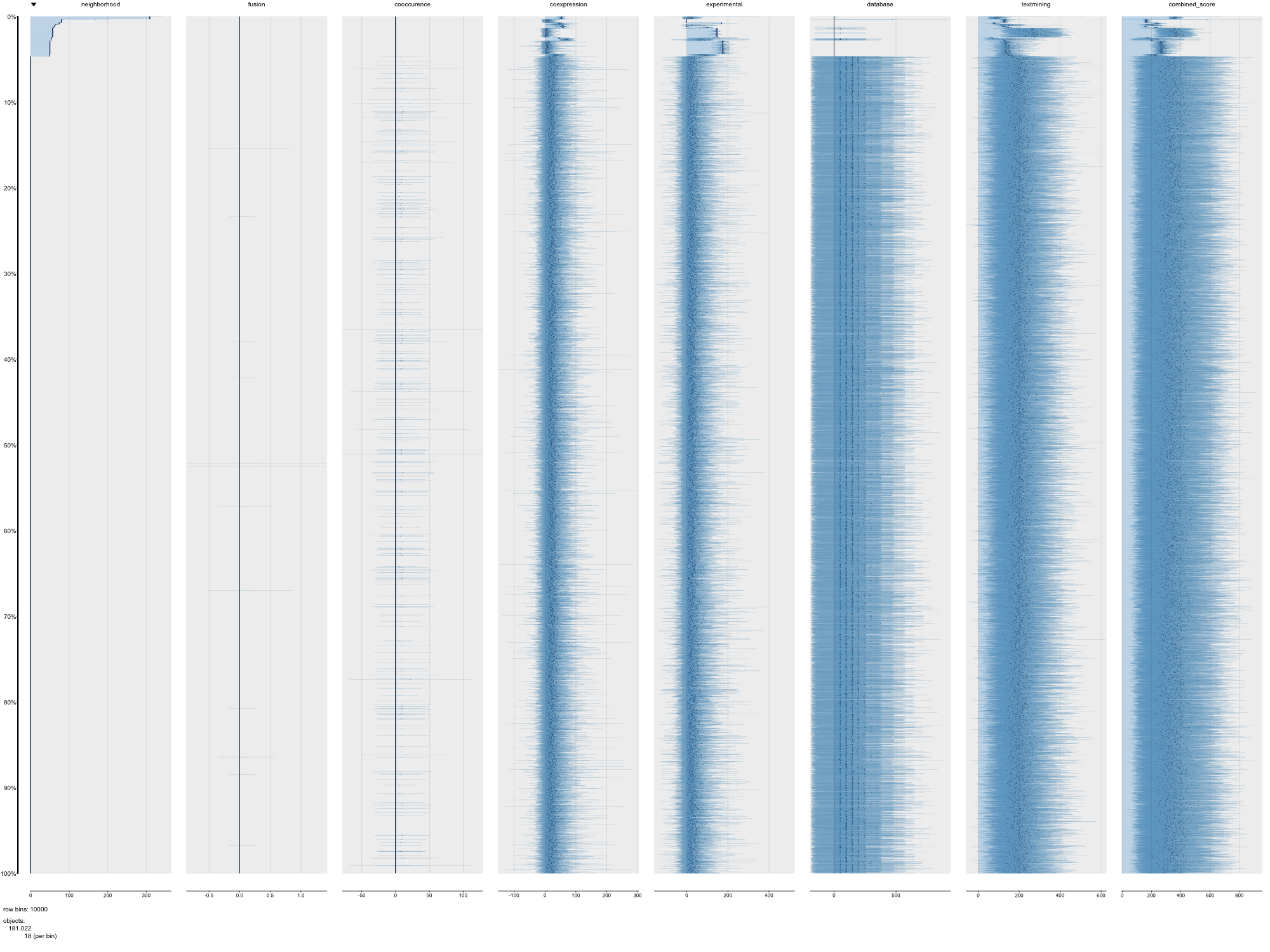

### SWISSPROT_STRING_Mmu_PAIRPLOT.png

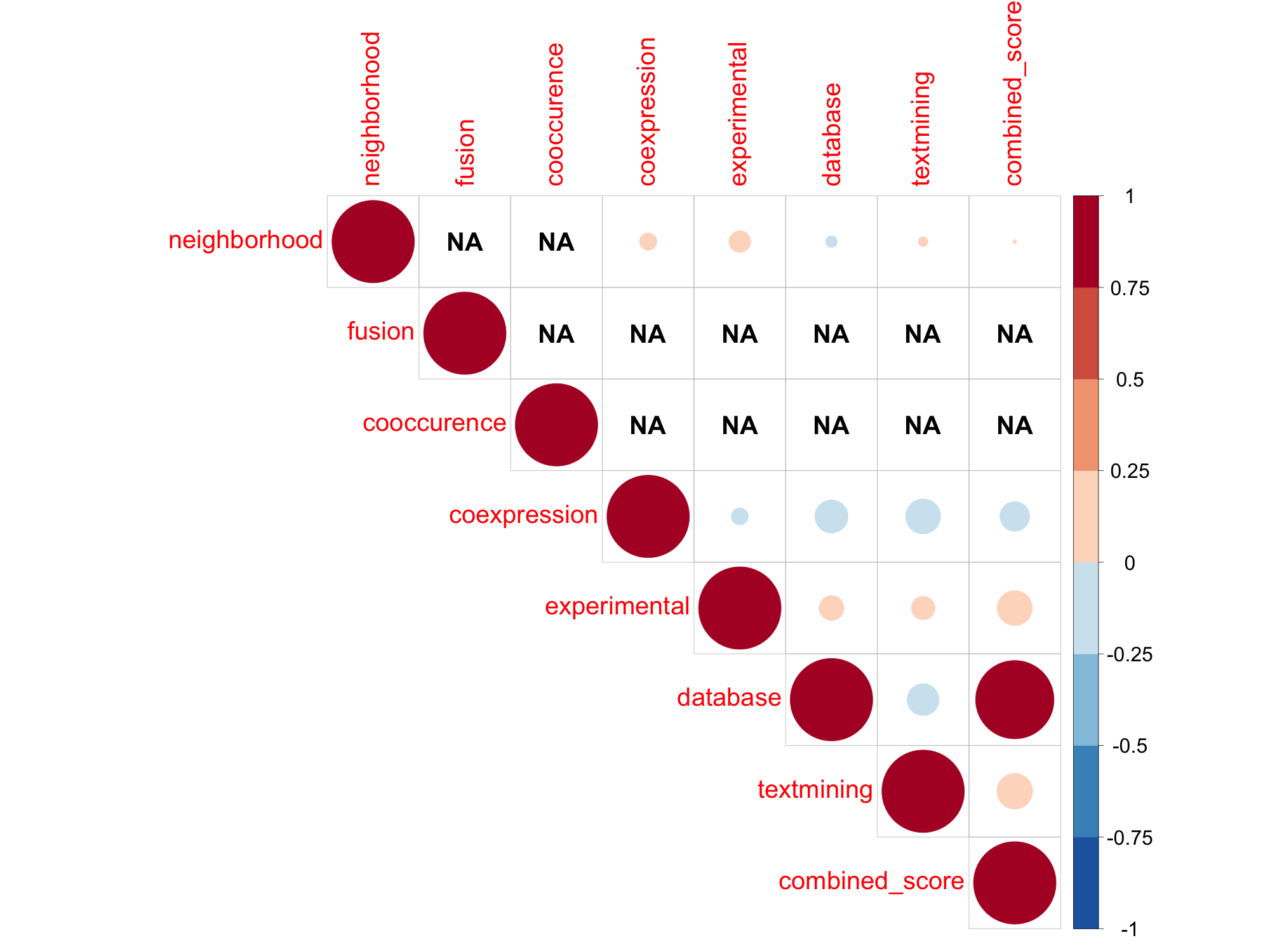

### SWISSPROT_STRING_Mmu_TABLEPLOT.png

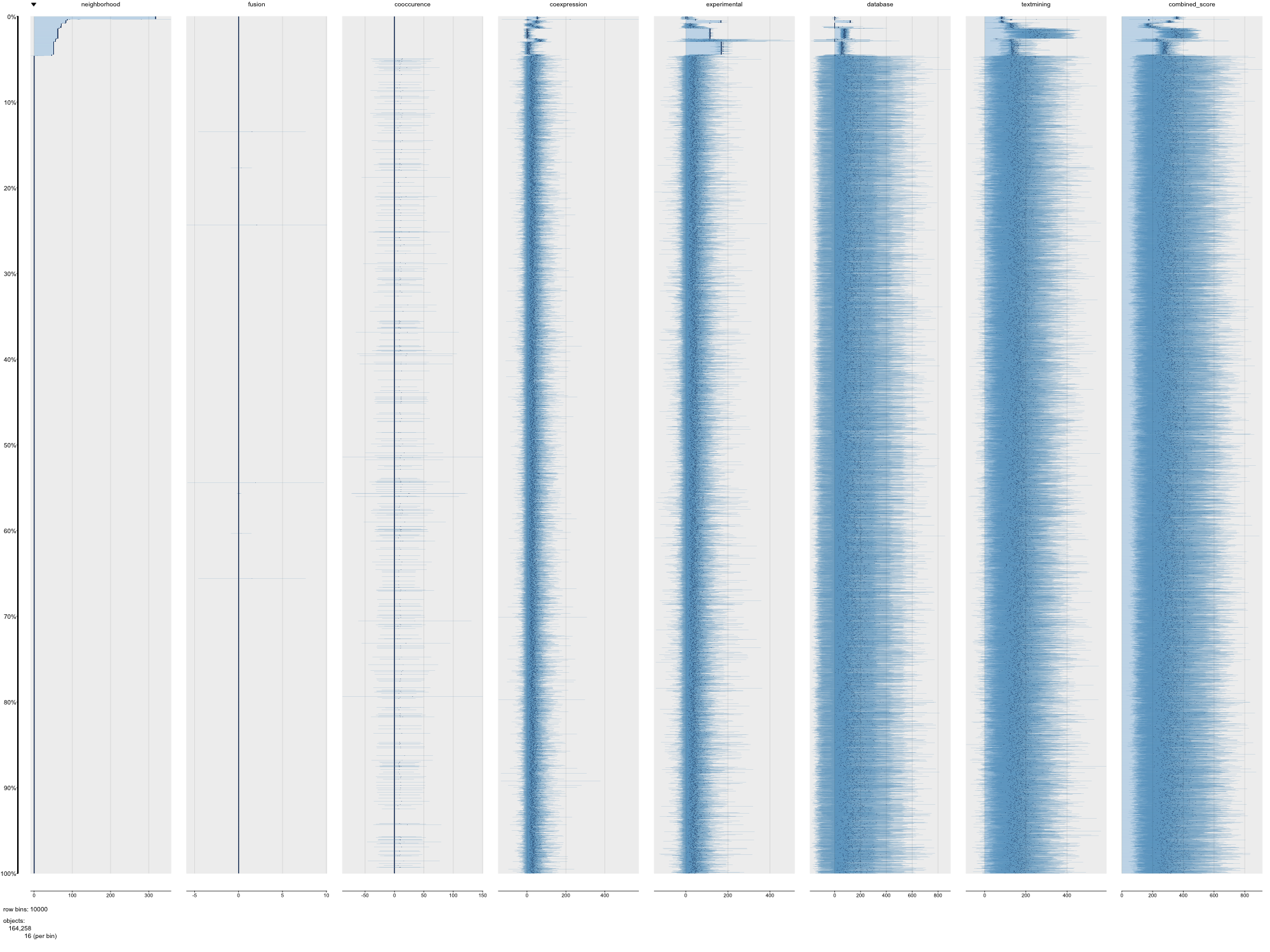

### SWISSPROT_STRING_Pab_PAIRPLOT.png

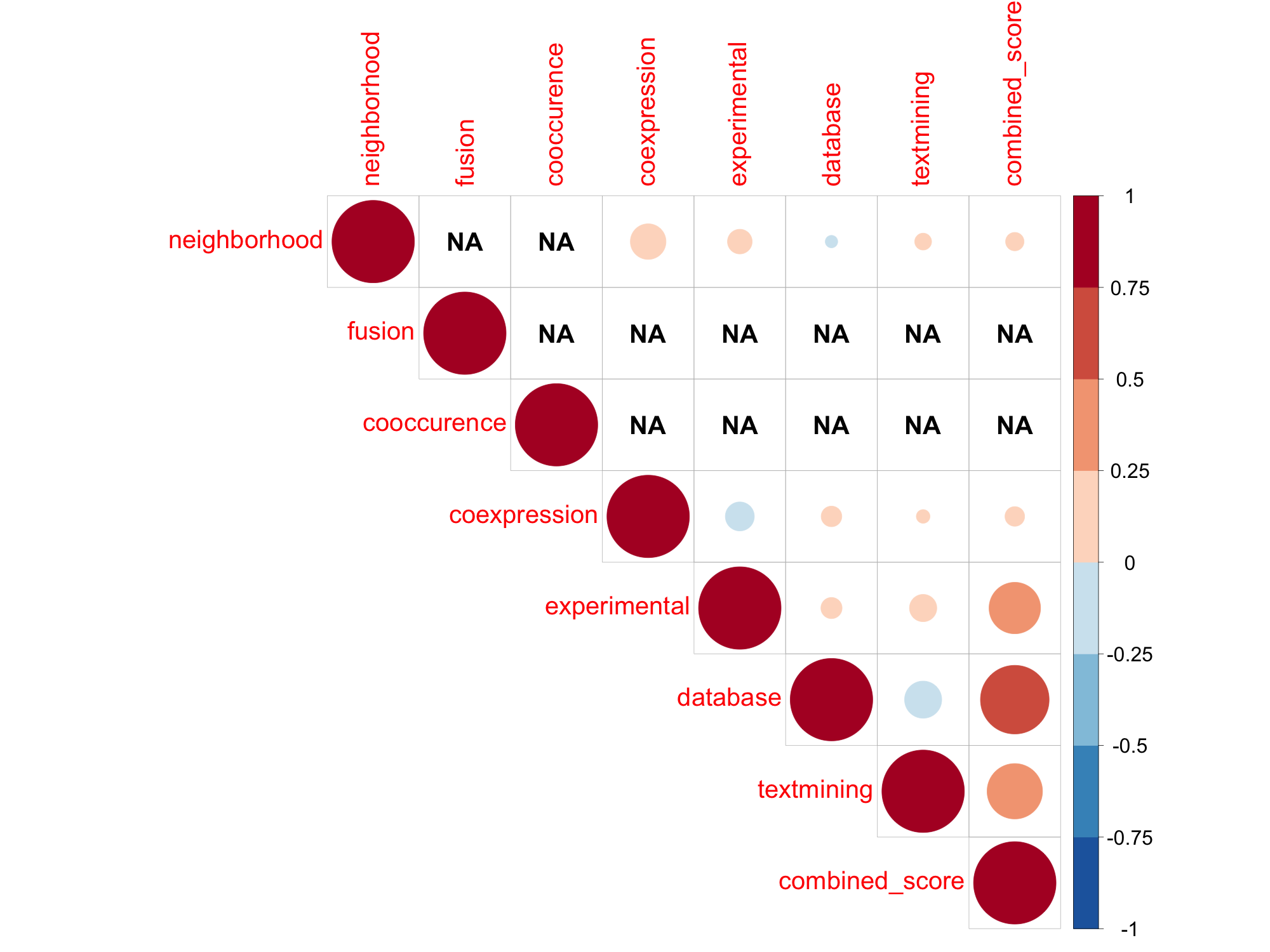

### SWISSPROT_STRING_Pab_TABLEPLOT.png

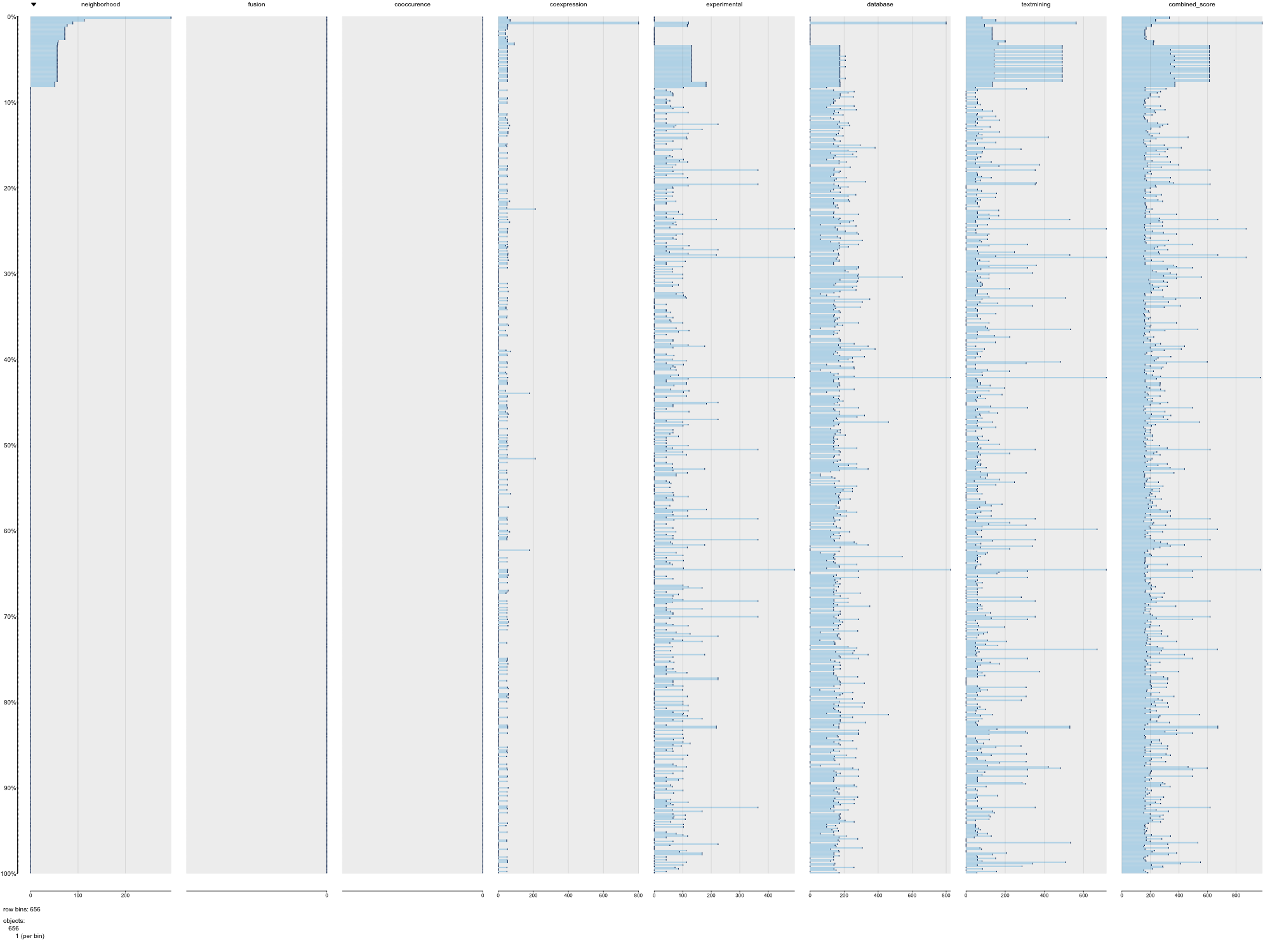

### SWISSPROT_STRING_Rno_PAIRPLOT.png

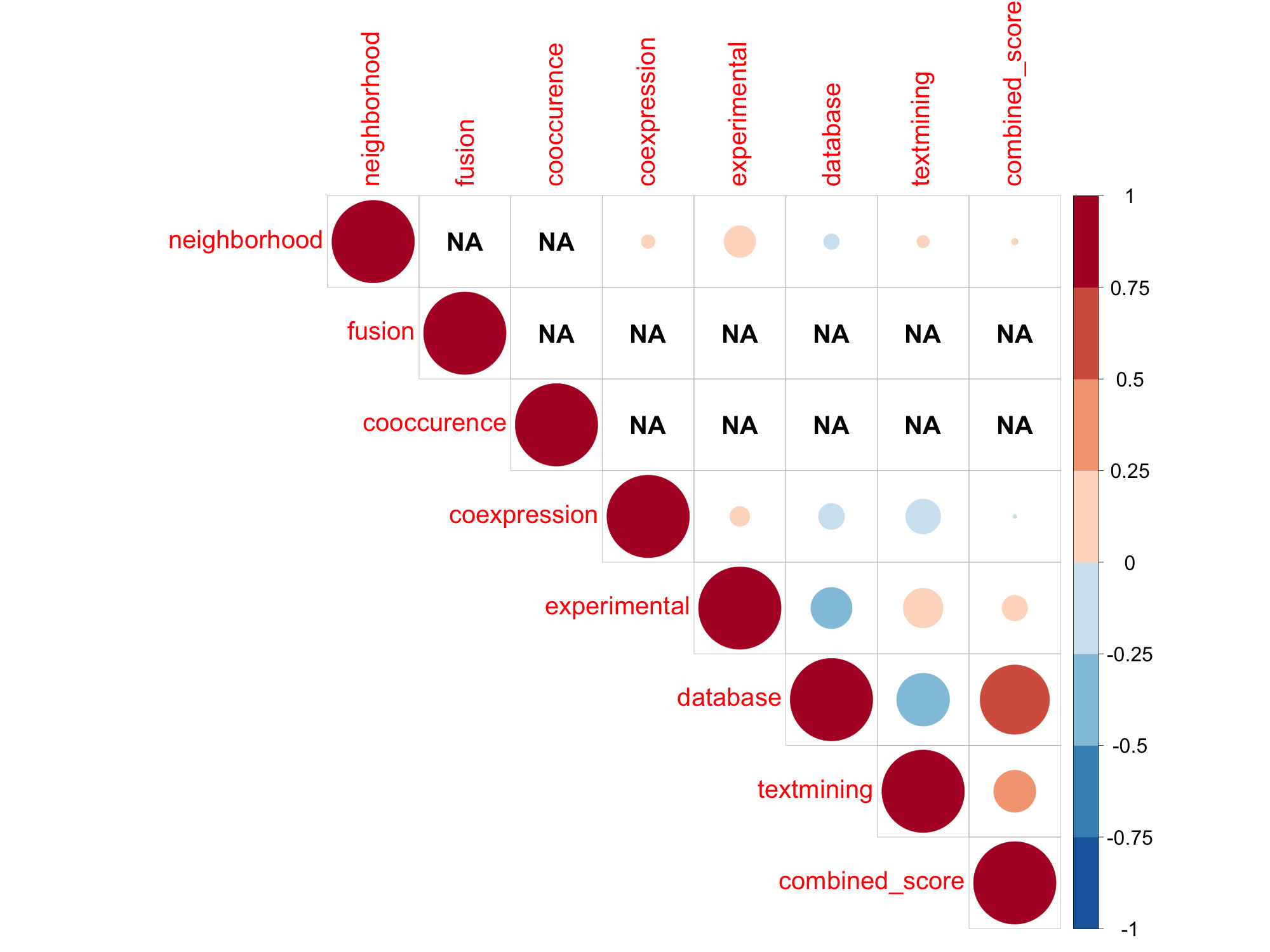

### SWISSPROT_STRING_Rno_TABLEPLOT.png

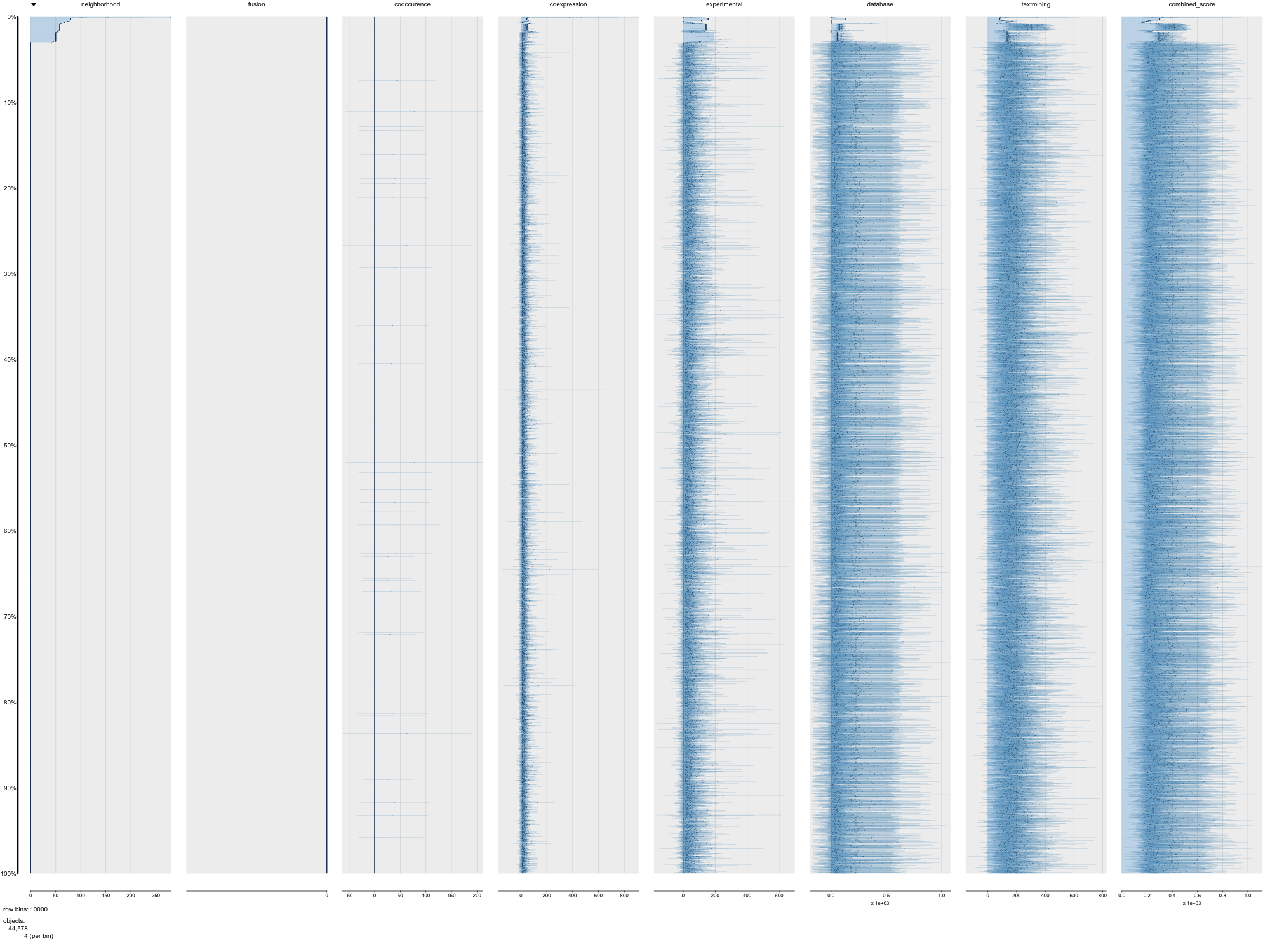

### SWISSPROT_STRING_Ssc_PAIRPLOT.png

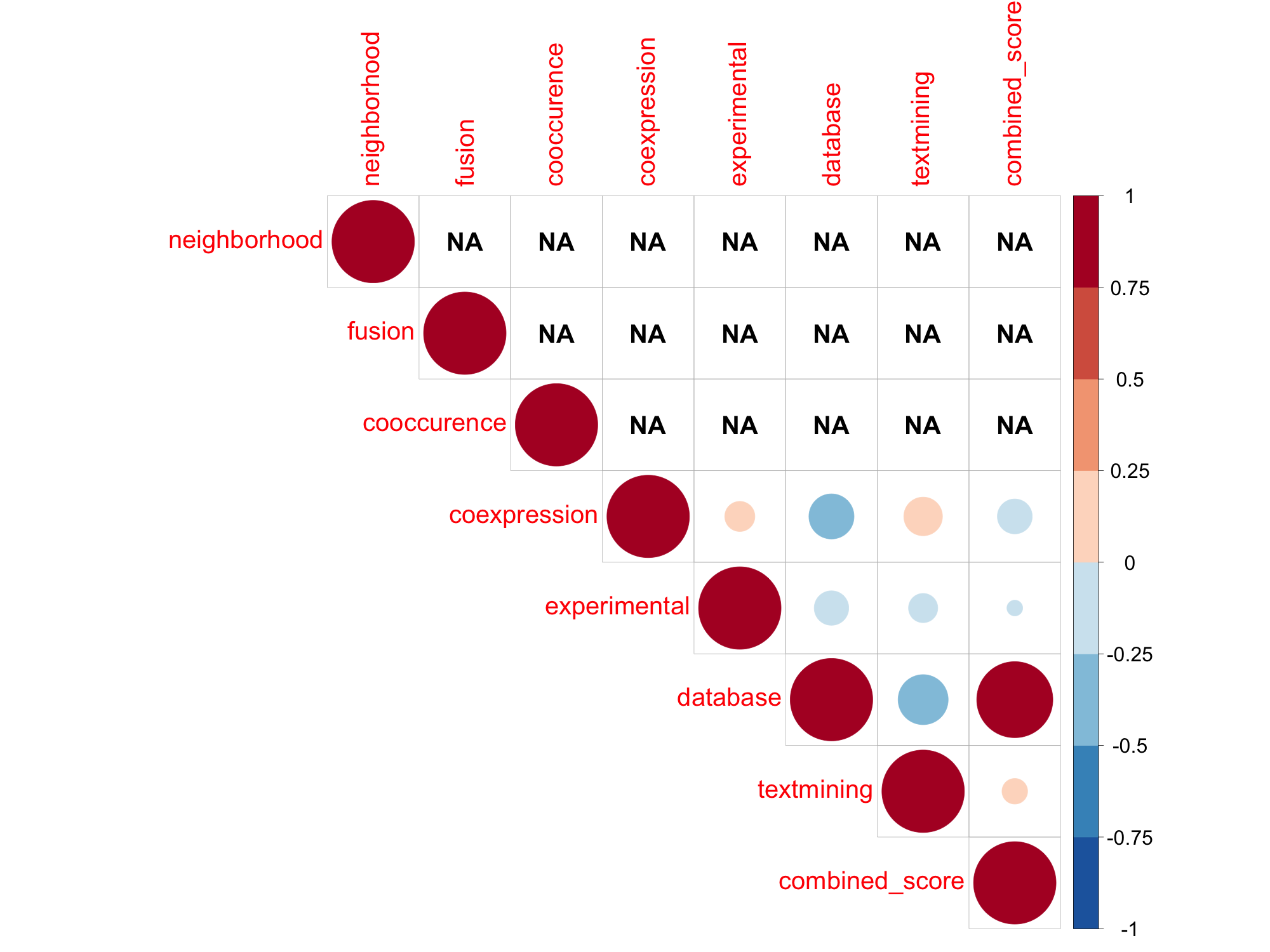

### SWISSPROT_STRING_Ssc_TABLEPLOT.png

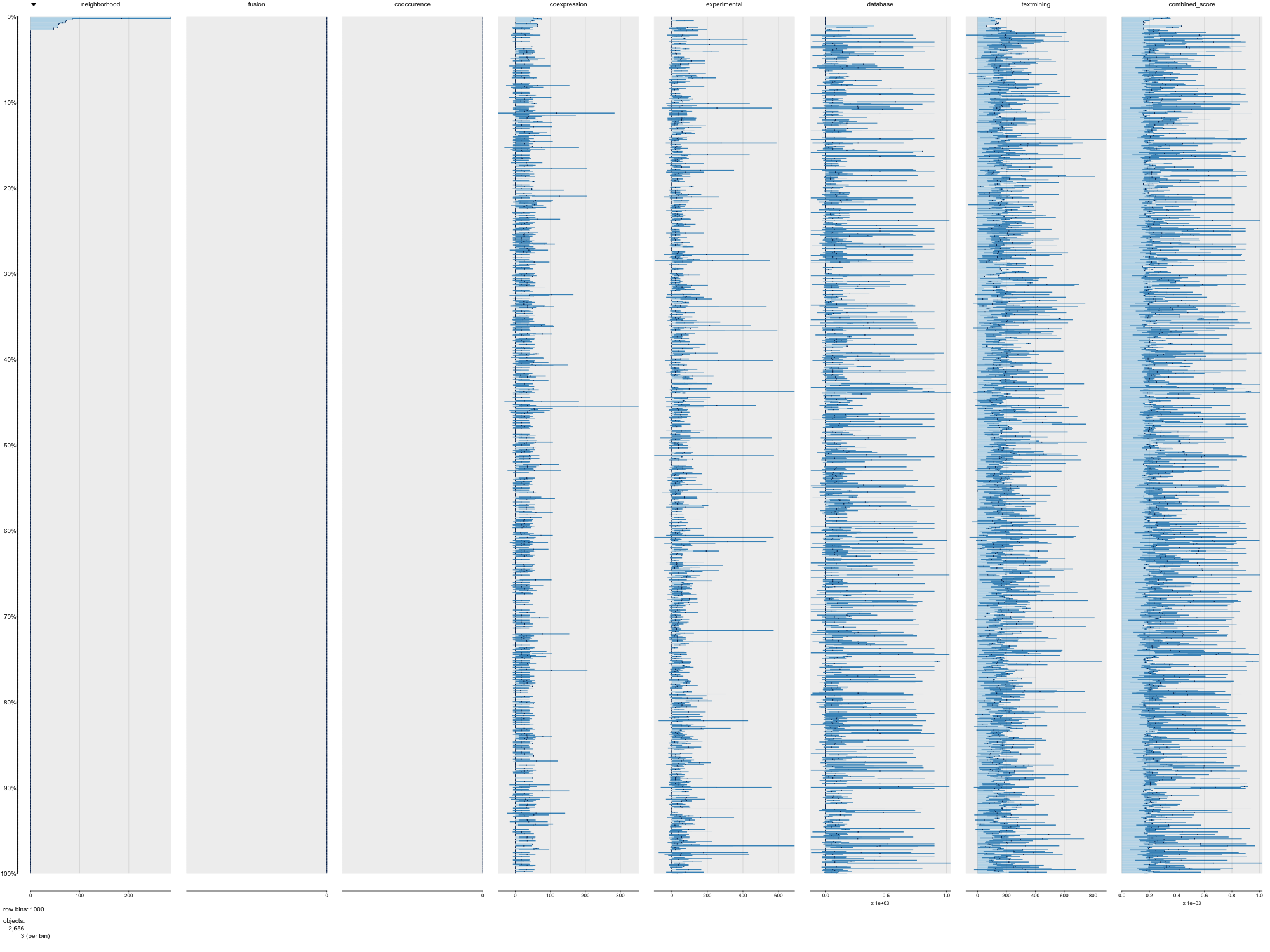

### SWISSPROT_STRING_Xtr_PAIRPLOT.png

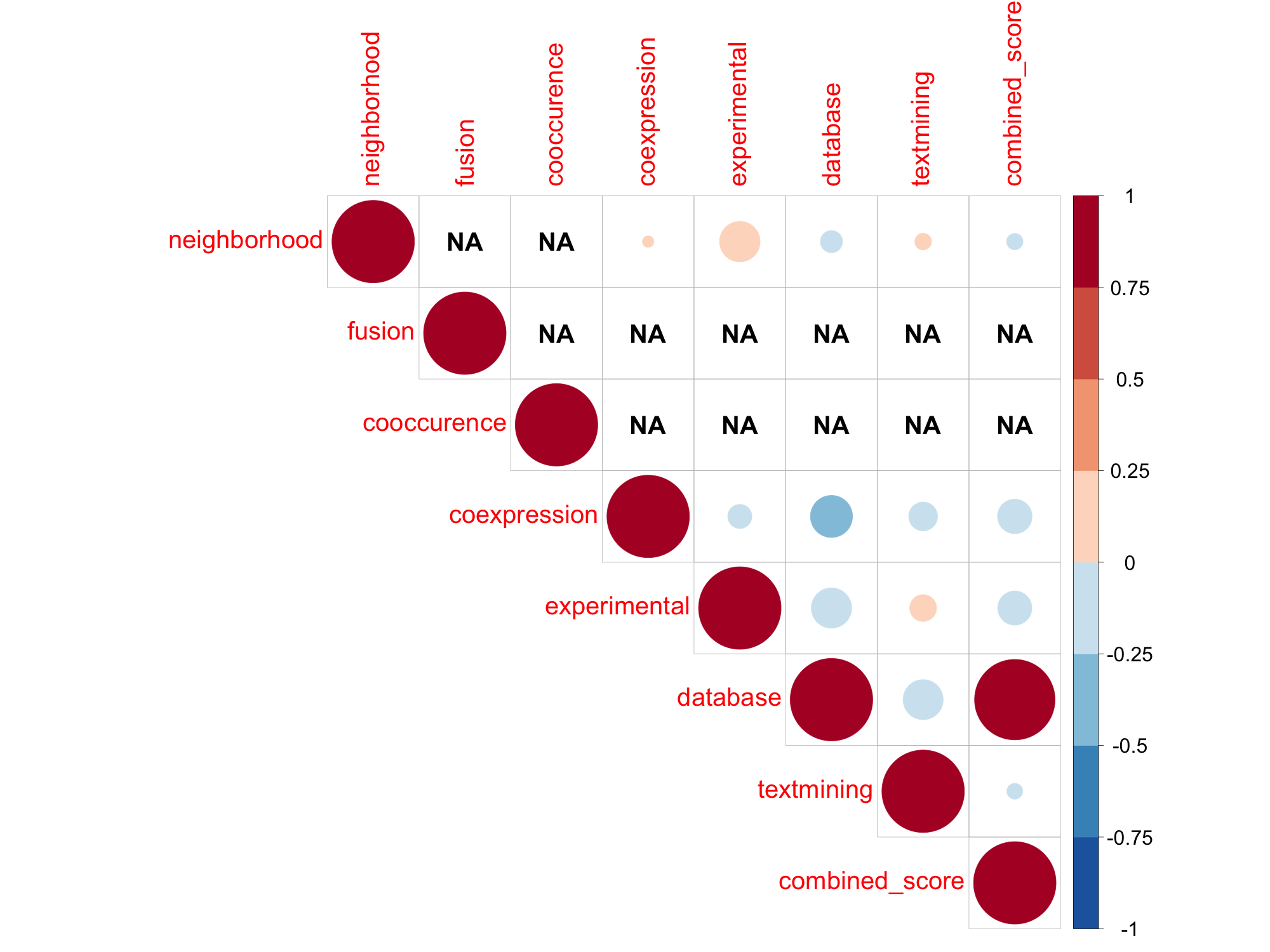

### SWISSPROT_STRING_Xtr_TABLEPLOT.png

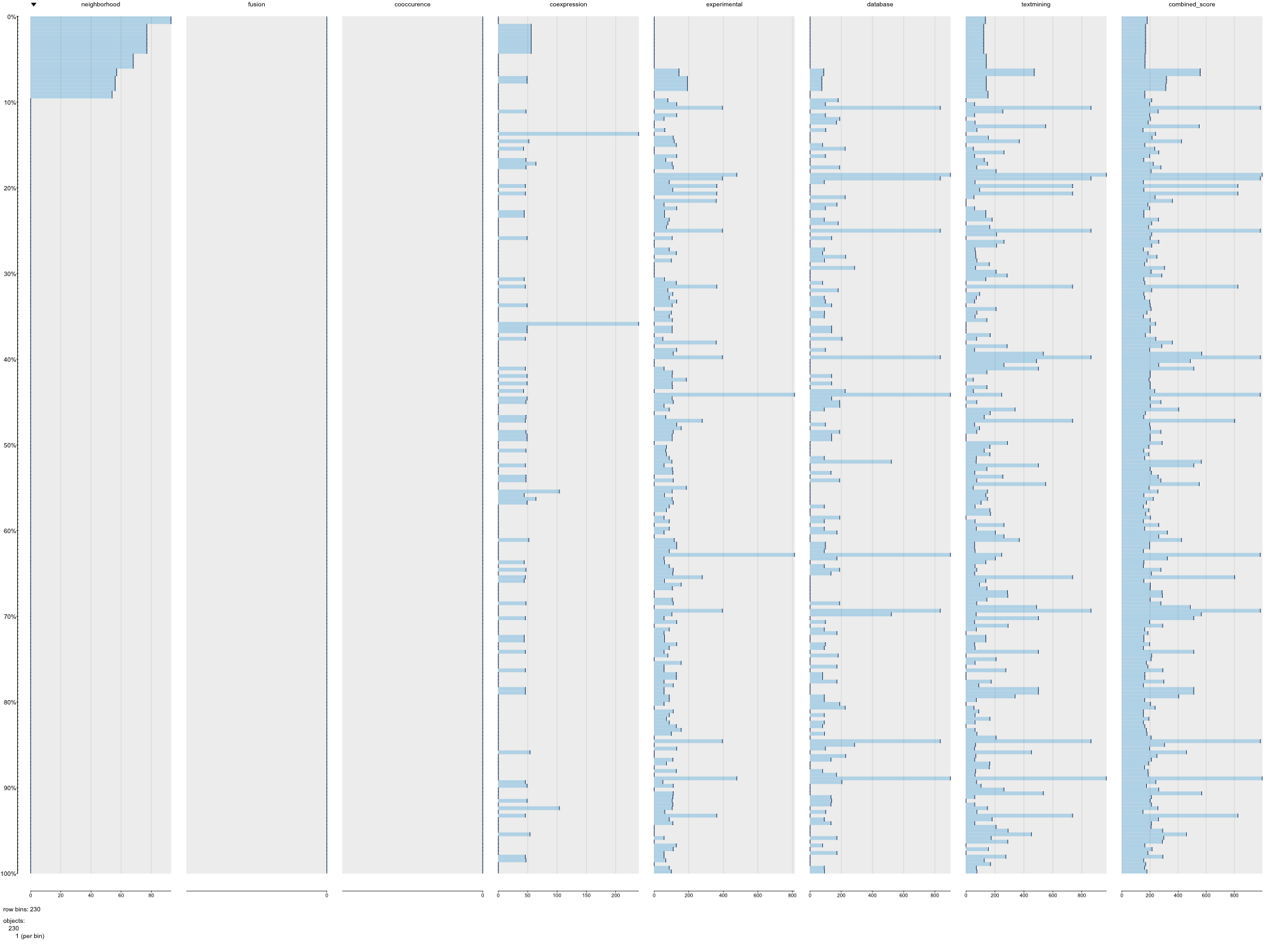

### TREMBL_STRING_Ath_PAIRPLOT.png

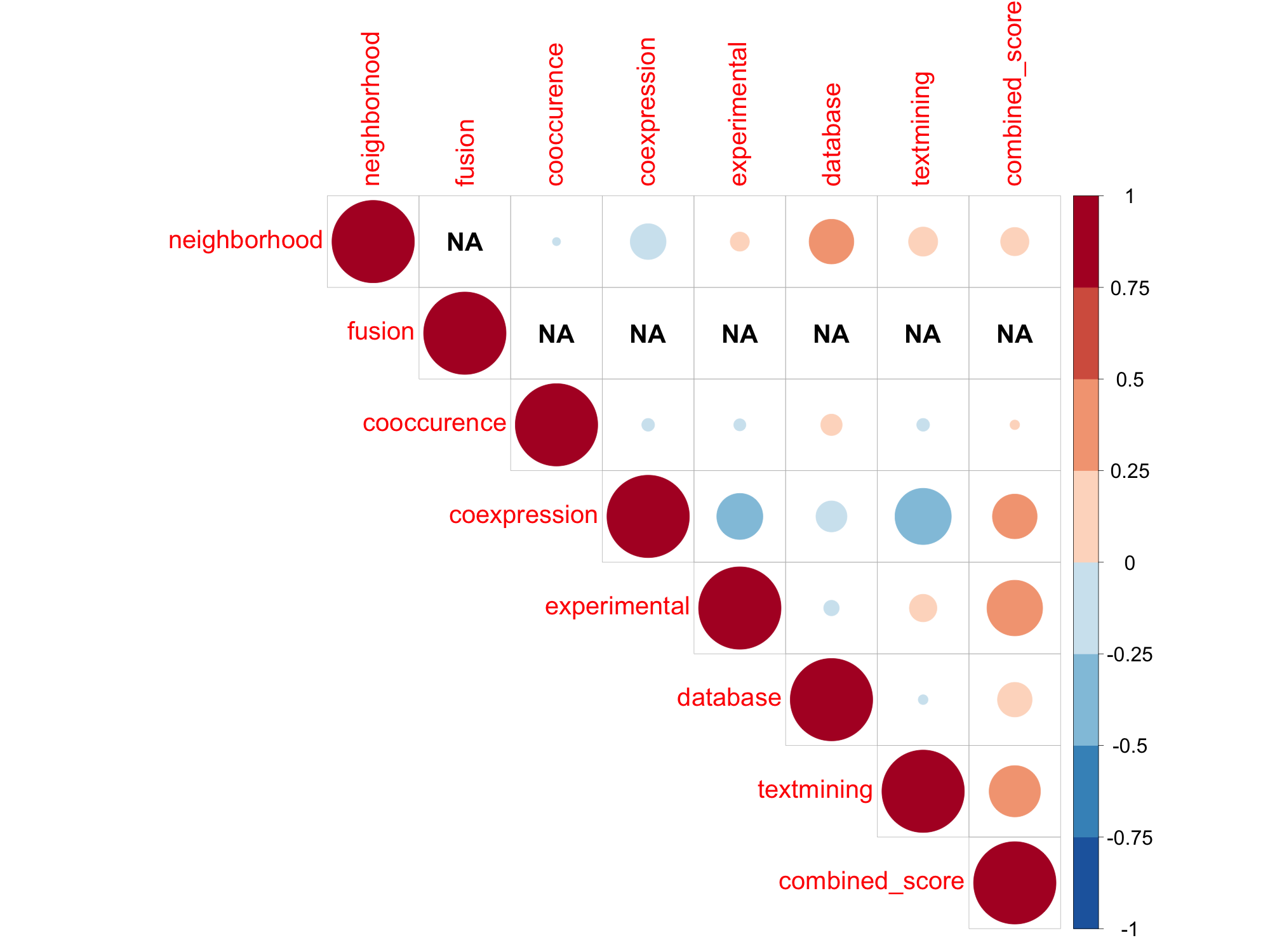

### TREMBL_STRING_Ath_TABLEPLOT.png

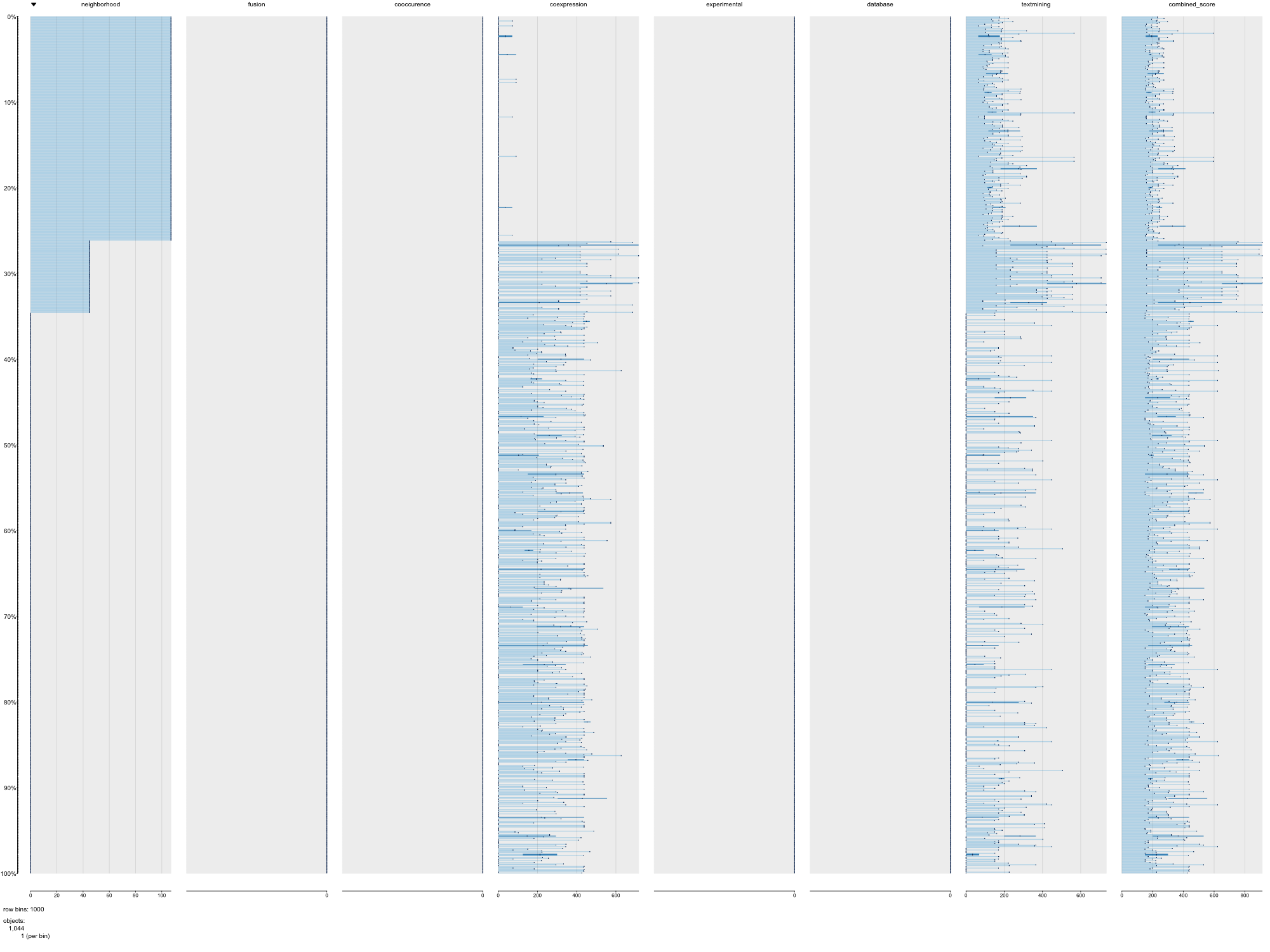

### TREMBL_STRING_Bta_PAIRPLOT.png

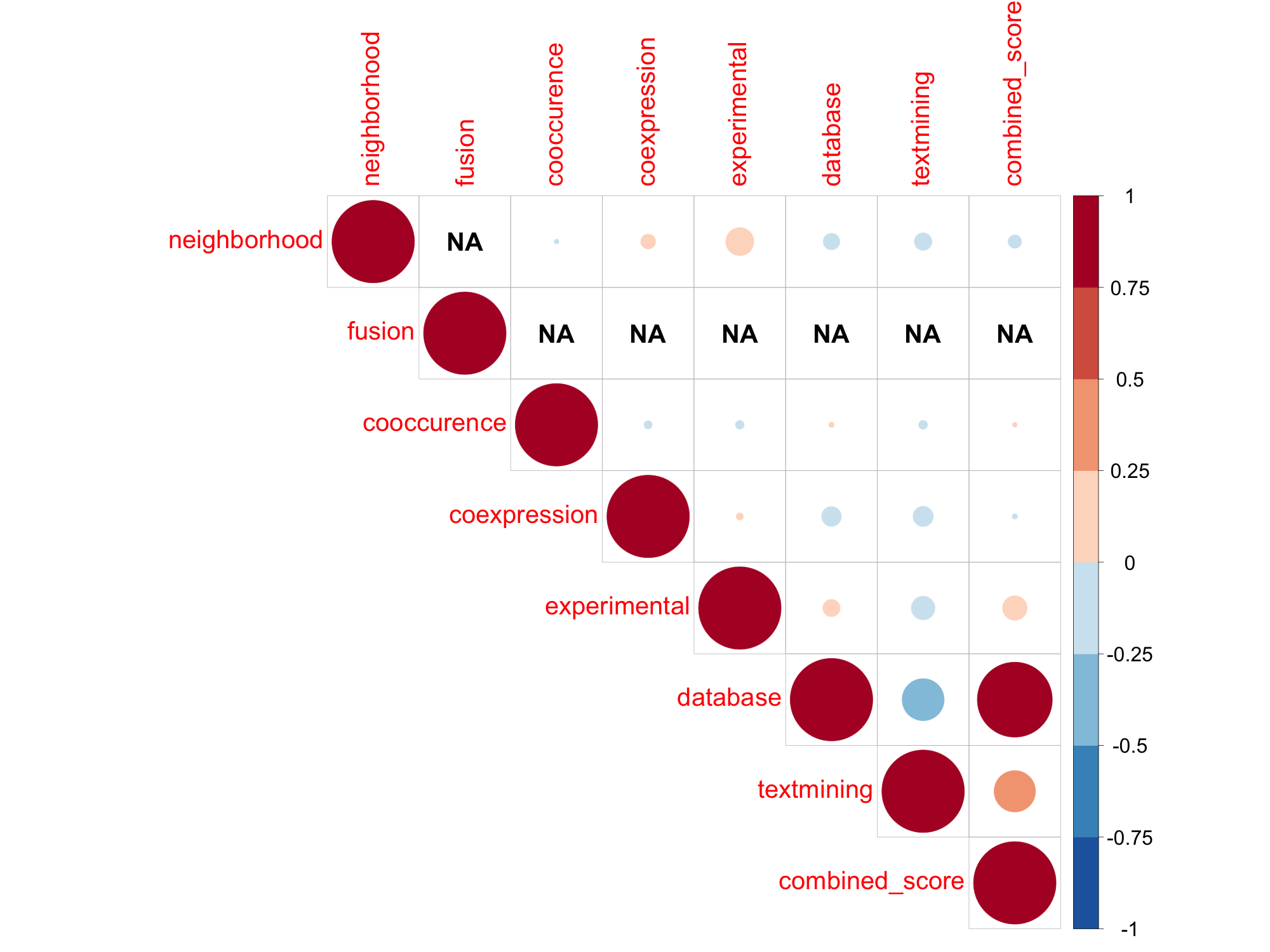

### TREMBL_STRING_Bta_TABLEPLOT.png

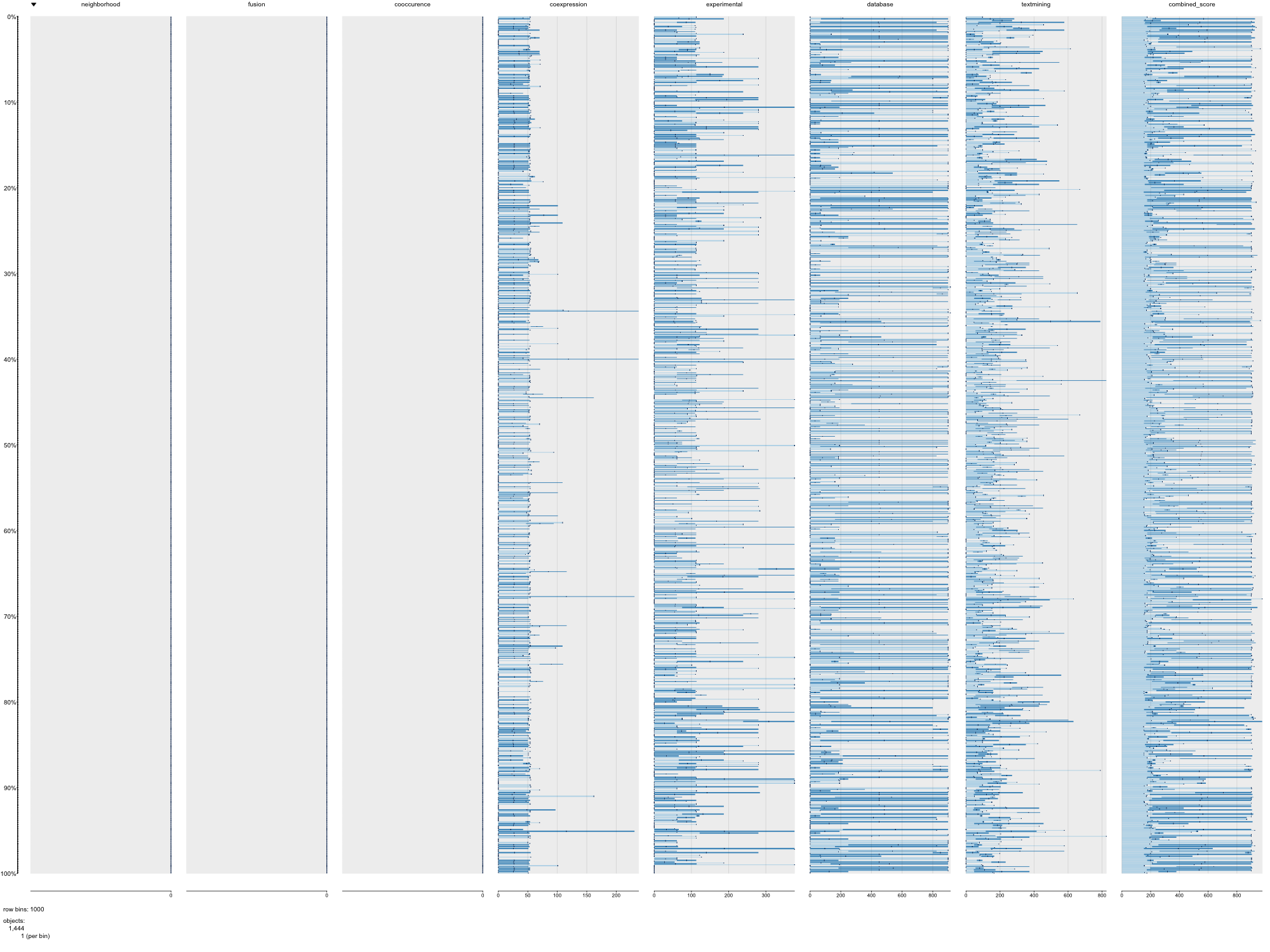

### TREMBL_STRING_Cel_PAIRPLOT.png

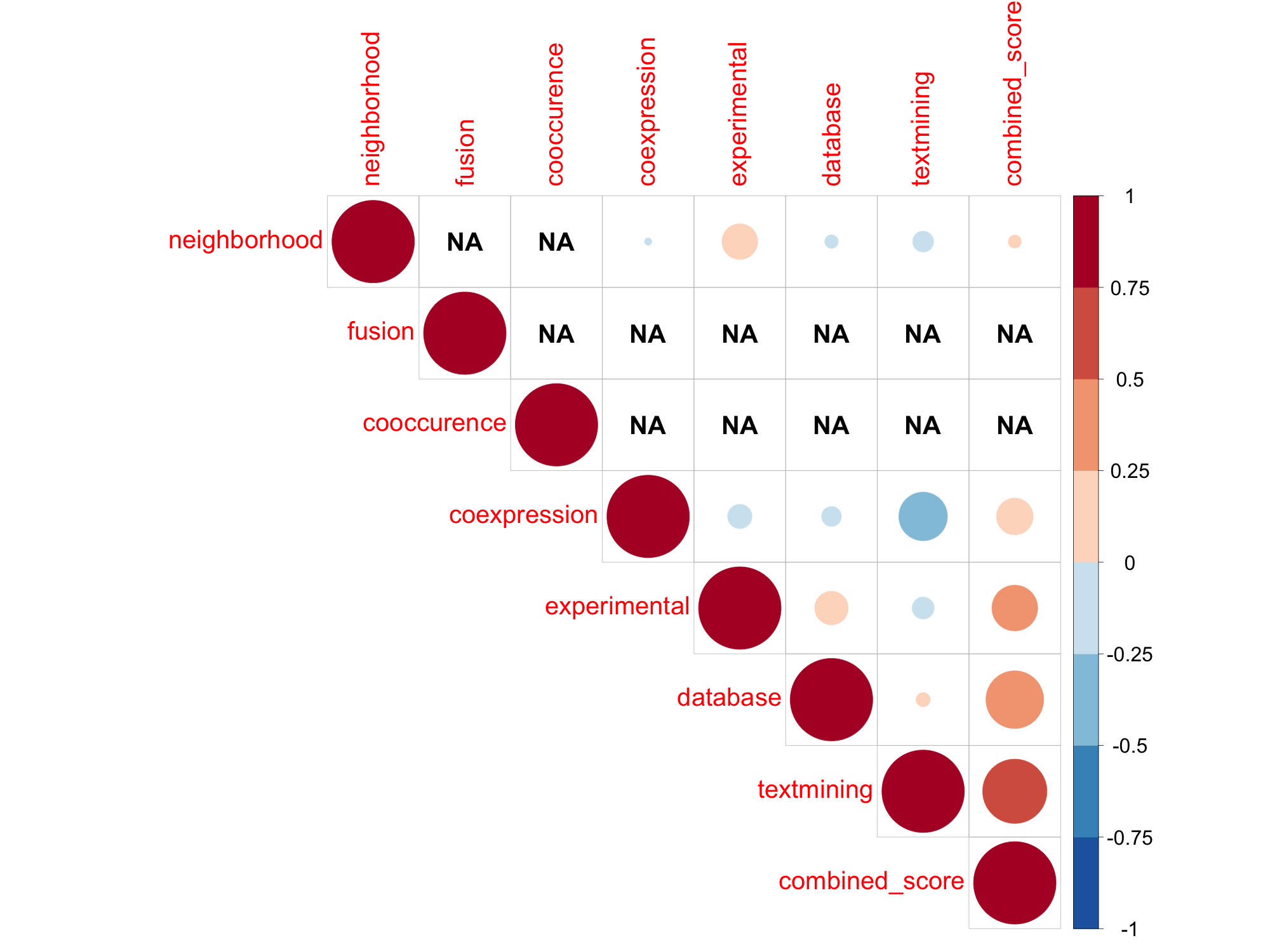

### TREMBL_STRING_Cel_TABLEPLOT.png

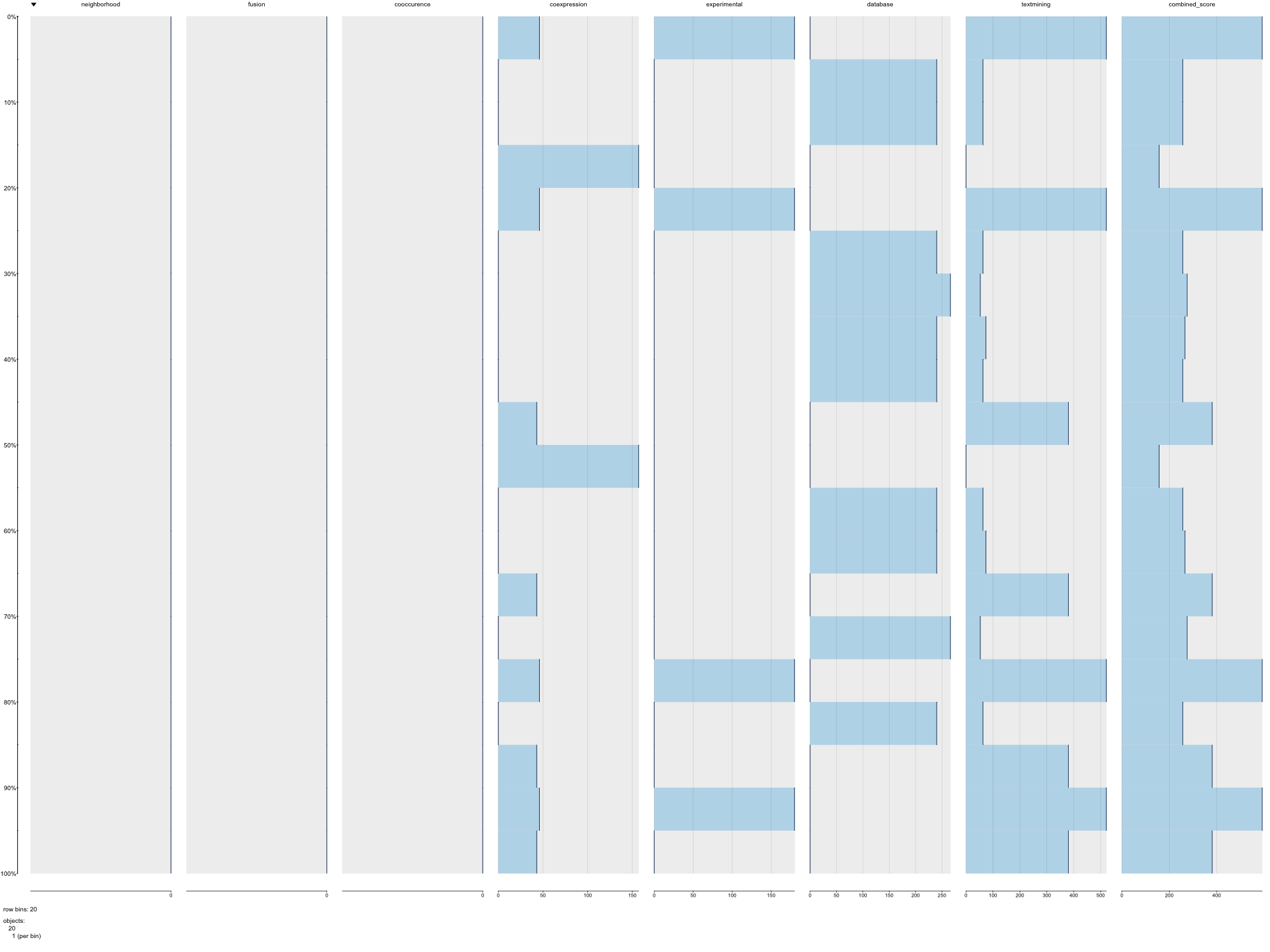

### TREMBL_STRING_Dme_PAIRPLOT.png

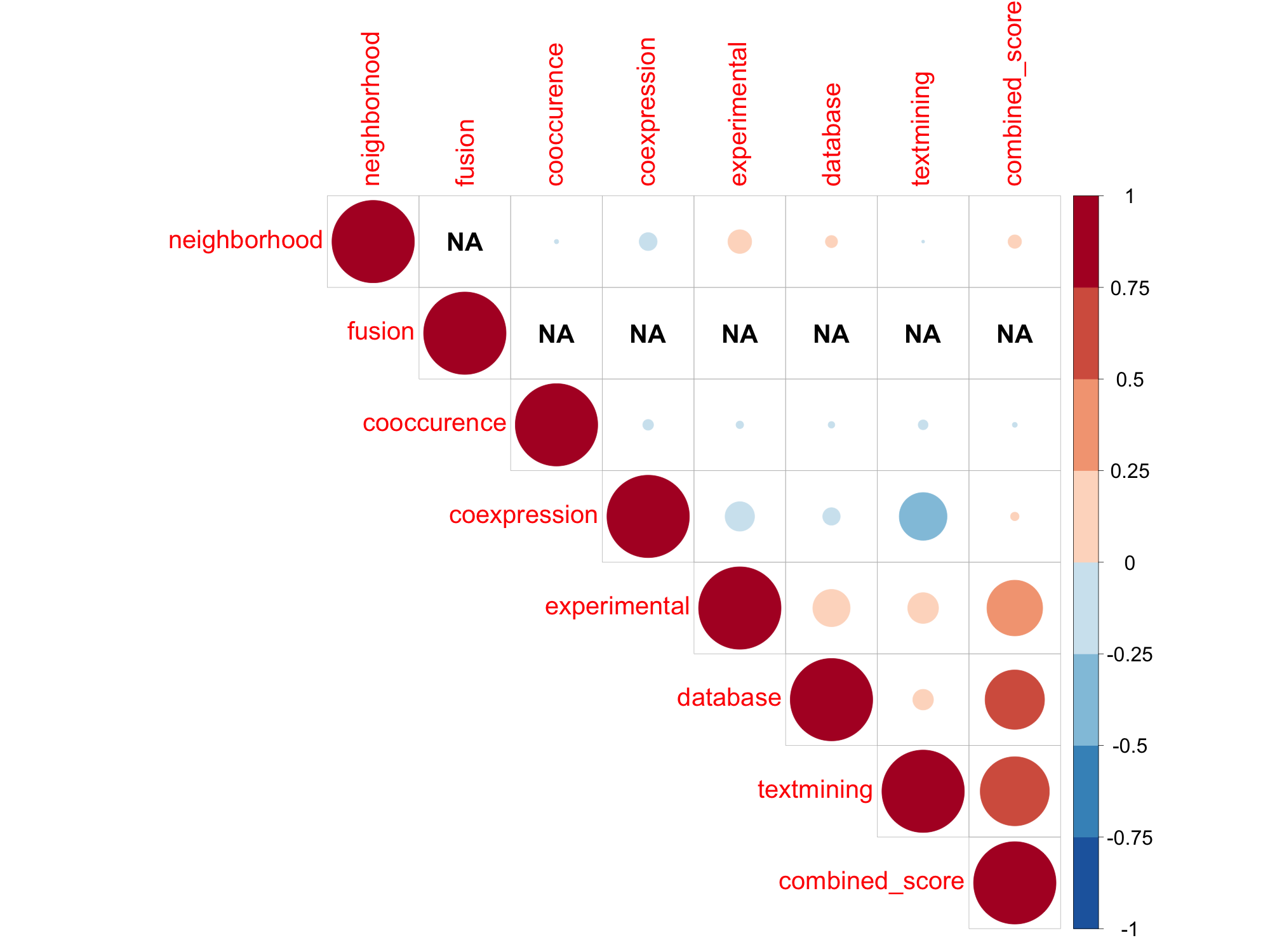

### TREMBL_STRING_Dme_TABLEPLOT.png

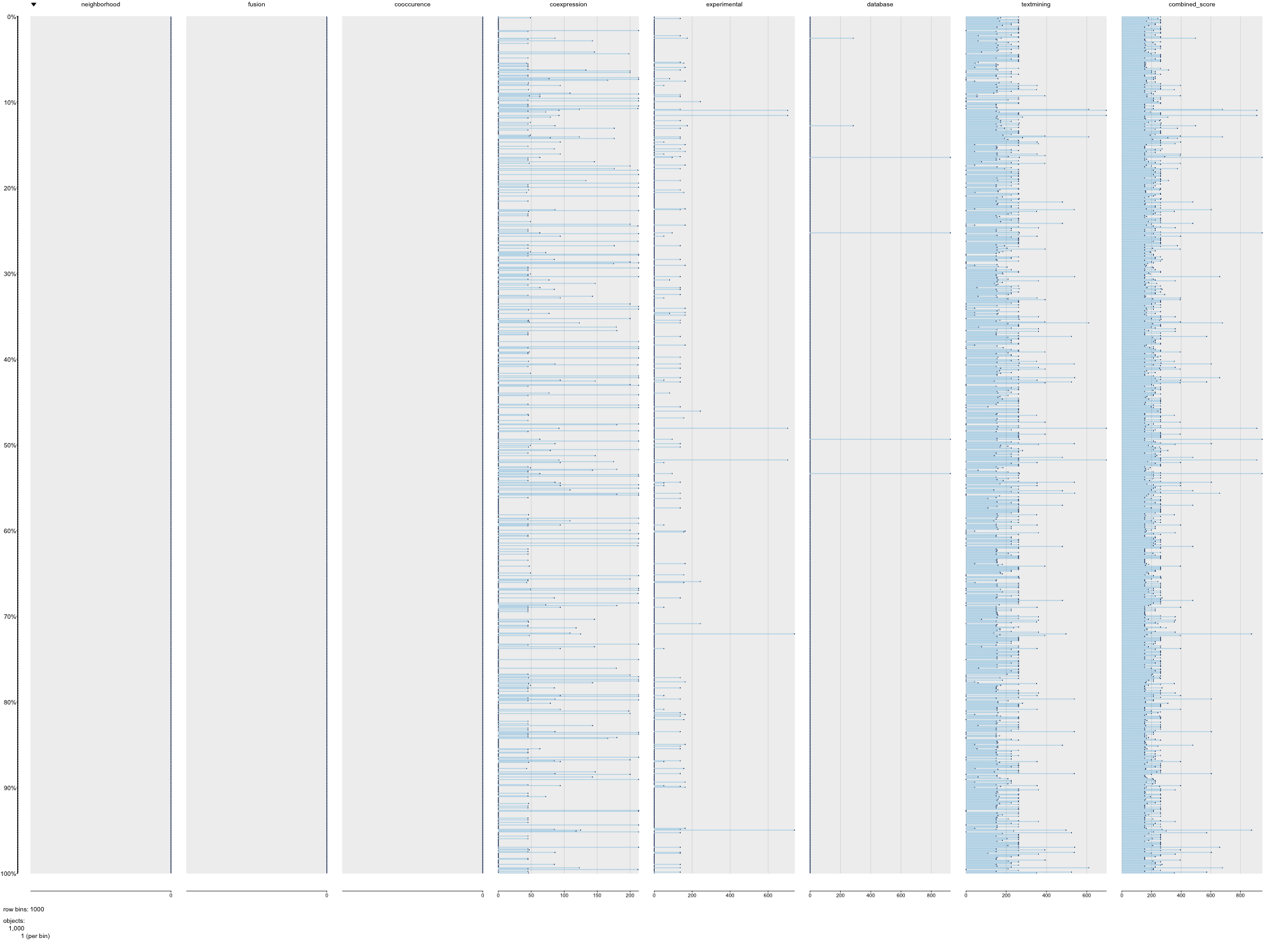

### TREMBL_STRING_Dre_PAIRPLOT.png

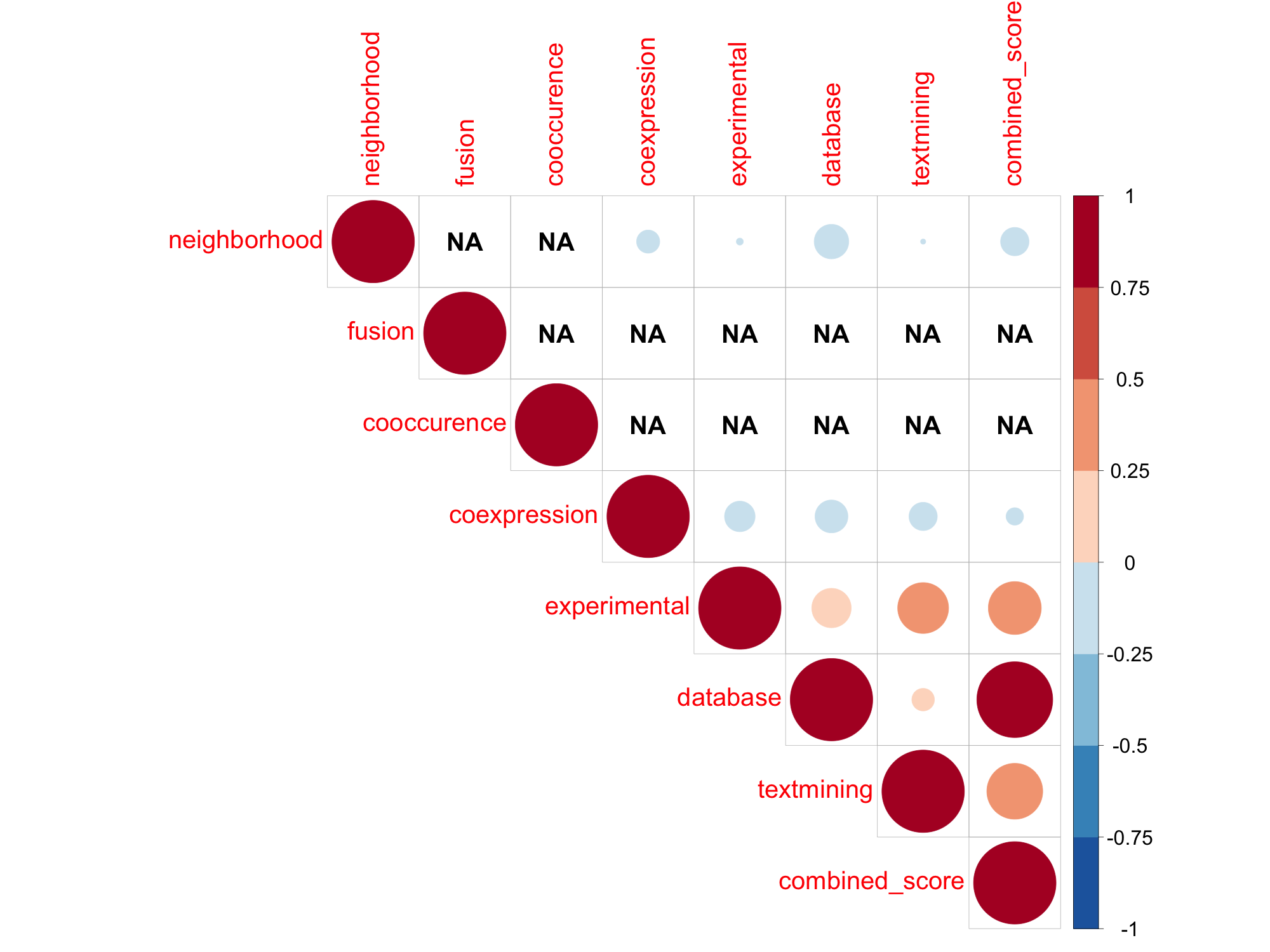

### TREMBL_STRING_Dre_TABLEPLOT.png

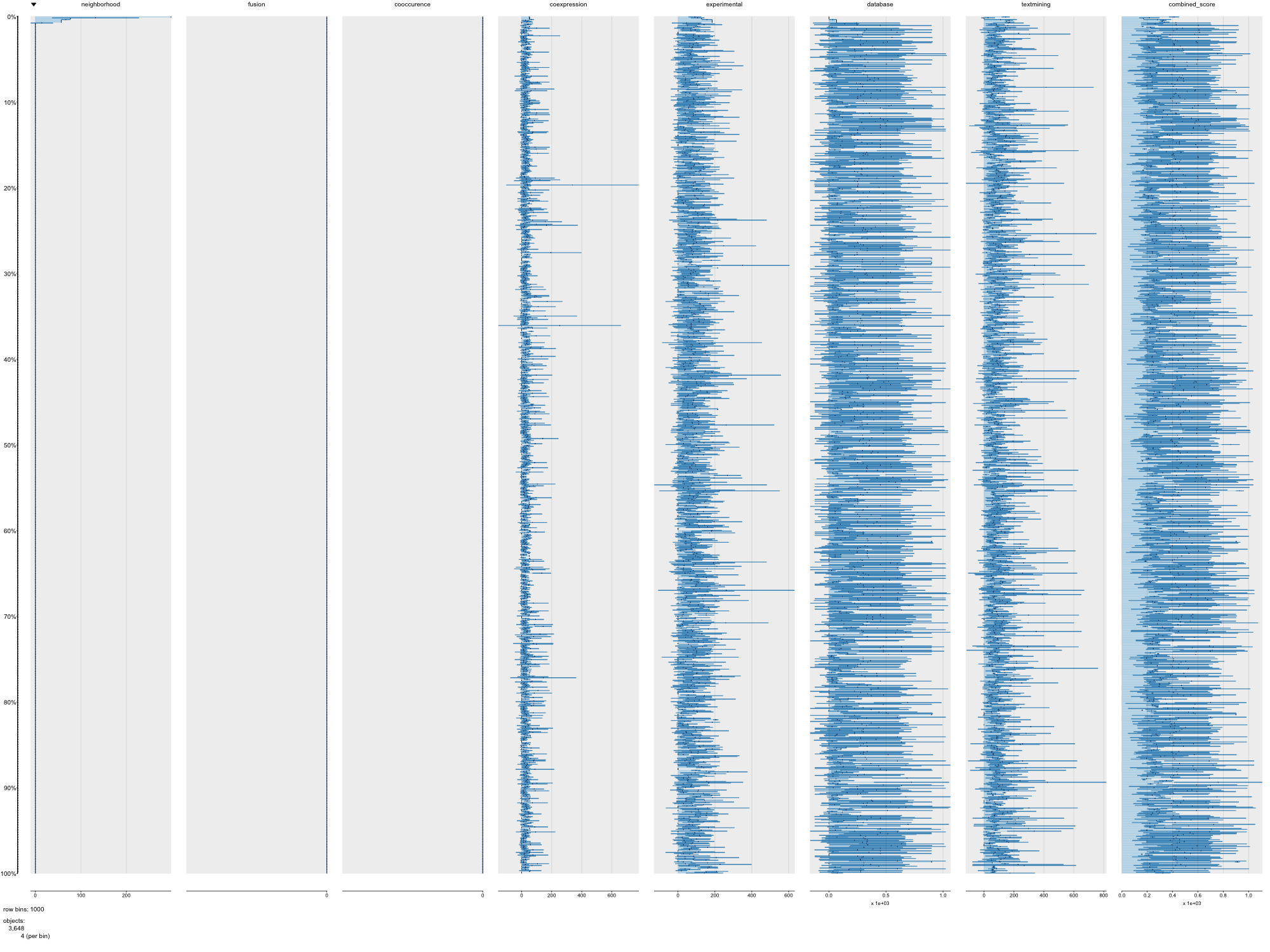

### TREMBL_STRING_Gga_PAIRPLOT.png

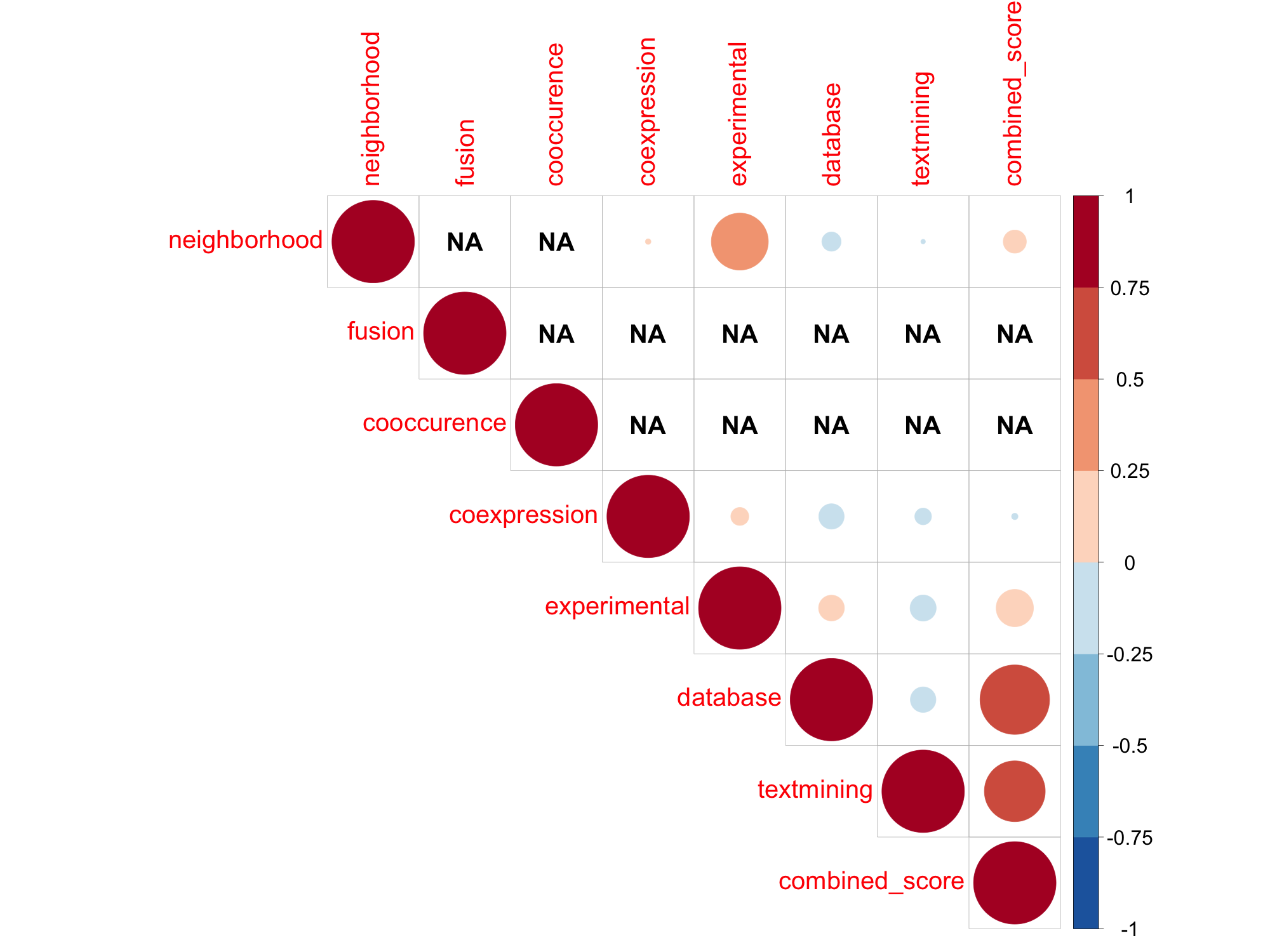

### TREMBL_STRING_Gga_TABLEPLOT.png

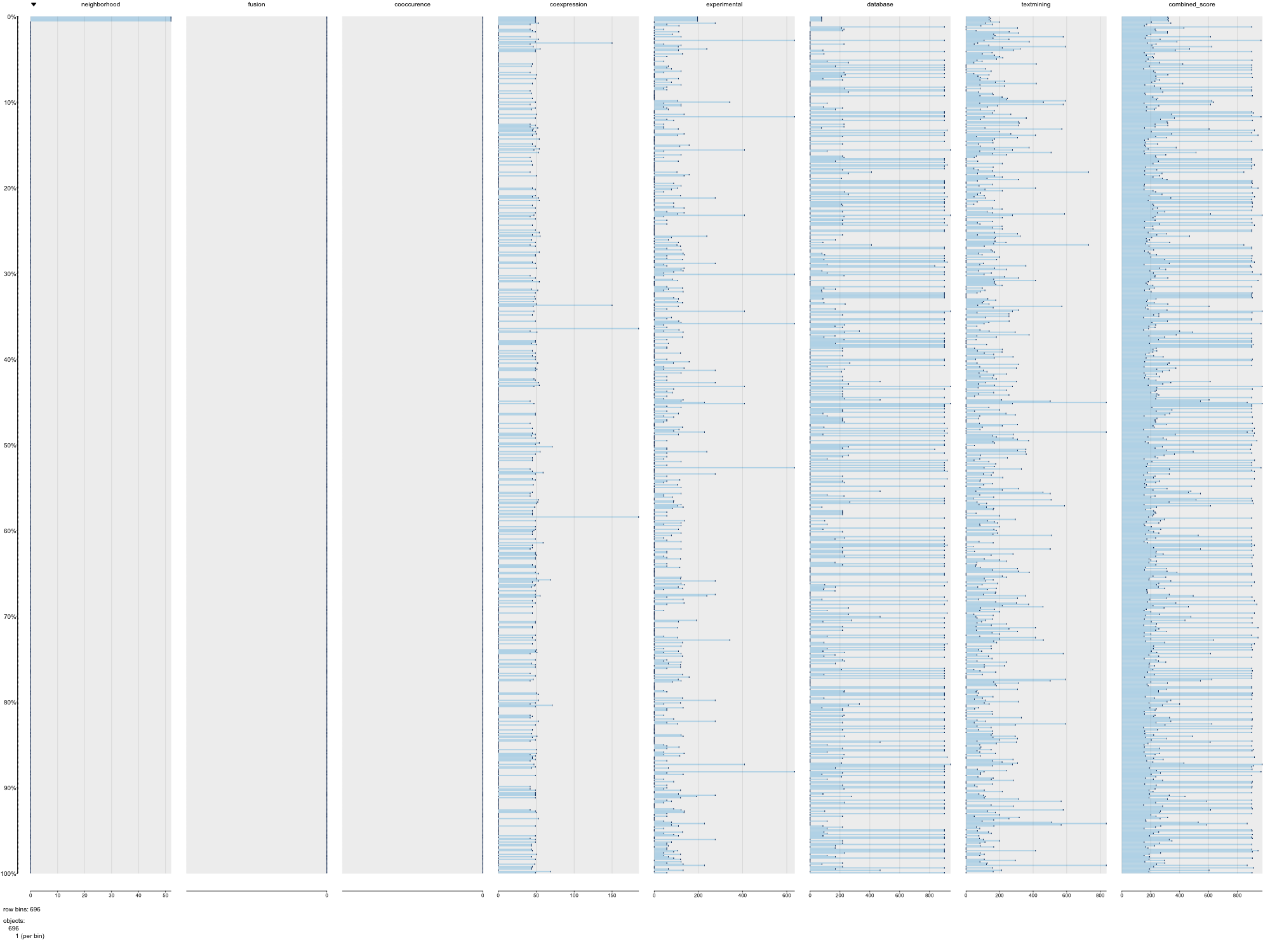

### TREMBL_STRING_Hsa_PAIRPLOT.png

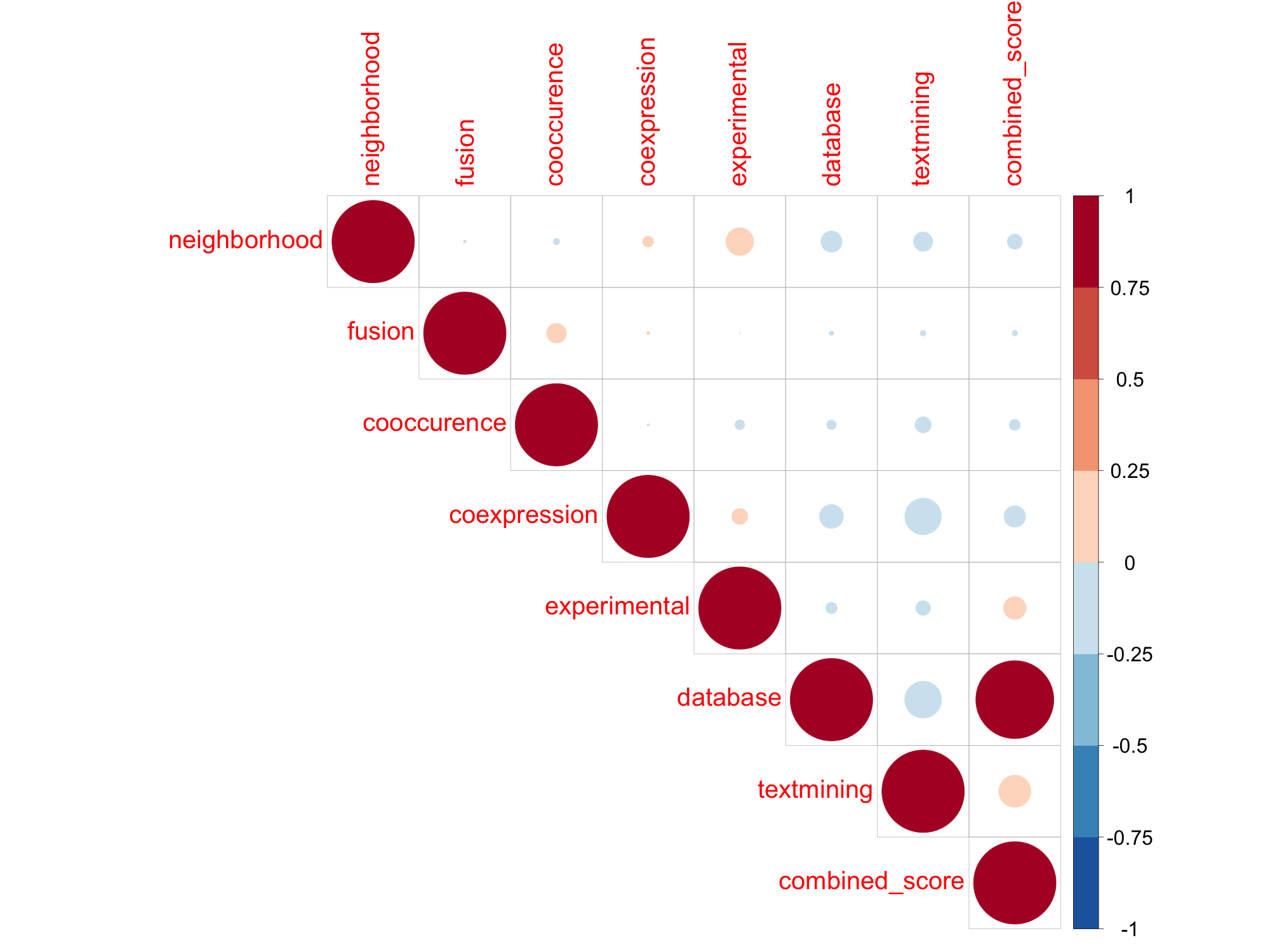

### TREMBL_STRING_Hsa_TABLEPLOT.png

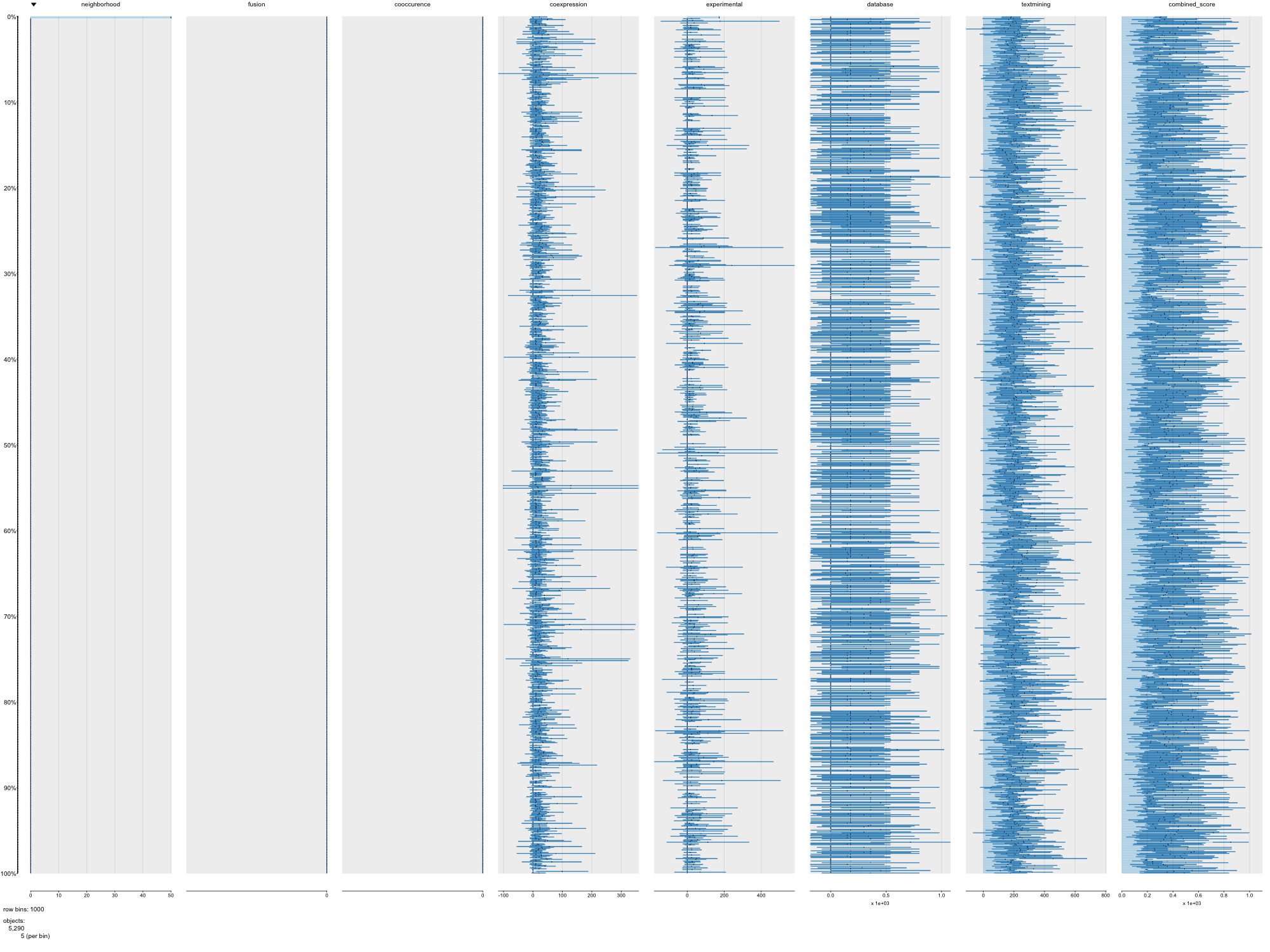

### TREMBL_STRING_Mmu_PAIRPLOT.png

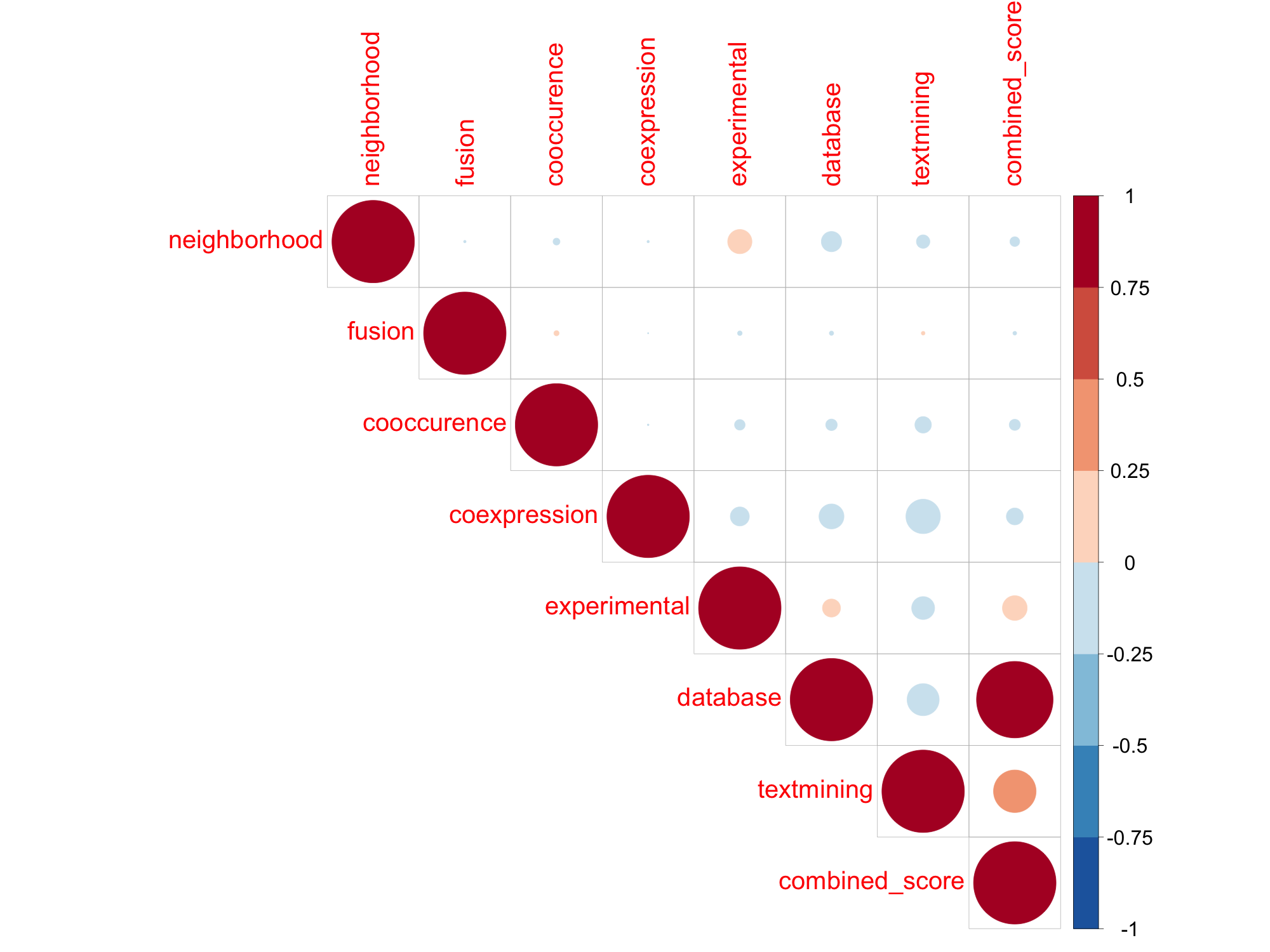

### TREMBL_STRING_Mmu_TABLEPLOT.png

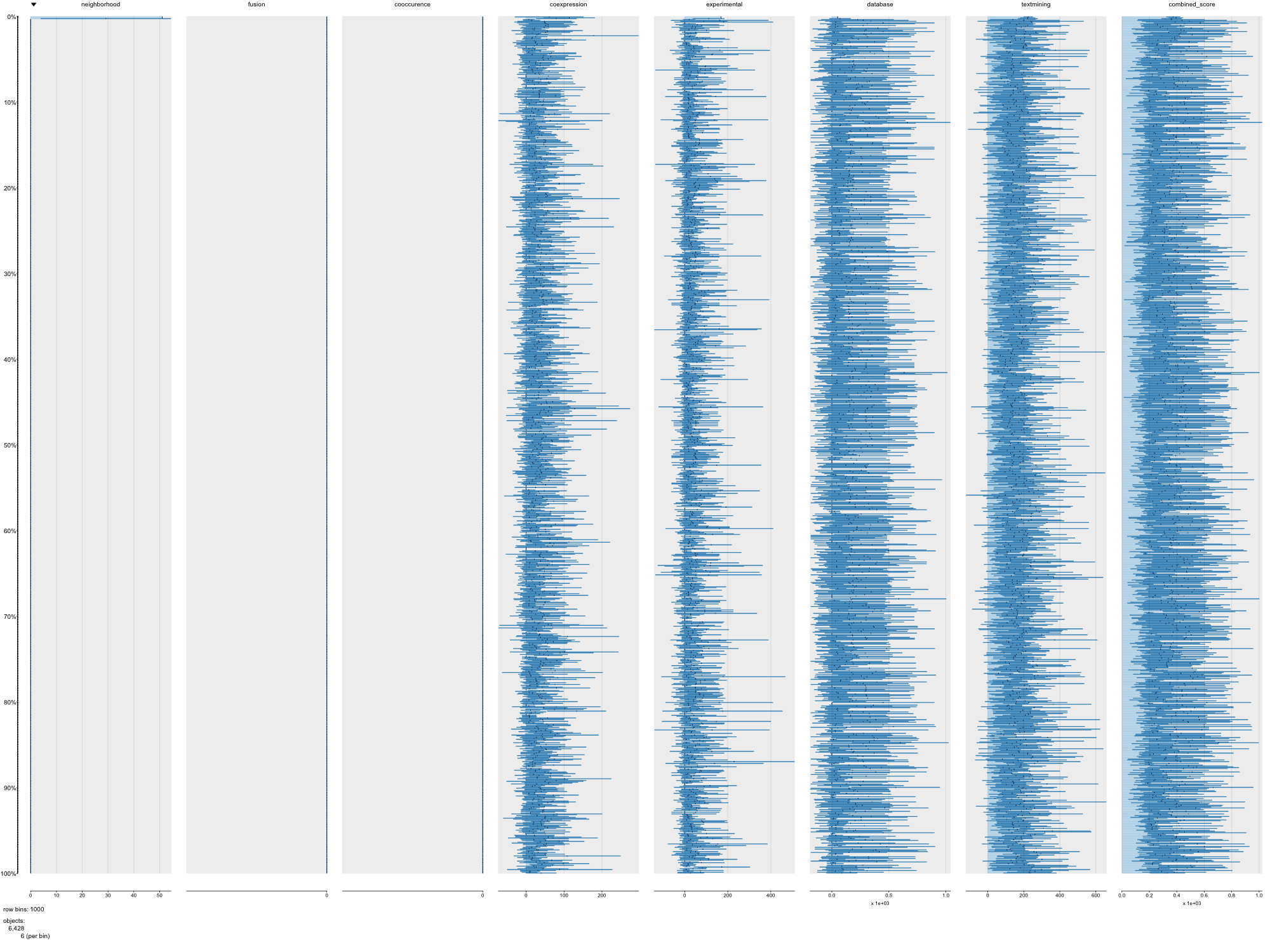

### TREMBL_STRING_Pab_PAIRPLOT.png

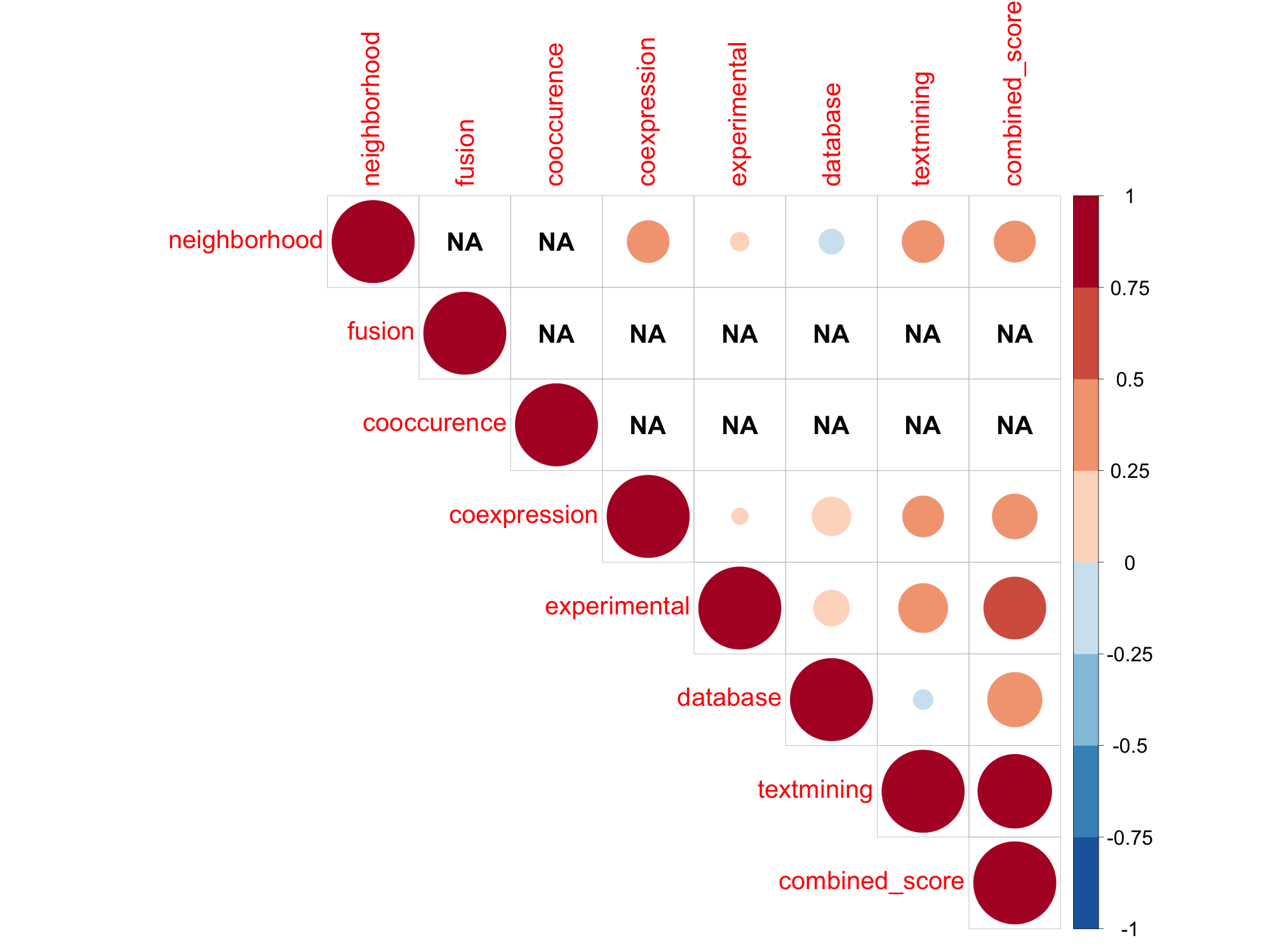

### TREMBL_STRING_Pab_TABLEPLOT.png

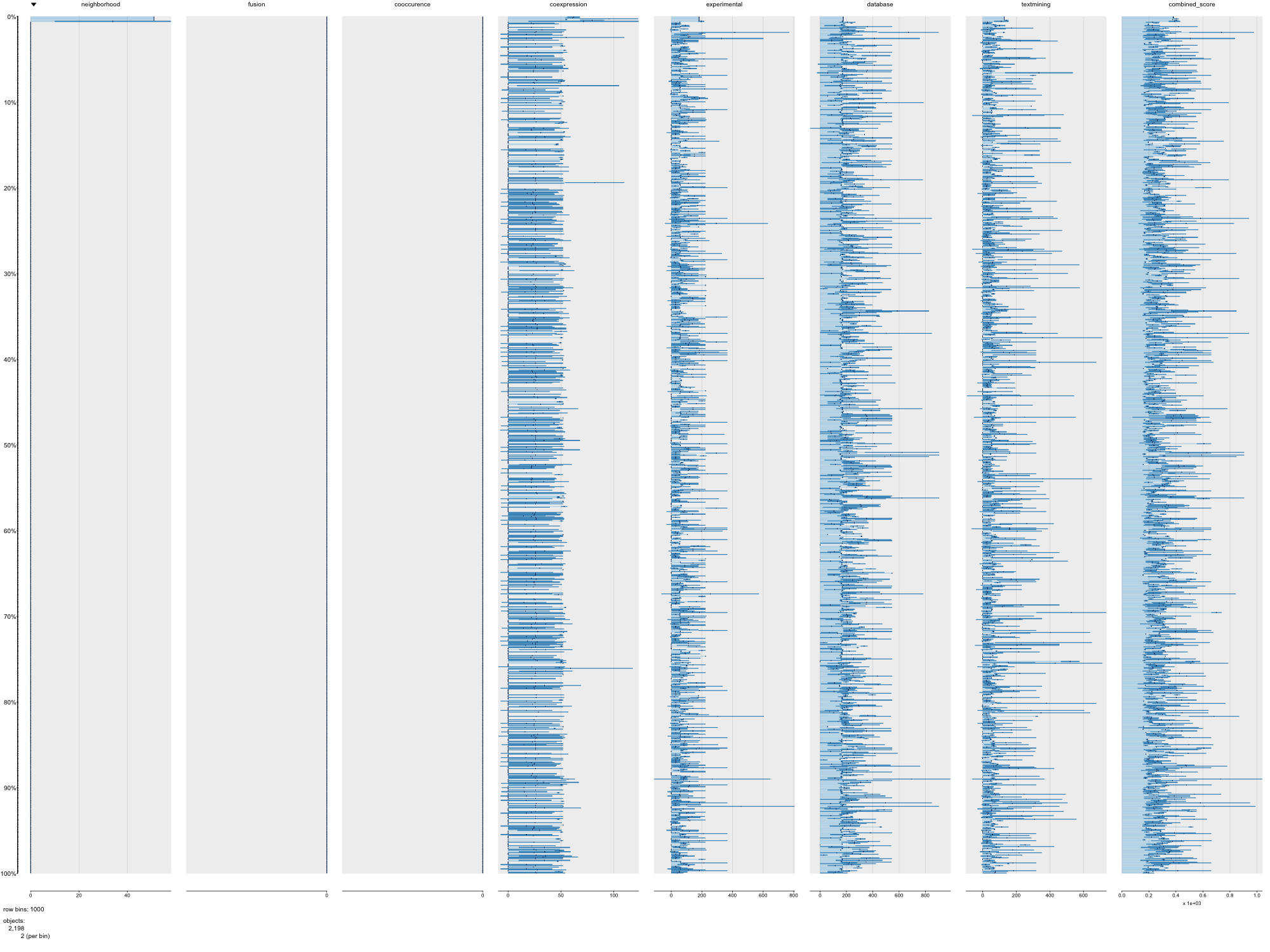

### TREMBL_STRING_Rno_PAIRPLOT.png

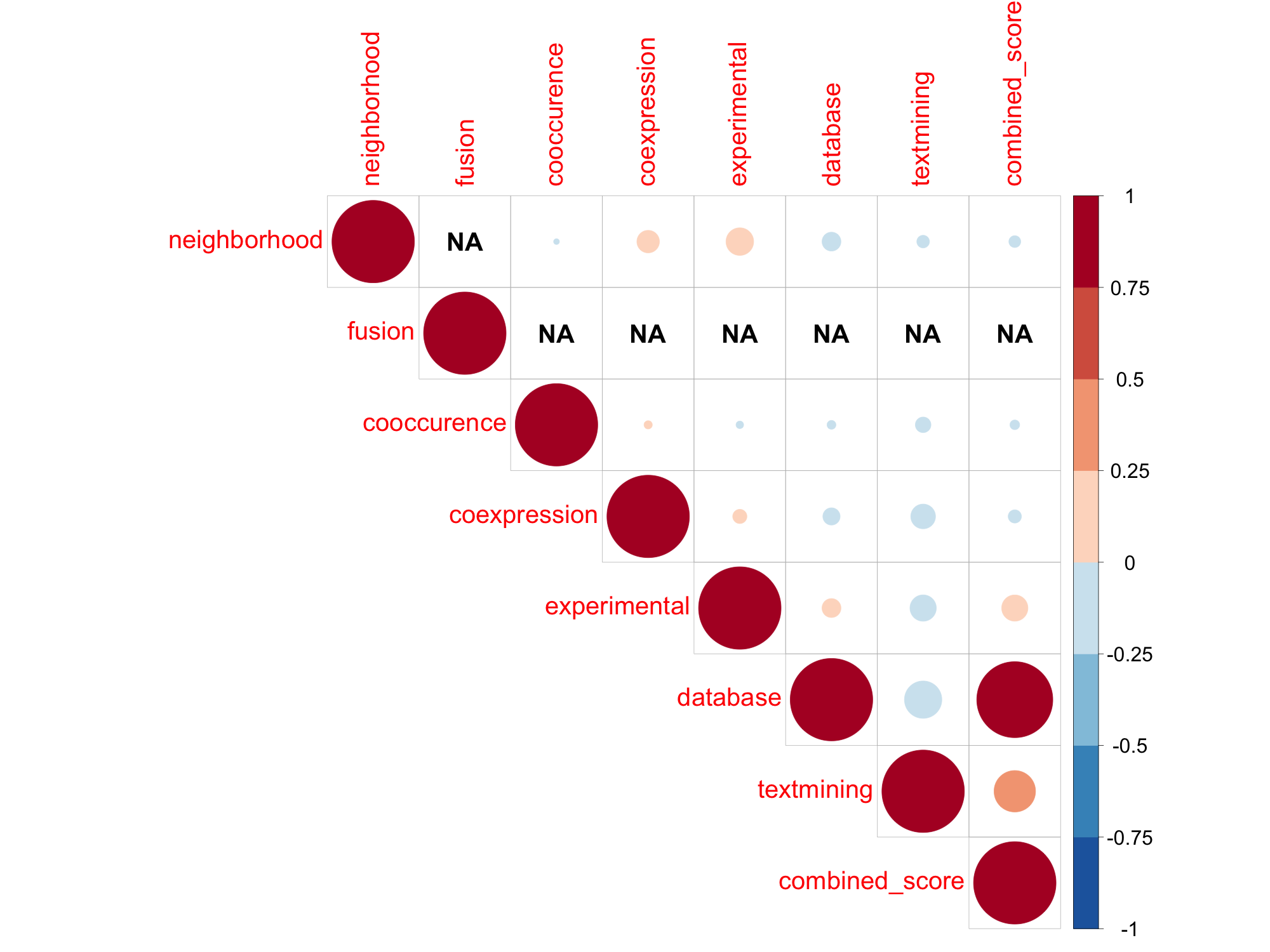

### TREMBL_STRING_Rno_TABLEPLOT.png

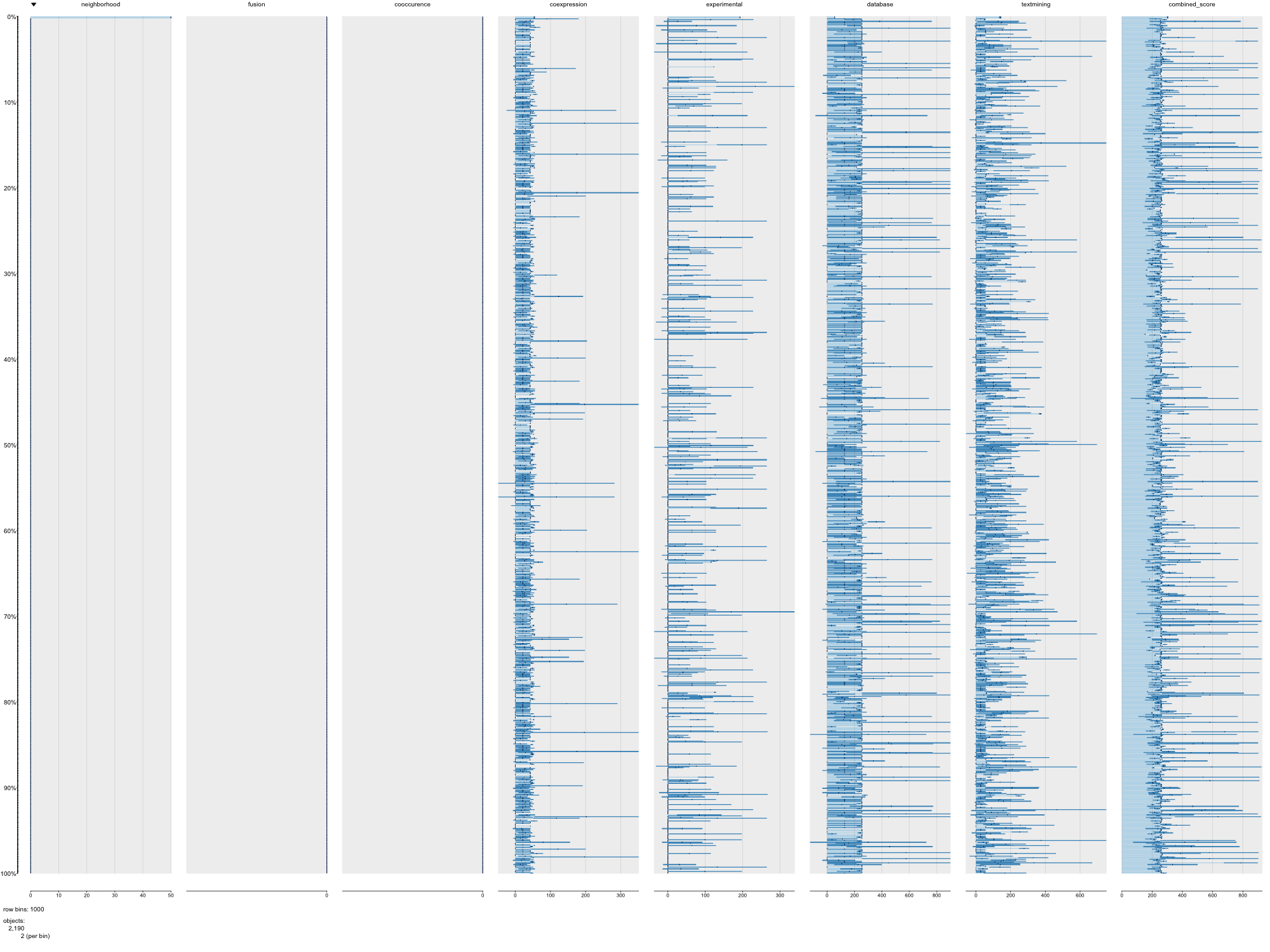

### TREMBL_STRING_Ssc_PAIRPLOT.png

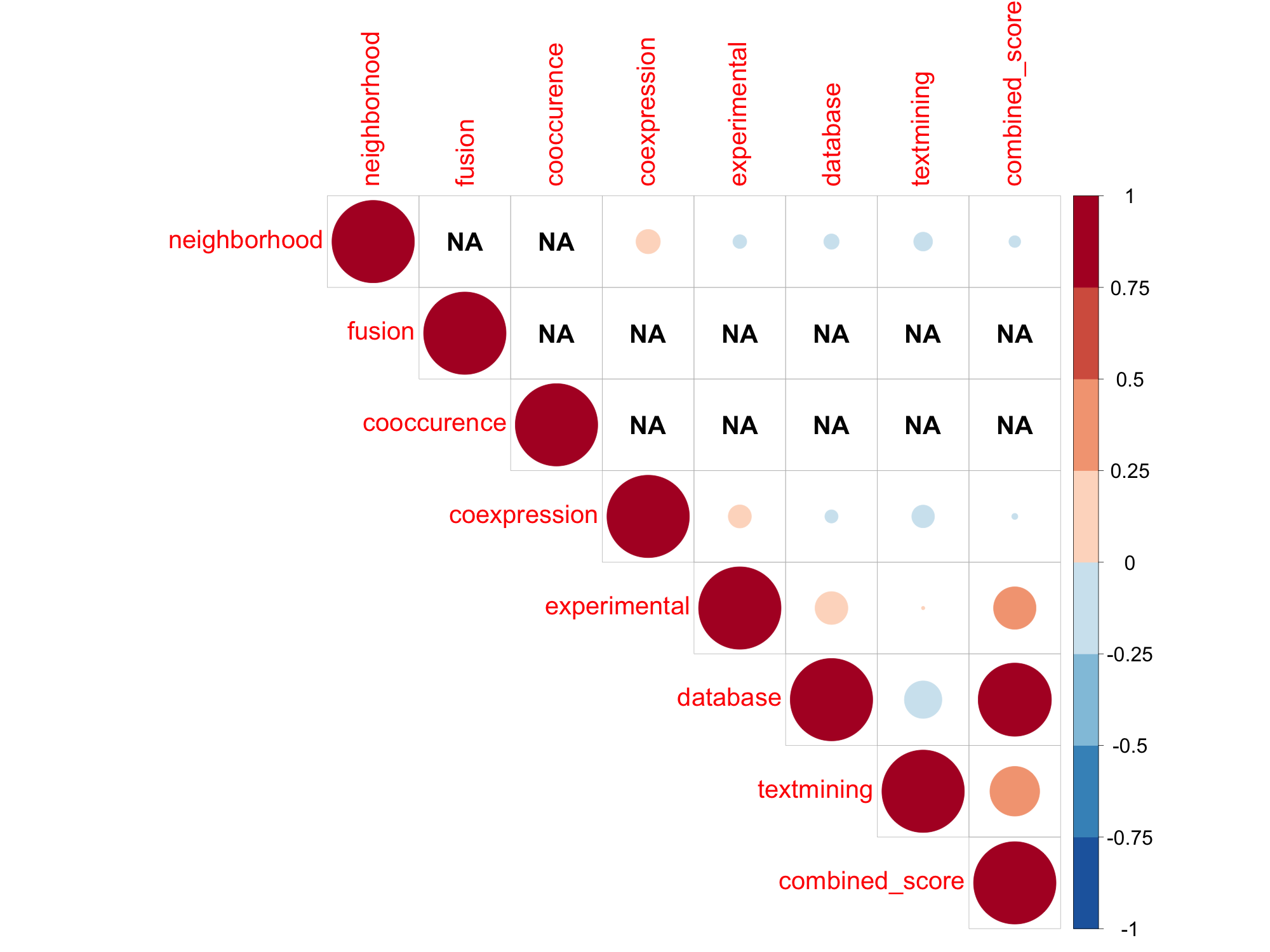

### TREMBL_STRING_Ssc_TABLEPLOT.png

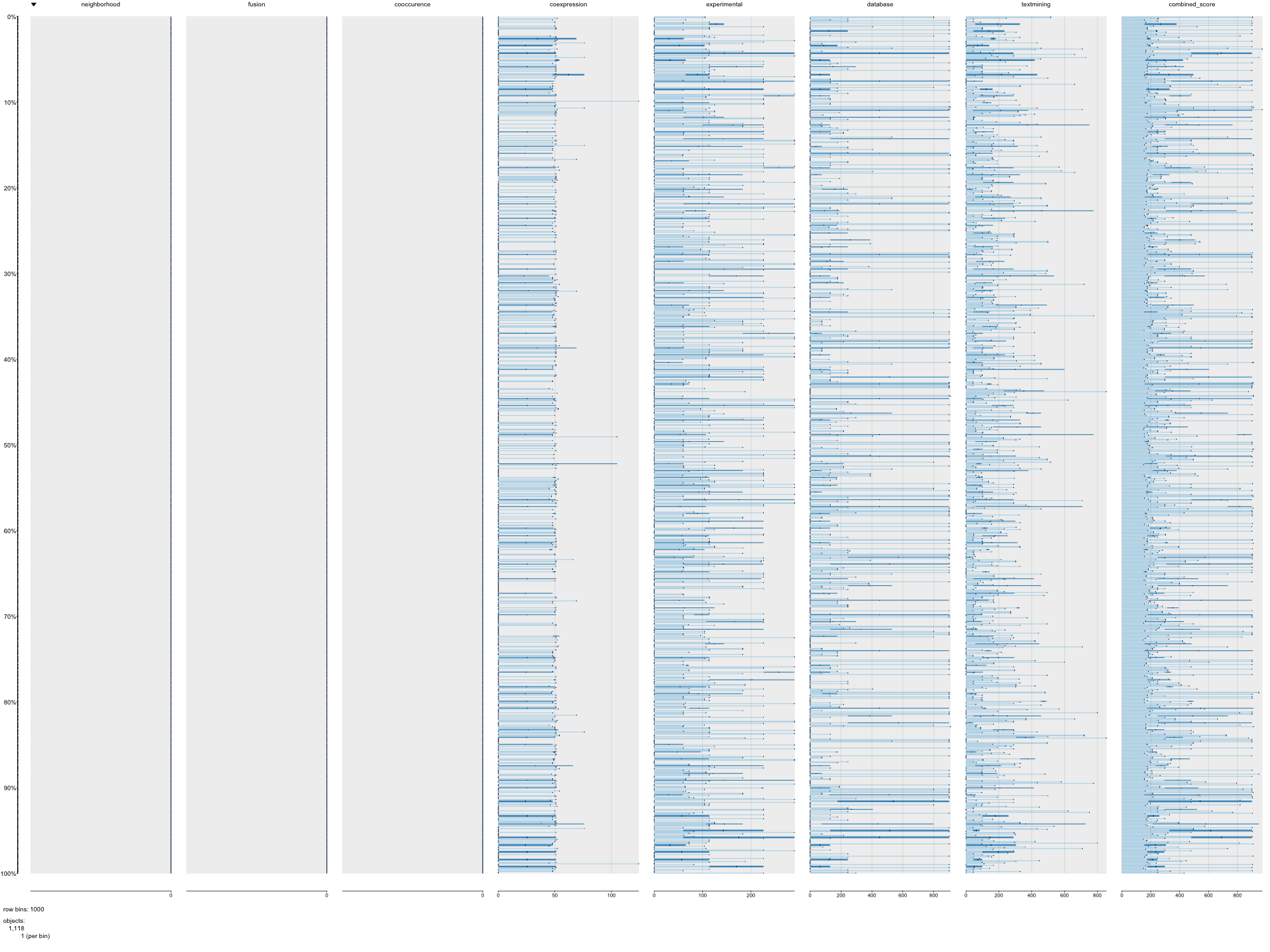

### TREMBL_STRING_Xtr_PAIRPLOT.png

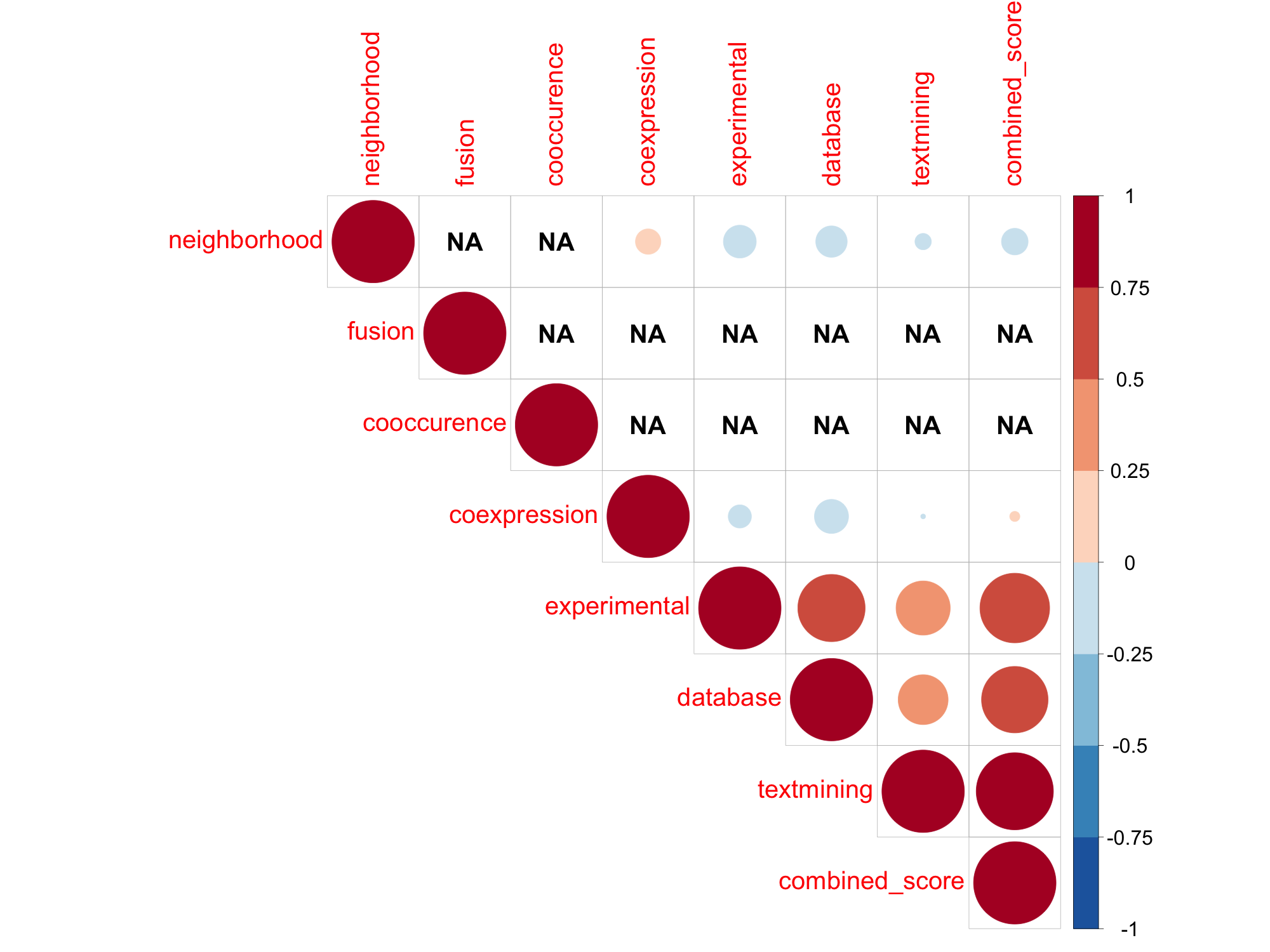

### TREMBL_STRING_Xtr_TABLEPLOT.png

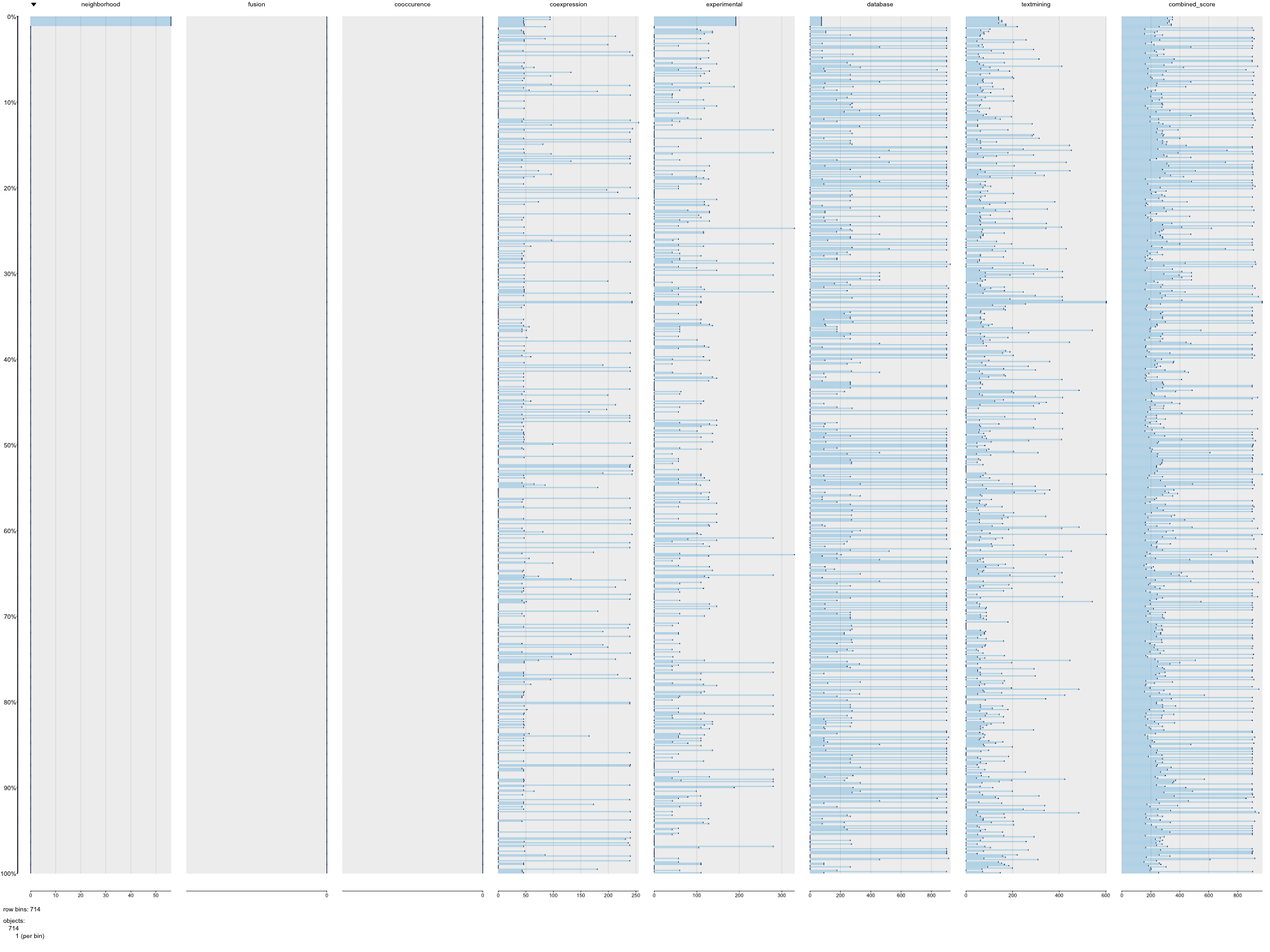
