## Additional File 3 for "Uncovering hypergraphs of cell-cell interaction from single cell RNA-sequencing data"

### Supplementary material

### SUPPLEMENTARY ANALYSES

#### Convergence of NMF, NTF, and NTD with toy model data

Here, we evaluate the convergence of non-negative matrix factorization (NMF), non-negative tensor decomposition, non-negative CP decomposition (NTF), and non-negative Tucker decomposition (NTD) with a toy model. We implemented these algorithms as a function within the nnTensor R/CRAN package (https://cran.r-project.org/web/packages/nnTensor/index.html). NTD is used in the scTensor package, and NMF is used as the initialization method for NTD. NTF is performed to compare the difference between Tucker and CP models.

All factorization methods above commonly optimize the minimization of divergence between a data matrix/tensor and a reconstructed matrix/tensor as follows:

NMF: $Divergence(X - WH)$

NTF: $Divergence(\chi - A^{(1)} \times_2 A^{(2)} \times_3 A^{(3)})$

NTD: $Divergence(\chi - G \times_1 A^{(1)} \times_2 A^{(2)} \times_3 A^{(3)})$.

A wide variety of divergence measures can be considered, such as Frobenius (Euclidean) norm, Kullback–Leibler (KL) divergence, Itakura–Saito (IS) divergence, Pearson divergence, Hellinger divergence, and Neyman chi-square divergence [56]. Some divergence measures, such as alpha divergence and beta divergence, generalize divergence as different values of $\alpha$ and $\beta$ parameters. Additionally, some different update rules of factor matrices are proposed, such as projected gradient descent (PGD), multiplicative update (MU), hierarchical alternating least squares (HALS), and greedy coordinate descent (GCD). Although, there have been some empirical experiments for comparing different algorithms, there is no comprehensive benchmark for understanding which algorithm performs better in realistic situations. Here, we have implemented different algorithms for NMF, NTF, and NTD and compared their performance with toy model data and empirical datasets. Each algorithm can be specified using the optional parameter "algorithm" for the NMF, NTF, and NTD functions of nnTensor (Table S1).

**Supplementary Table 1 | Algorithms implemented in the nnTensor package.** NMF, NTF, and NTD functions have algorithm options and a wide variety of algorithms can be selected.

| Algorithm | R functions and algorithm option |
|---|---|
| **NMF with Frobenius norm** | NMF(…, algorithm="**Frobenius**", …) |
| **NMF with KL divergence** | NMF(…, algorithm="**KL**", …) |
| **NMF with IS divergence** | NMF(…, algorithm="**IS**", …) |
| **NMF with Pearson divergence** | NMF(…, algorithm="**Pearson**", …) |
| **NMF with Hellinger divergence** | NMF(…, algorithm="**Hellinger**", …) |
| **NMF with Neyman chi-square divergence** | NMF(…, algorithm="**Neyman**", …) |
| **NMF with HALS** | NMF(…, algorithm="**HALS**", …) |
| **NMF with PGD** | NMF(…, algorithm="**PGD**", …) |
| **NMF with GCD** | NMF(…, algorithm="**GCD**", …) |
| **NMF with alpha divergence** | NMF(…, algorithm="**Alpha**", Alpha=1, …) |
| **NMF with beta divergence** | NMF(…, algorithm="**Beta**", Beta=2, …) |
| **NTF with Frobenius norm** | NTF(…, algorithm="**Frobenius**", …) |
| **NTF with KL divergence** | NTF(…, algorithm="**KL**", …) |
| **NTF with IS divergence** | NTF(…, algorithm="**IS**", …) |
| **NTF with Pearson divergence** | NTF(…, algorithm="**Pearson**", …) |
| **NTF with Hellinger divergence** | NTF(…, algorithm="**Hellinger**", …) |
| **NTF with Neyman chi-square divergence** | NTF(…, algorithm="**Neyman**", …) |
| **NTF with HALS** | NTF(…, algorithm="**HALS**", …) |
| **NTF with α-HALS** | NTF(…, algorithm="**Alpha-HALS**", …) |
| **NTF with β-HALS** | NTF(…, algorithm="**Beta-HALS**", …) |
| **NTD with Frobenius norm** | NTD(…, algorithm="**Frobenius**", …) |
| **NTD with KL divergence** | NTD(…, algorithm="**KL**", …) |
| **NTD with IS divergence** | NTD(…, algorithm="**IS**", …) |

| | |
|---|---|
| **NTD with Pearson divergence** | NTD(…, algorithm=”**Pearson**”, …) |
| **NTD with Hellinger divergence** | NTD(…, algorithm=”**Hellinger**”, …) |
| **NTD with Neyman chi-square divergence** | NTD(…, algorithm=”**Neyman**”, …) |
| **NTD with HALS** | NTD(…, algorithm=”**HALS**”, …) |

To validate convergence by visual inspection, we prepared three kinds of data by simulating the data generation processes of NMF, NTF, and NTD algorithms (Figure S1). Most elements of each matrix and tensor were filled with non-negative Poisson-distributed values ($\lambda = 1$), and in some regions, the lambda parameter was varied ($\lambda = 100$, 150, or 200) to generate much larger values and confirm the block structures (https://github.com/rikenbit/nnTensor/blob/master/R/toyModel.R). The number of dimensions of each row, column, or depth space (rank/ranks) to explain the data structure is constrained to 5 (NMF-type), {4,4,4} (NTF-type), and {5,5,5} (NTD-type).

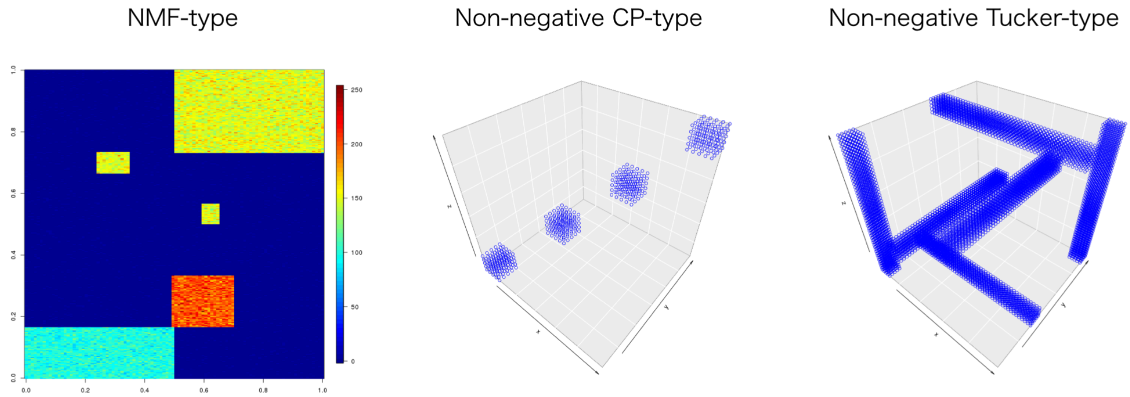

**Supplementary Figure 1 | Toy models for NMF-, NTF-, and NTD-type data**

We prepared three kinds of non-negative simulated data and evaluated the convergence of various algorithms for NMF, NTF, and NTD.

First, we evaluated the convergence of NMF algorithms with NMF-type synthetic data. We performed the NMF 30 times with rank values of 3, 5, and 10. We also performed SVD, where the solution can be uniquely determined without iteration steps, but the eigenvectors have negative values. The distribution of 30 reconstruction error values for each algorithm with rank 3 is shown in Figure S2. The error values of some algorithms, such as IS, Neyman, PGD, and GCD, showed unstable convergence, while the values of other algorithms are distributed around 3.5 and seem stable. However, the reconstructed matrix has some missing blocks. With rank 5, some algorithms, such as Frobenius and KL, recovered such blocks, and other algorithms can detect such blocks, even if the rank is set to a larger value, such as 10 (Figure S3-S4). Accordingly, we specified KL divergence as the initialization method for NTD used in the corresponding scTensor function.

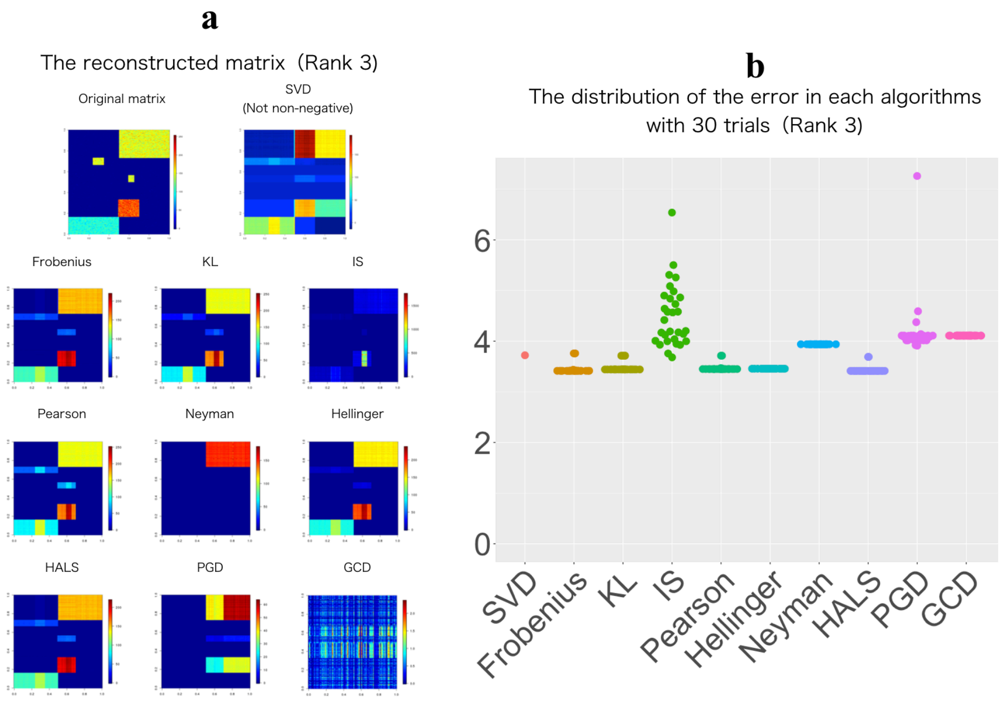

**Supplementary Figure 2 | Convergence of NMF algorithms with toy NMF-type data (Rank 3)**

(a) Reconstructed matrix from the result of each algorithm. (b) Distribution of the reconstruction error of each algorithm based on 30 trials.

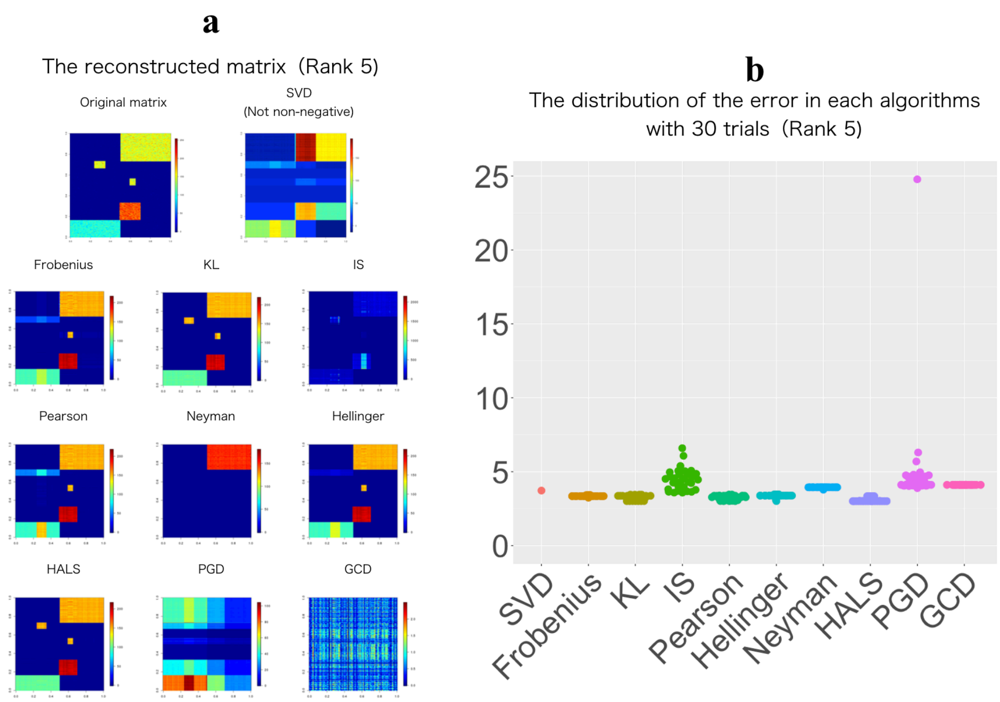

**Supplementary Figure 3 | Convergence of NMF algorithms with toy NMF-type data (Rank 5)**

(a) Reconstructed matrix from the result of each algorithm. (b) Distribution of the reconstruction error of each algorithm based on 30 trials.

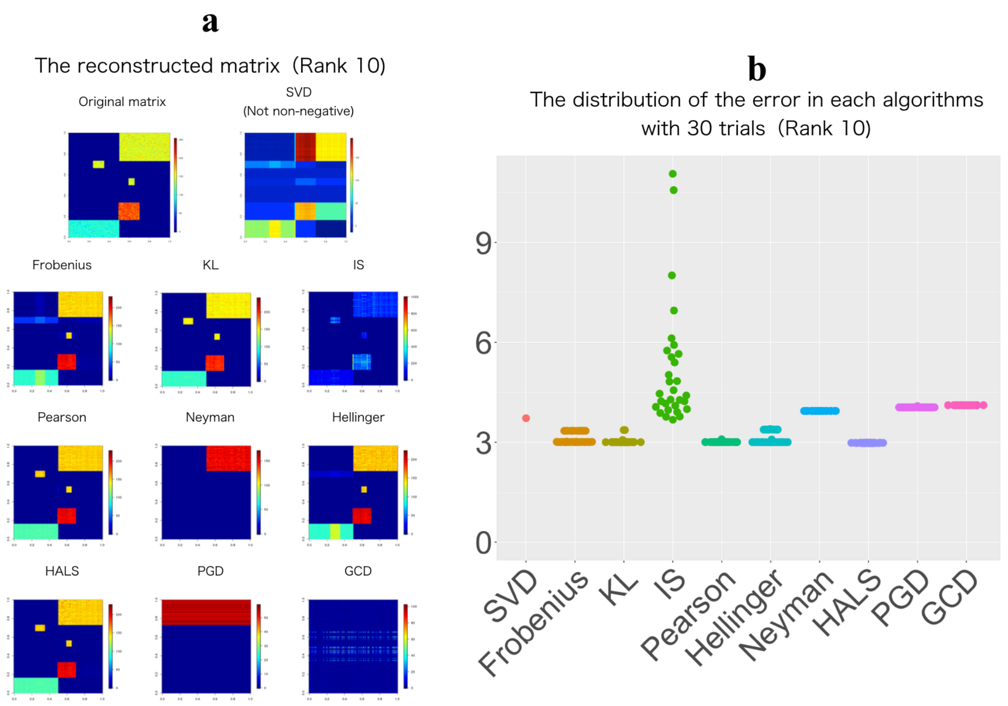

**Supplementary Figure 4 | Convergence of NMF algorithms with toy NMF-type data (Rank 10)**

(a) Reconstructed matrix from the result of each algorithm. (b) Distribution of the reconstruction error of each algorithm based on 30 trials.

Next, we evaluated NTF and NTD in the same way. Using a non-negative CP-type toy model (Figure S1, center), a wide variety of NTF and NTD algorithms were performed 30 times (Figures S5-S7). Even though the data were generated according to CP decomposition, the convergence of NTF was slightly worse than that of NTD. This tendency continued even for larger rank values, such as {5,5,5} and {10,10,10}.

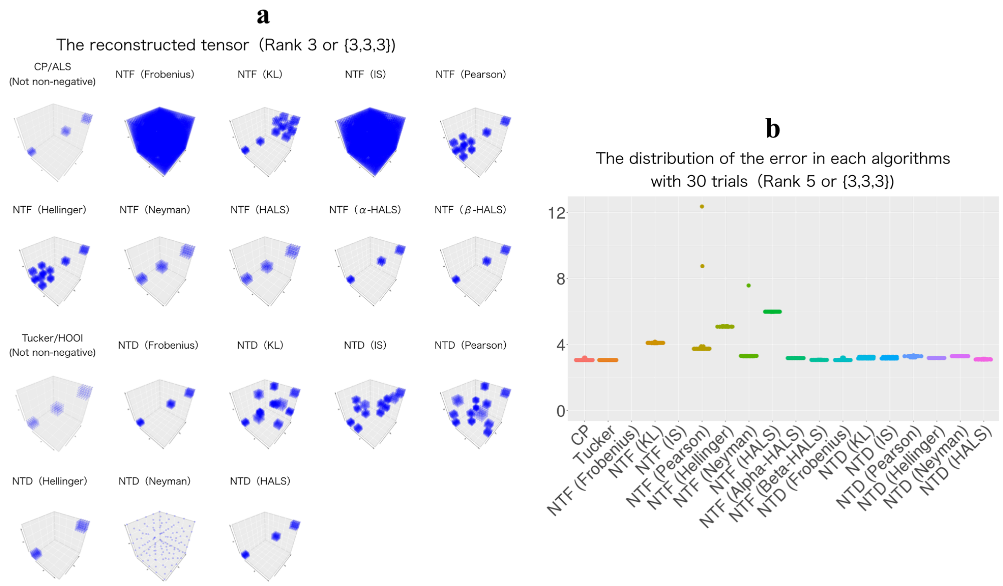

**Supplementary Figure 5 | Convergence of NTF/NTD algorithms with toy non-negative CP-type data (Ranks {3,3,3})**

(a) Reconstructed tensor from the result of each algorithm. (b) Distribution of the reconstruction error of each algorithm based on 30 trials.

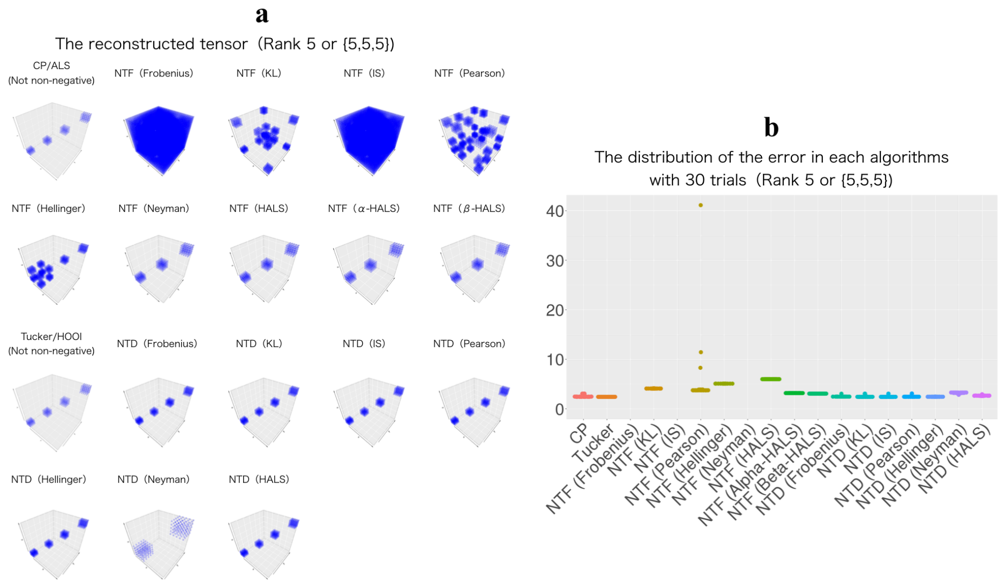

**Supplementary Figure 6 | Convergence of NTF/NTD algorithms with toy non-negative CP-type data (Ranks {5,5,5})**

(a) Reconstructed tensor from the result of each algorithm. (b) Distribution of the reconstruction error of each algorithm based on 30 trials.

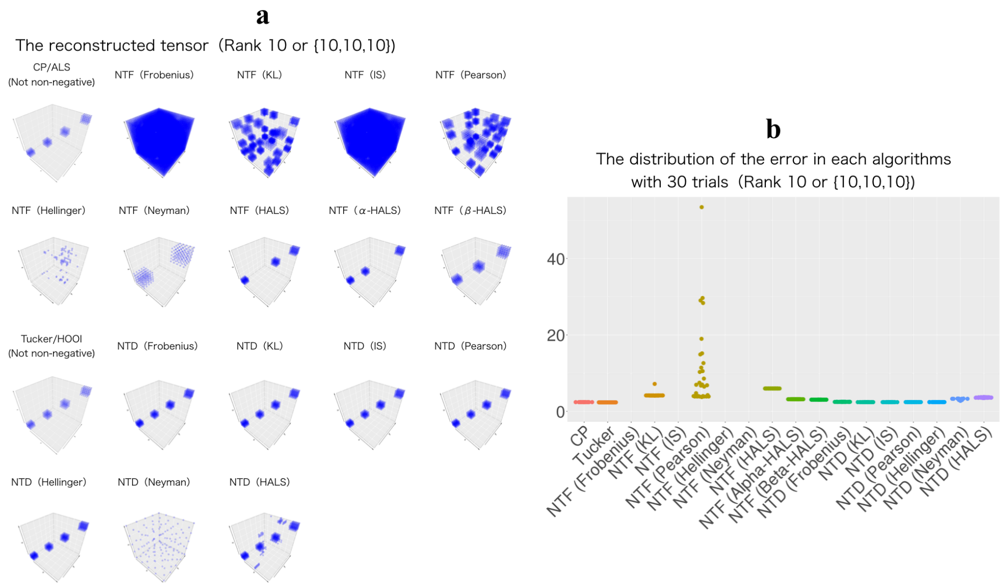

**Supplementary Figure 7 | Convergence of NTF/NTD algorithms with toy non-negative CP-type data (Ranks {10,10,10})**

(a) Reconstructed tensor from the result of each algorithm. (b) Distribution of the reconstruction error of each algorithm based on 30 trials.

Finally, we evaluated NTF and NTD algorithms using a non-negative Tucker-type toy model (Figure S1, right). Here, the performance of NTF was even lower, with some false positive blocks generated and unstable convergence (Figures S8-S10). With appropriate ranks such as {5,5,5} and {10,10,10}, NTD was able to capture such relatively complex block structures, achieving stable convergence was stable with a similar structure detected. For the above reasons, NTD with KL-divergence was introduced as the decomposition method for CCI-tensors in scTensor.

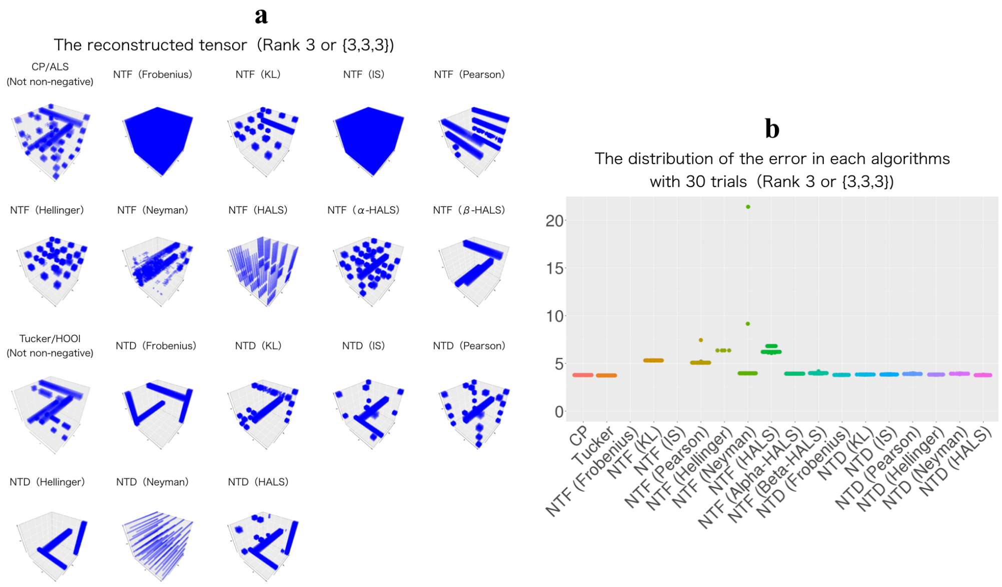

**Supplementary Figure 8 | Convergence of NTF/NTD algorithms with toy non-negative Tucker-type data (Ranks {3,3,3})**

(a) Reconstructed tensor from the result of each algorithm. (b) Distribution of the reconstruction error of each algorithm based on 30 trials.

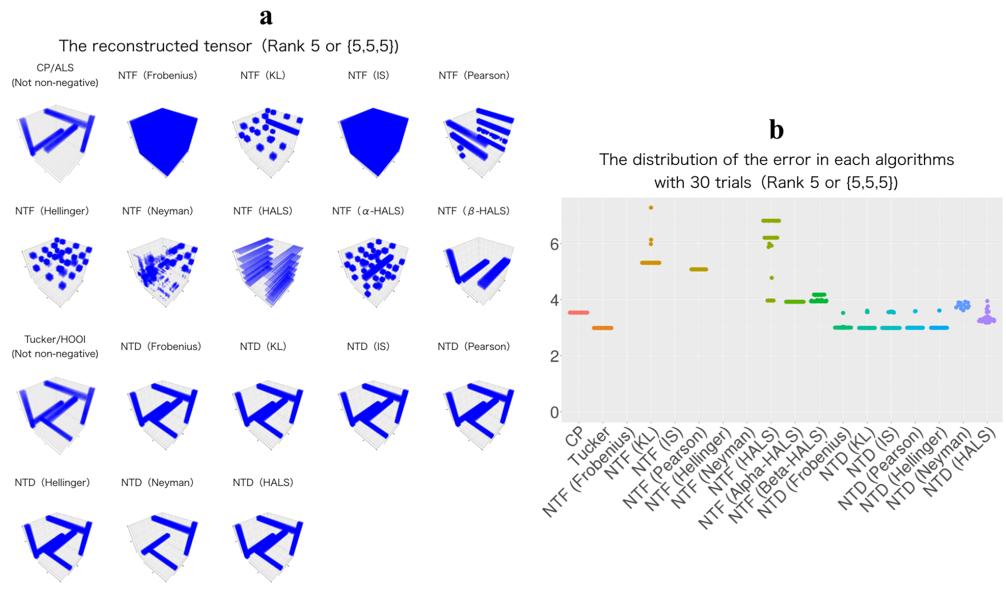

**Supplementary Figure 9 | Convergence of NTF/NTD algorithms with toy non-negative Tucker-type data (Ranks {5,5,5})**

(a) Reconstructed tensor from the result of each algorithm. (b) Distribution of the reconstruction error of each algorithm based on 30 trials.

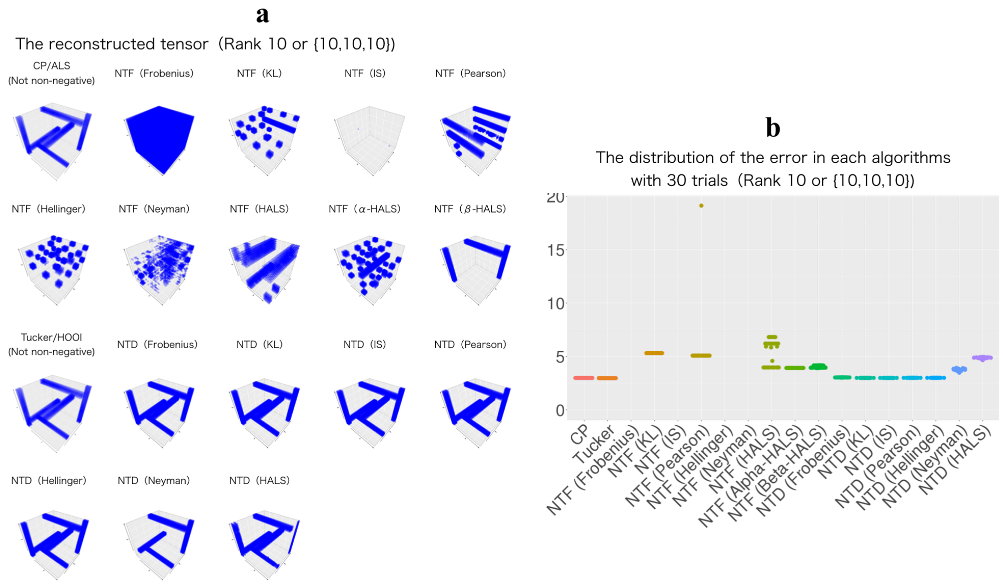

**Supplementary Figure 10 | Convergence of NTF/NTD algorithms with toy non-negative Tucker-type data (Ranks {10,10,10})**
(a) Reconstructed tensor from the result of each algorithm. (b) Distribution of the reconstruction error of each algorithm based on 30 trials.

Although a large number of ranks stabilize the convergence, setting too large of a rank is counter to the motivation of dimensionality reduction in order to project high-dimensional data to lower-dimension data. With large ranks, even the noise signal may be captured as false positives owing to overfitting. Therefore, we introduced a simple rule for estimating appropriate ranks for NTD; we performed SVD with metricized tensors in each mode-$n$ ($n$ = 1, 2, 3) and then retained only the top column vectors explaining 80% to 90% of cumulative variance. The simple rule can be a practical threshold; the rule estimated almost the same ranks (Figure S11, red lines) as the true ranks specified when the toy-model was generated (Figure S11, elbows of plots). The estimated ranks are then used for NTD.

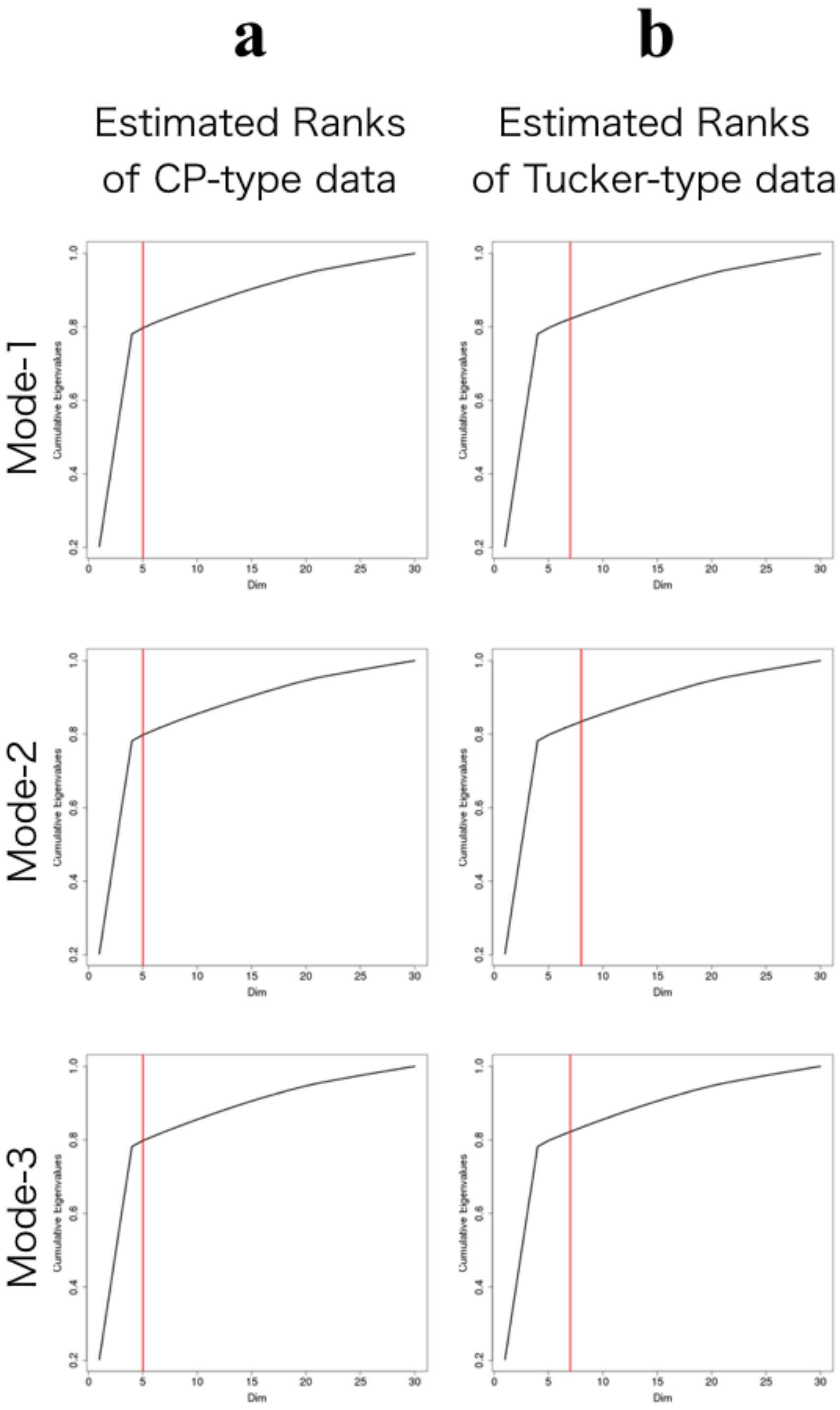

**Supplementary Figure 11 | Estimation of ranks of CP/Tucker-type data**

Estimated ranks for (a) CP-type data and (b) Tucker-type data; black lines represent the cumulative variance calculated by SVD, red lines represent the true rank for generating the datasets.

#### Convergence of NMF, NTD, and NTD with empirical data

Here, we evaluated the convergence of NTD with four empirical datasets: Germline_Female, Germline_Male, Melanoma, and NonMyocyte. These datasets are detailed in the main manuscript, where their analyses are also presented.

As with Figure S11, we estimated the appropriate ranks of four empirical datasets using the cumulative variance of SVD for each matricized tensor (Figure S12), and the ranks of Germline_Female, Germline_Male, Melanoma, and NonMyocyte datasets were estimated as {3,3,16}, {2,3,12}, {3,3,16}, and {2,4,19}, respectively.

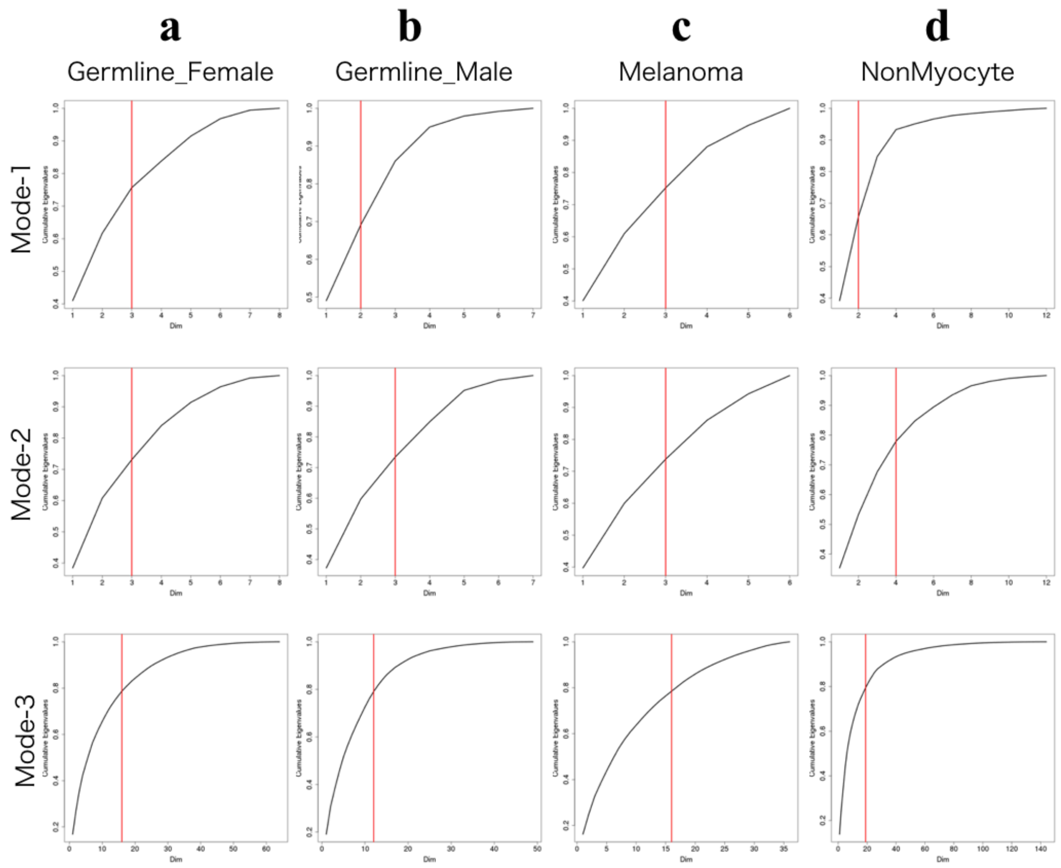

**Supplementary Figure 12 | Estimation of ranks of CP/Tucker-type data**

Estimated ranks for the following datasets: (a) Germline_Female, (b) Germline_Male, (c) Melanoma, and (d) NonMyocyte; black lines represent the cumulative variance calculated by SVD, red lines represent the true rank for generating the datasets.

Next, we evaluated the efficacy of Freeman–Tukey transformation [55] (FTT, $\sqrt{x} + \sqrt{x+1}$), which is a variance-stabilizing transformation of the data matrix, and compared its performance with that of the raw data matrix. The distribution of reconstruction errors shows the results of FTT are slightly better than the results based on raw data (Figure S13-S14).

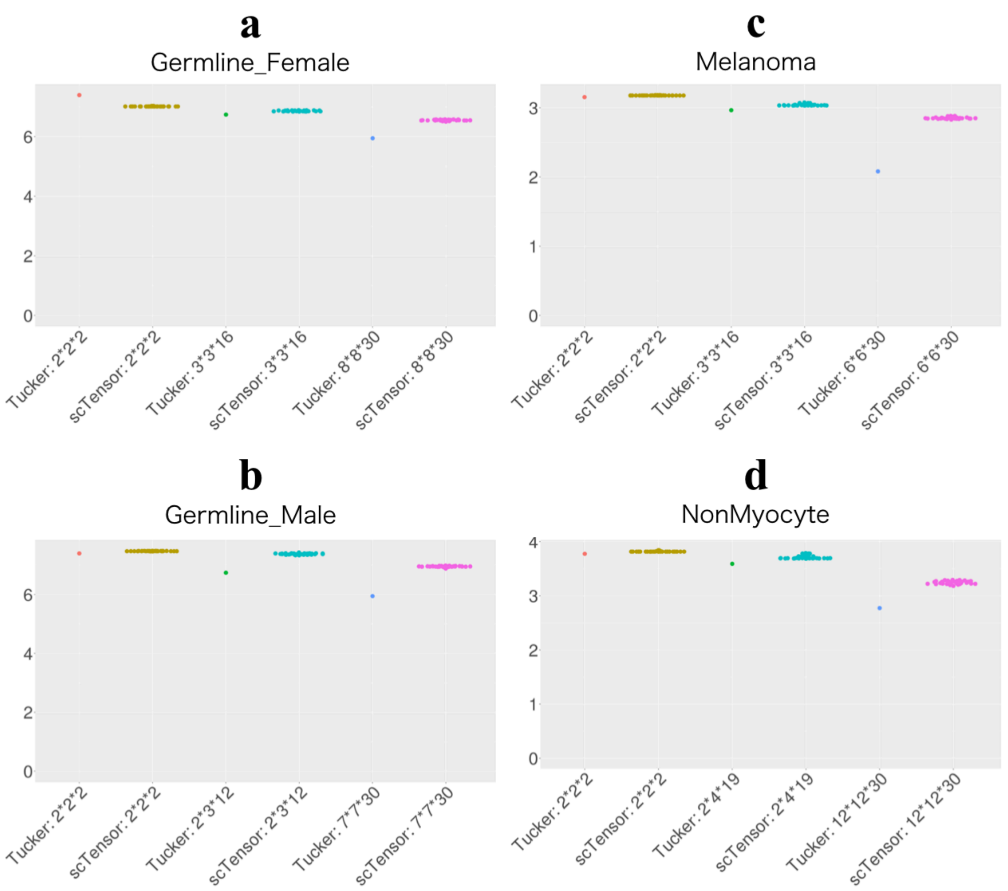

**Supplementary Figure 13 | The distributions of errors for scTensor based on 30 trials performed with four empirical datasets**

Reconstruction errors for the (a) Germline_Female, (b) Germline_Male, (c) Melanoma, and (d) NonMyocyte datasets were calculated. Tucker decomposition with the same ranks was also performed.

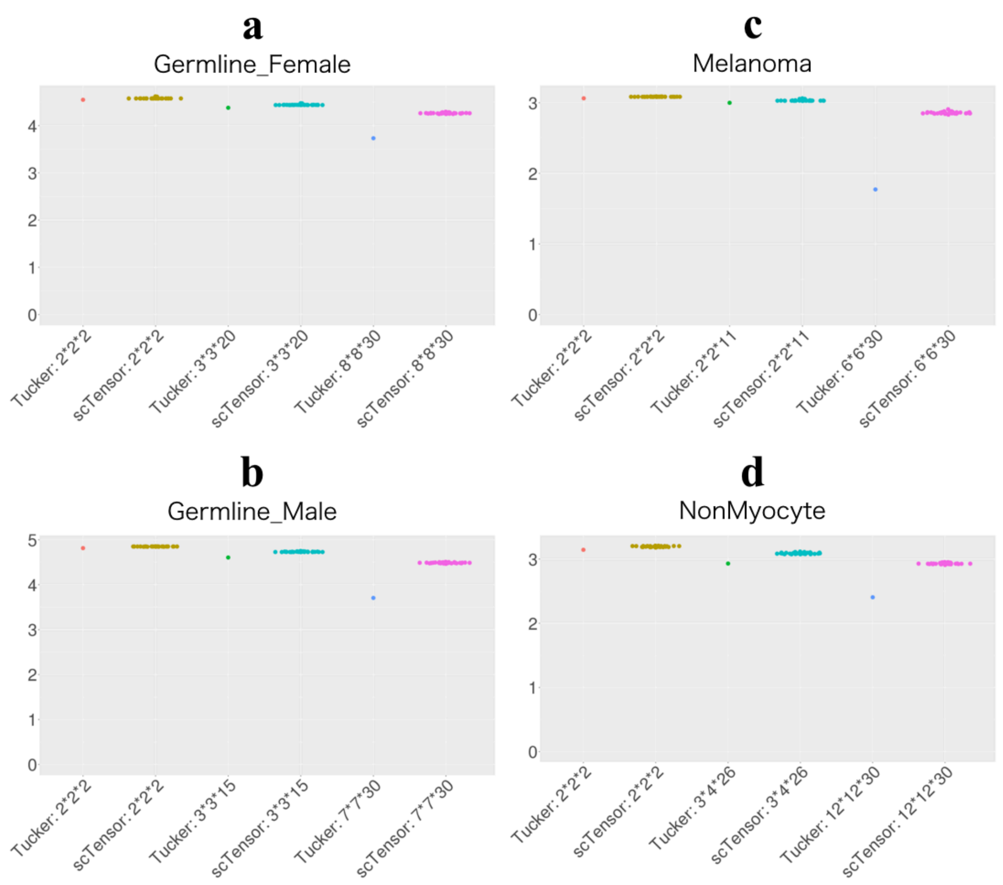

**Supplementary Figure 14 | Distributions of errors for scTensor based on 30 trials with four empirical datasets (FFT)**

Reconstruction errors for (a) Germline_Female, (b) Germline_Male, (c) Melanoma, and (d) NonMyocyte datasets were calculated. Tucker decomposition with the same ranks was also performed.

We also compared the appearance of CCI-strength (cf. main manuscript) for the four empirical datasets (Figure S15-S22). The CCI-strength based on 30 trials shows that the results with minimum error and maximum error are not so different, but the results of FTT are slightly better than the results based on raw data. These evaluations demonstrate that FTT can improve the convergence of NTD. Therefore, in the default mode of scTensor, FTT is applied to the input matrix in advance.

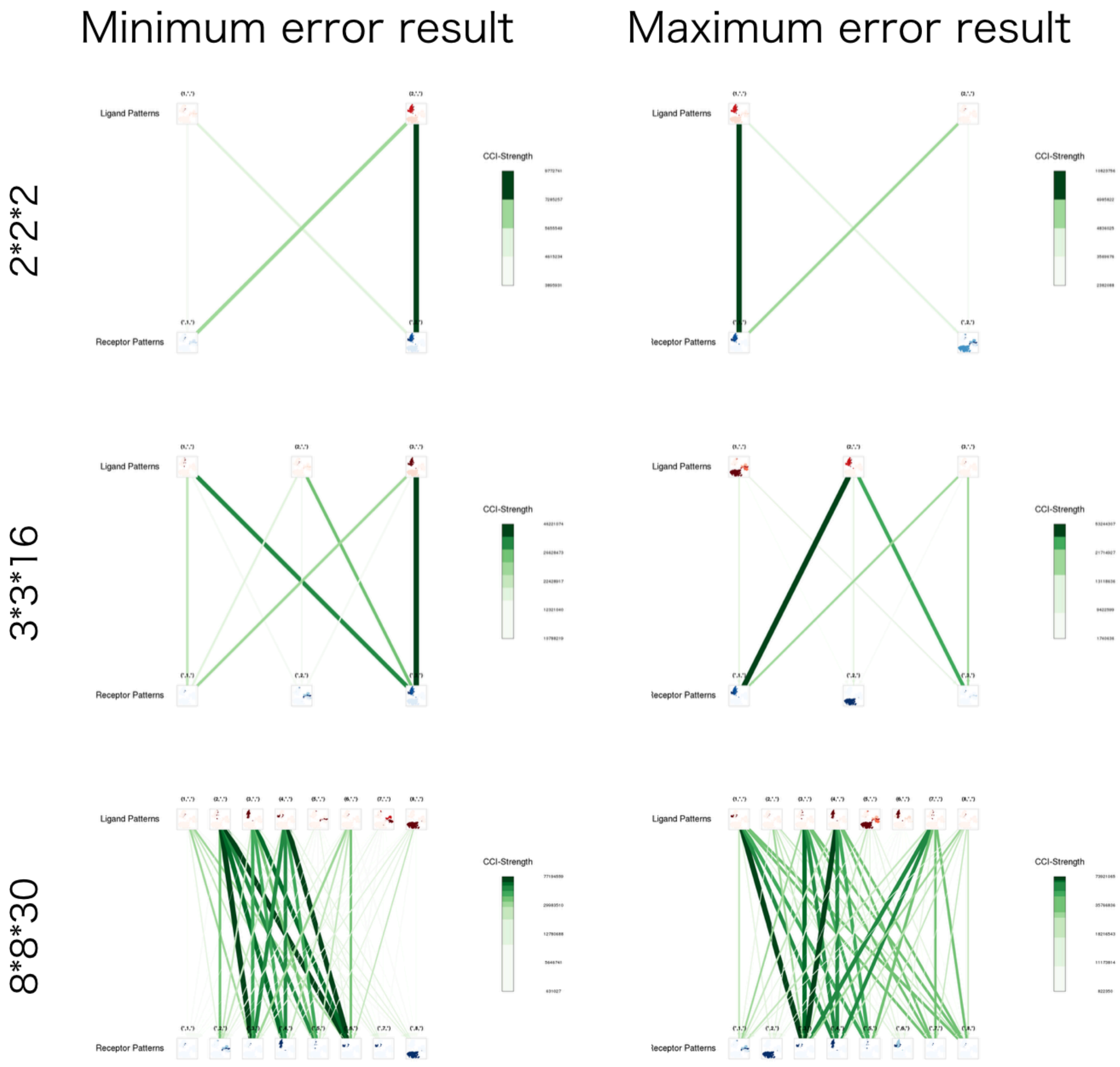

**Supplementary Figure 15 | CCI-strength with different ranks and max/min error results (Germline_Female)**

CCI-strength was calculated based on 30 trials, and only the results with minimum and maximum errors were selected.

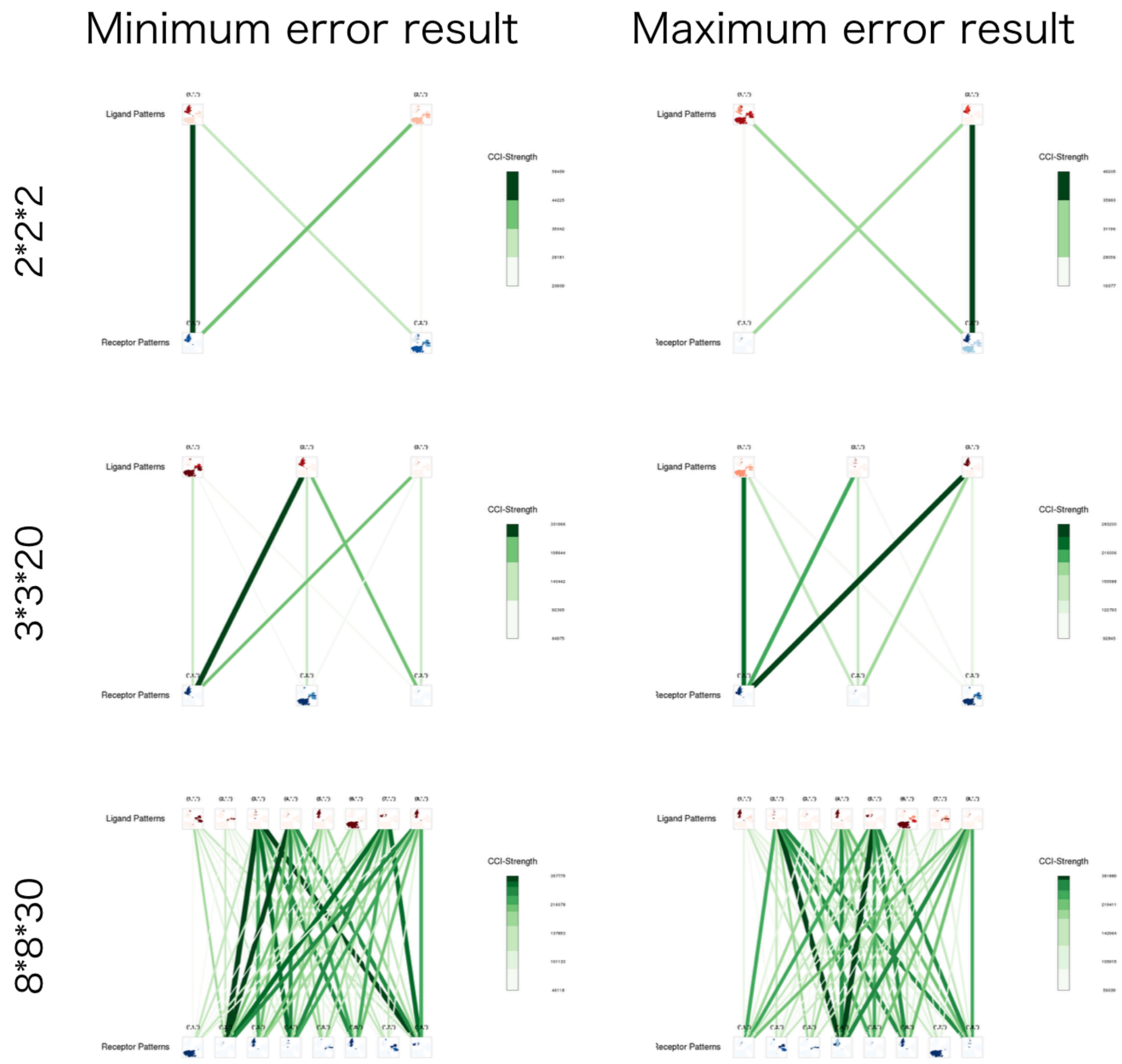

**Supplementary Figure 16 | CCI-strength with different ranks and max/min error results with FTT (Germline_Female)**

CCI-strength was calculated based on 30 trials, and only the results with minimum and maximum errors were selected.

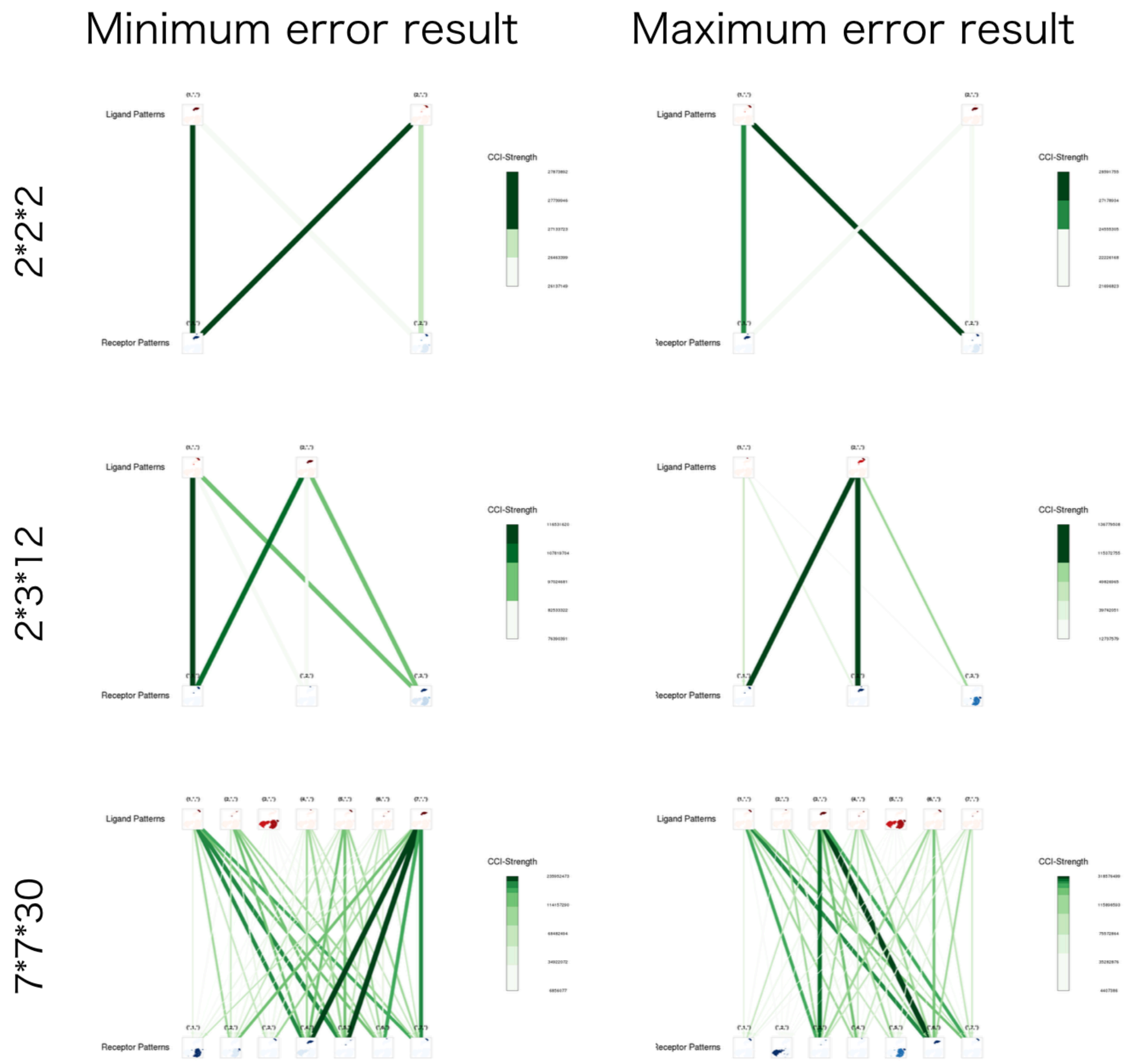

**Supplementary Figure 17 | CCI-strength with different ranks and max/min error results (Germline_Male)**

CCI-strength was calculated based on 30 trials, and only the results with minimum and maximum errors were selected.

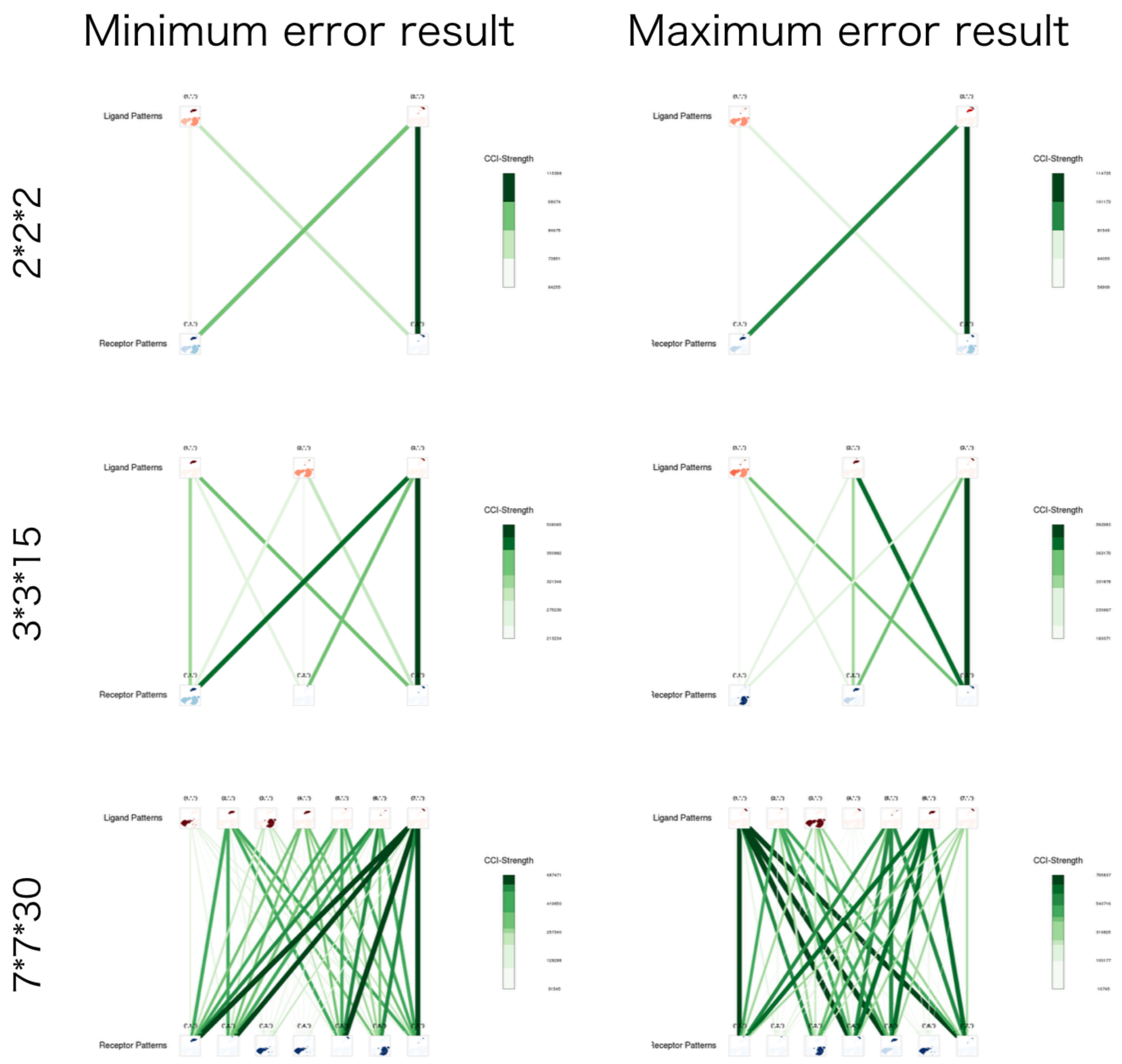

**Supplementary Figure 18 | CCI-strength with different ranks and max/min error results with FTT (Germline_Male)**

CCI-strength was calculated based on 30 trials, and only the results with minimum and maximum errors were selected.

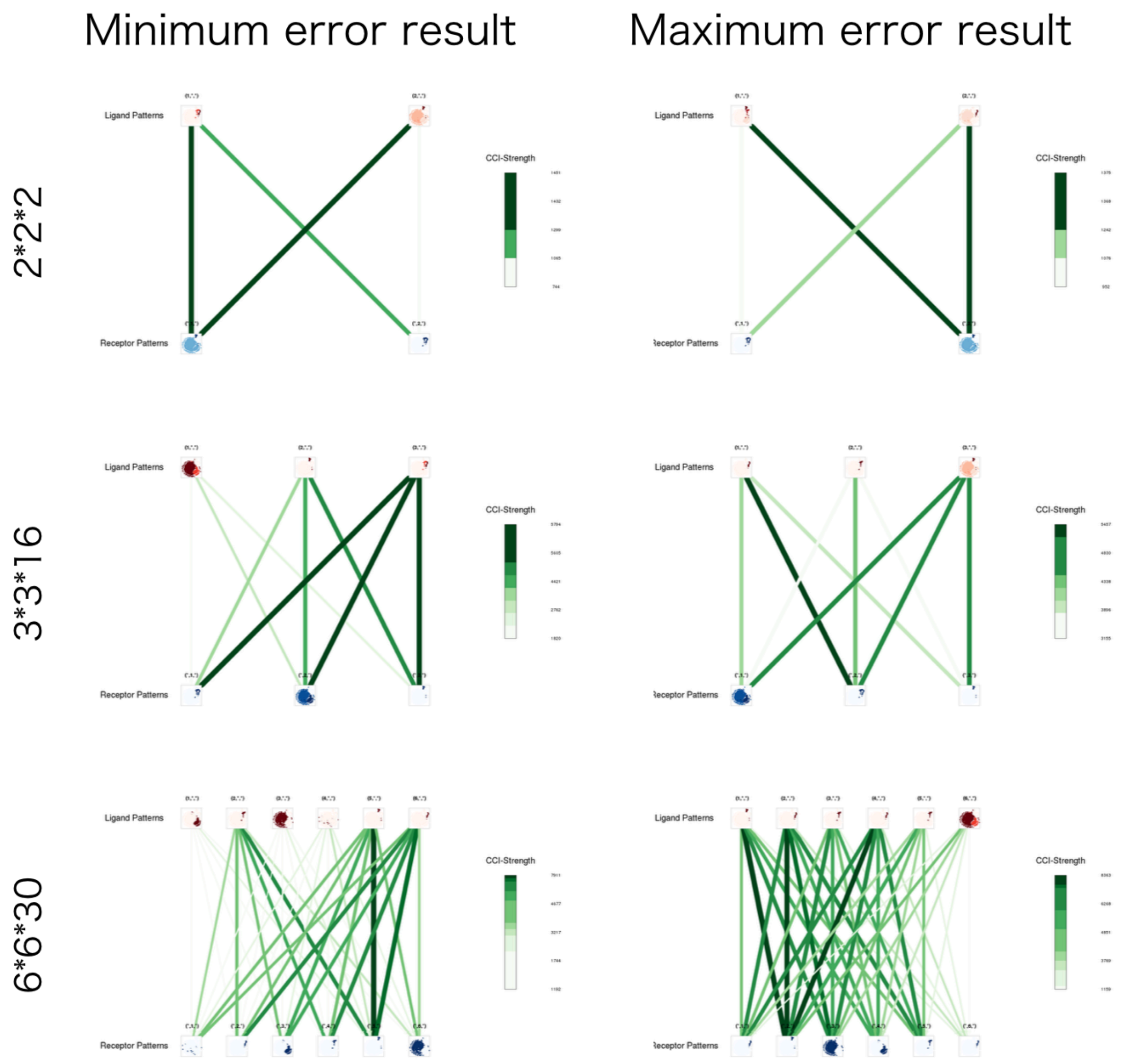

**Supplementary Figure 19 | CCI-strength with different ranks and max/min error results (Melanoma)**

CCI-strength was calculated based on 30 trials, and only the results with minimum and maximum errors were selected.

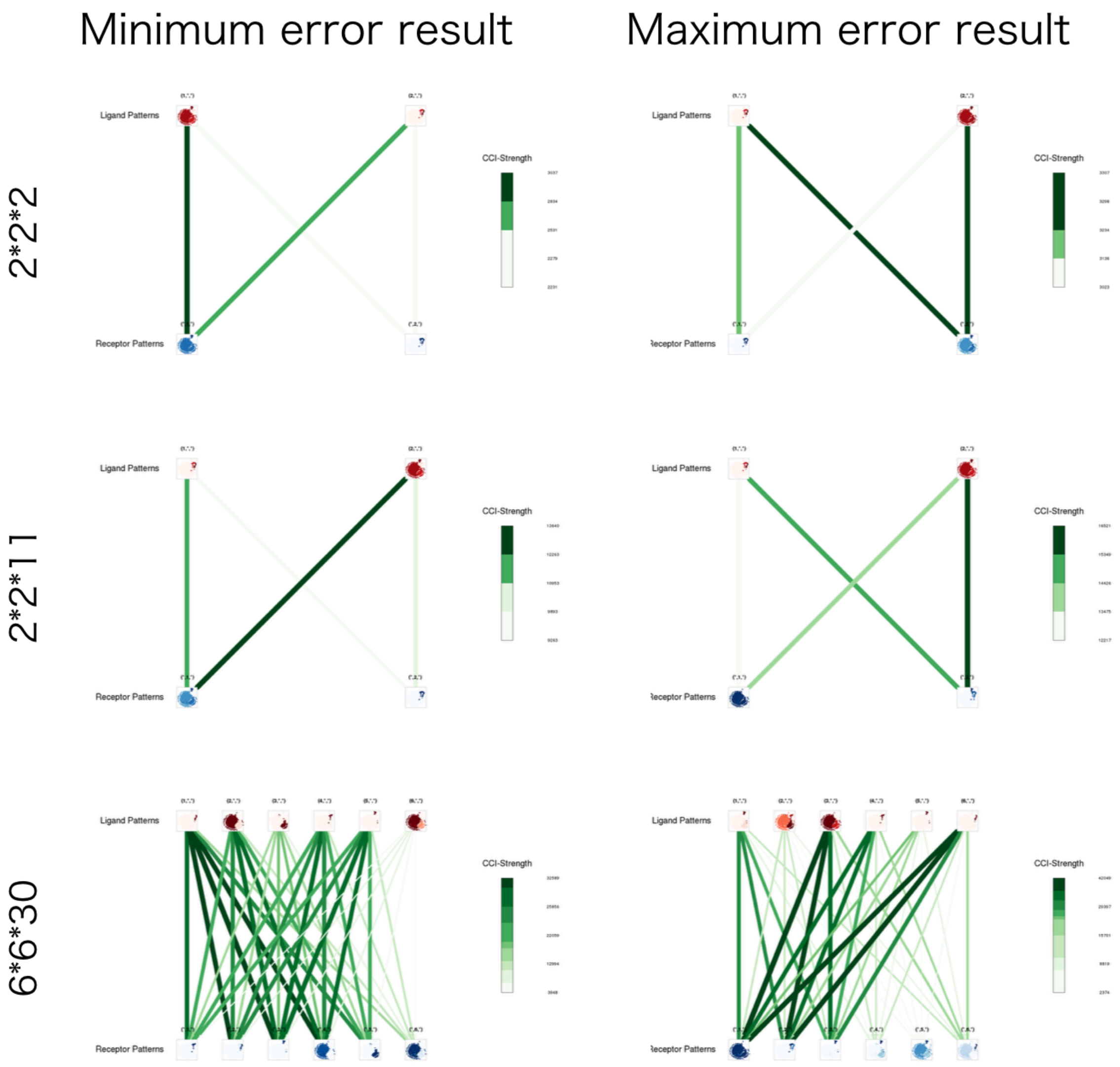

**Supplementary Figure 20 | CCI-strength with different ranks and max/min error results with FTT (Melanoma)**

CCI-strength was calculated based on 30 trials, and only the results with minimum and maximum errors were selected.

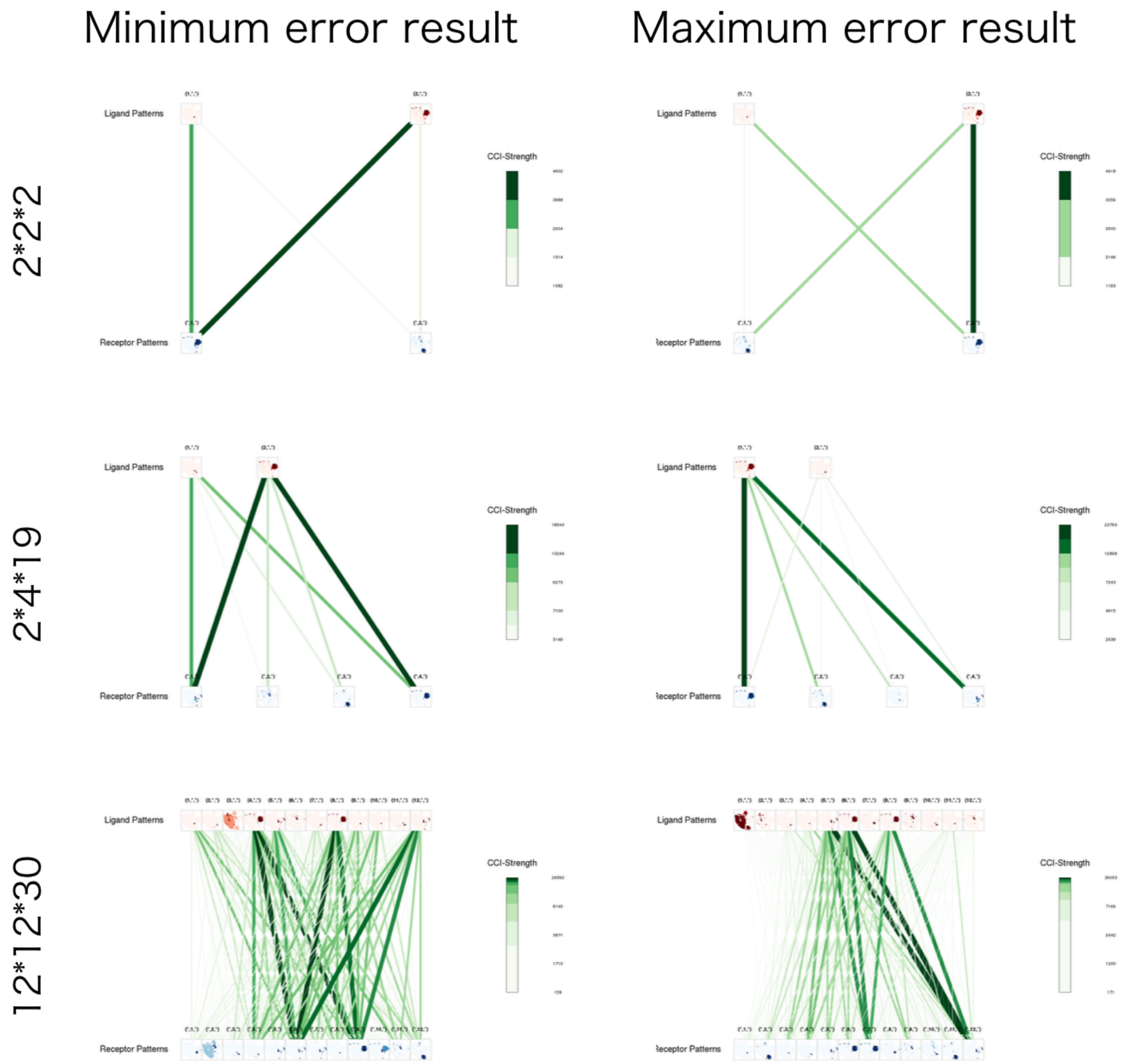

**Supplementary Figure 21 | CCI-strength with different ranks and max/min error results (NonMyocyte)**

CCI-strength was calculated based on 30 trials, and only the results with minimum and maximum errors were selected.

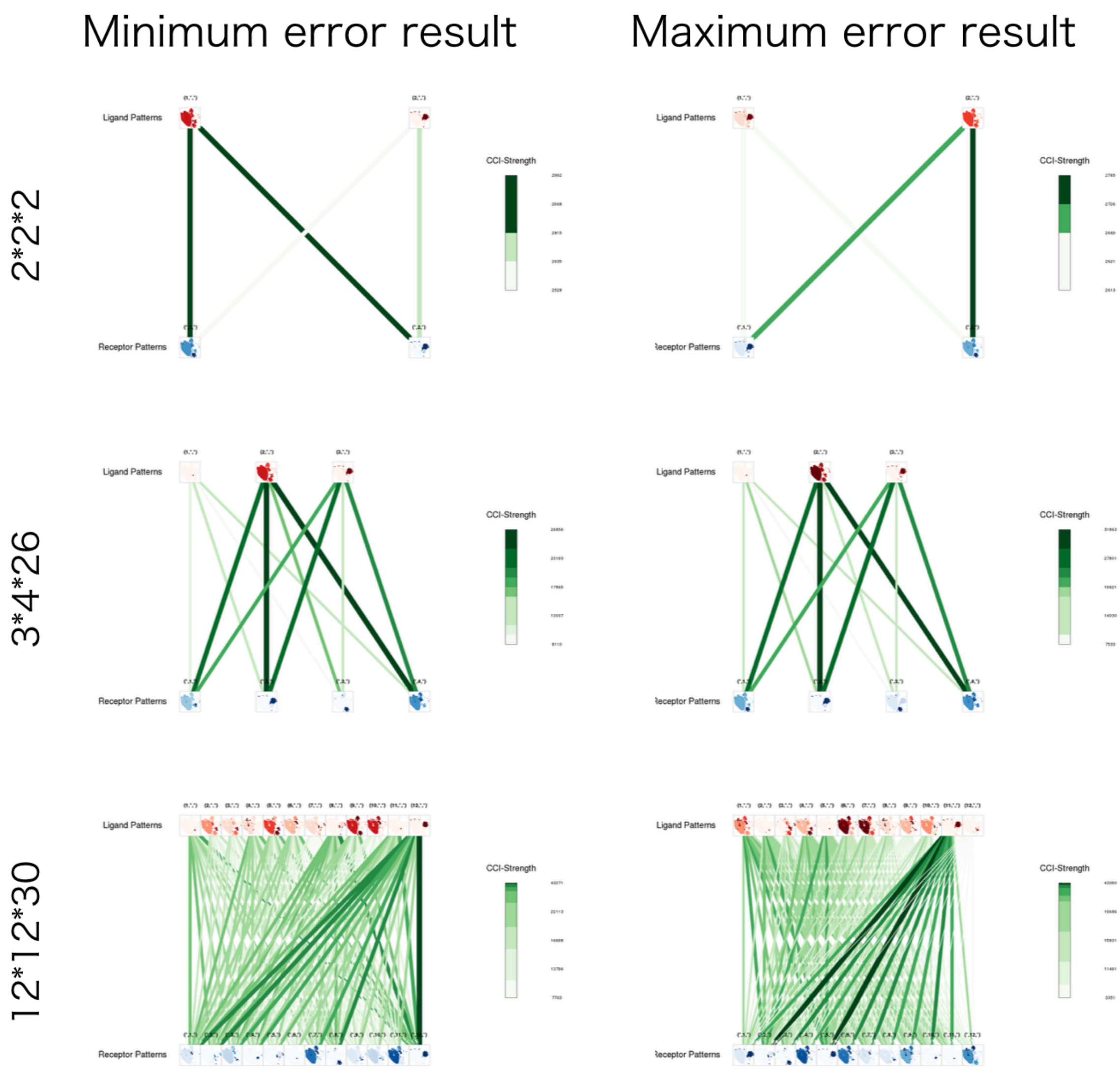

**Supplementary Figure 22 | CCI-strength with different ranks and max/min error results with FTT (NonMyocyte)**

CCI-strength was calculated based on 30 trials, and only the results with minimum and maximum errors were selected.
